# A stochastic model of hippocampal synaptic plasticity with geometrical readout of enzyme dynamics

**DOI:** 10.1101/2021.03.30.437703

**Authors:** Yuri Elias Rodrigues, Cezar Tigaret, Hélène Marie, Cian O’Donnell, Romain Veltz

**Affiliations:** Université Côte d’Azur, Nice, Alpes-Maritimes, France; Institut national de recherche en informatique et en automatique (INRIA), Sophia Antipolis, France; Institut de Pharmacologie Moléculaire et Cellulaire (IPMC), Valbonne, France; Neuroscience and Mental Health Research Institute, Division of Psychological Medicine and Clinical Neurosciences, School of Medicine, Cardiff University, Cardiff, UK; Computational Neuroscience Unit, School of Computer Science, Electrical and Electronic Engineering, and Engineering Mathematics, University of Bristol, Bristol, UK

**Author notes:** **For correspondence:** (RV). Co-senior authors (Hélène Marie, Cian O’Donnell, Romain Veltz). Life Sciences School, University of Sussex, UK. School of Computing, Engineering and Intelligent Systems, Ulster University, UK.

## Abstract

Discovering the rules of synaptic plasticity is an important step for understanding brain learning. Existing plasticity models are either 1) top-down and interpretable, but not flexible enough to account for experimental data, or 2) bottom-up and biologically realistic, but too intricate to interpret and hard to fit to data. To avoid the shortcomings of these approaches, we present a new plasticity rule based on a geometrical readout mechanism that flexibly maps synaptic enzyme dynamics to predict plasticity outcomes. We apply this readout to a multi-timescale model of hippocampal synaptic plasticity induction that includes electrical dynamics, calcium, CaMKII and calcineurin, and accurate representation of intrinsic noise sources. Using a single set of model parameters, we demonstrate the robustness of this plasticity rule by reproducing nine published *ex vivo* experiments covering various spike-timing and frequency-dependent plasticity induction protocols, animal ages, and experimental conditions. Our model also predicts that *in vivo*-like spike timing irregularity strongly shapes plasticity outcome. This geometrical readout modelling approach can be readily applied to other excitatory or inhibitory synapses to discover their synaptic plasticity rules.

## Introduction

To understand how brains learn, we need to identify the rules governing how synapses change their strength in neural circuits. What determines whether each synapse strengthens, weakens, or stays the same? The dominant principle at the basis of current models of synaptic plasticity is the Hebb postulate (***Hebb, 1949***) which states that neurons with correlated electrical activity strengthen their synaptic connections, while neurons active at different times weaken their connections. In particular, spike-timing-dependent plasticity (STDP) models (***Blum and Abbott, 1996; Gerstner et al., 1996; Eurich et al., 1999***) were formulated based on experimental observations that precise timing of pre- and post-synaptic spiking determines whether synapses are strengthened or weakened (***Debanne et al., 1996; Tsodyks and Markram, 1997; Bi and Poo, 1998; Markram et al., 2011***). However, experiments also found that plasticity induction depends on the rate and number of stimuli delivered to the synapse (***Dudek and Bear, 1992; Sjöström et al., 2001***), and the level of dendritic spine depolarisation (***Artola et al., 1990; Magee and Johnston, 1997; Sjöström and Häusser, 2006; Golding et al., 2002; Hardie and Spruston, 2009***). The lack of satisfactory plasticity models based solely on neural spiking prompted researchers to consider simple models based on synapse biochemistry (***Castellani et al., 2001, 2005***). Following a proposed role for postsynaptic calcium (Ca^2+^) signalling in synaptic plasticity (***Lisman, 1989***), previous models assumed that the amplitude of postsynaptic calcium controls long-term alterations in synaptic strength, with moderate levels of calcium causing long-term depression (LTD) and high calcium causing long-term potentiation (LTP) (***Shouval et al., 2002; Karmarkar and Buonomano, 2002***). However experimental data suggests that calcium dynamics are also important (***Yang et al., 1999; Mizuno et al., 2001; Wang et al., 2005; Nevian and Sakmann, 2006; Tigaret et al., 2016***). As a result, subsequent phenomenological models of plasticity incorporated slow variables that integrate the fast synaptic input signals, loosely modelling calcium and its downstream effectors (***Abarbanel et al., 2003; Rubin et al., 2005; Rackham et al., 2010; Clopath and Gerstner, 2010; Kumar and Mehta, 2011; Graupner and Brunel, 2012; Honda et al., 2013; Standage et al., 2014; De Pittà and Brunel, 2016***). Concurrently, more detailed models tried to explicitly describe the molecular pathways integrating the calcium dynamics and its stochastic nature (***Cai et al., 2007; Shouval and Kalantzis, 2005; Miller et al., 2005; Zeng and Holmes, 2010; Yeung et al., 2004***). However, even these models do not account for data showing that plasticity is highly sensitive to physiological conditions such as the developmental age of the animal (***Dudek and Bear, 1993; Meredith et al., 2003; Cao and Harris, 2012; Cizeron et al., 2020***), extracellular calcium and magnesium concentrations (***Mulkey and Malenka, 1992; Inglebert et al., 2020***) and tissue temperature (***Volgushev et al., 2004; Wittenberg and Wang, 2006; Klyachko and Stevens, 2006***). The fundamental issue is that the components of these phenomenological models do not directly map to biological components of synapses, so they cannot automatically model alterations due to physiological and experimental conditions. This absence limits the predictive power of this class of plasticity models.

An alternative approach taken by several groups (***Bhalla and Iyengar, 1999; Jędrzejewska-Szmek et al., 2017; Blackwell et al., 2019; Chindemi et al., 2020; Zhang et al., 2021***) was to model the complex molecular cascade leading to synaptic weight changes. The main benefit of this approach is the direct correspondence between the model’s components and biological elements, but this comes at the price of a large number of poorly constrained parameters. Additionally, the increased number of nonlinear equations and stochasticity makes fitting to plasticity experiment data difficult (***Mäki-Marttunen et al., 2020***). Subtle differences between experimental STDP protocols can produce completely different synaptic plasticity outcomes, indicative of finely tuned synaptic behaviour. This raises major challenges for both simple and complex models.

To tackle this problem, we devised a new plasticity rule based on a bottom-up, data-driven approach by building a biologically-grounded model of plasticity induction at a single rat hippocampal CA3–CA1 synapse. We focused on this synapse type because of the abundant published experimental data that can be used to quantitatively constrain the model parameters. Compared to previous models in the literature, we aimed for an intermediate level of detail: enough biophysical components to capture the key dynamical processes underlying plasticity induction, but not the detailed molecular cascade underlying plasticity expression; much of which is poorly quantified for the various experimental conditions we cover in this study.

Our model is centred on dendritic spine electrical dynamics, calcium signalling and immediate downstream molecules, which we then map to synaptic strength change via a conceptually new dynamical, geometric readout mechanism. Crucially, the model also captured intrinsic noise based on the stochastic switching of synaptic receptors and ion channels (***Yuste et al., 1999; Ribrault et al., 2011***). We found that, with a single set of parameters, the model can account for published data from spike-timing and frequency-dependent plasticity experiments, and variations in physiological parameters influencing plasticity outcomes. We also tested how the model responded to *in vivo-* like spike timing jitter and spike failures, and found that the plasticity rules were highly sensitive to these subtle input alterations.

## Results

### A multi-timescale model of synaptic plasticity induction

We built a computational model of plasticity induction ata single CA3-CA1 rat glutamatergic synapse (***Figure 1***). Our goal was to reproduce results on synaptic plasticity that explored the effects of several experimental parameters: fine timing differences between pre and postsynaptic spiking (***Figure 2*** and ***Figure 3***); stimulation frequency (***Figure 4***); animal age (***Figure 5***); external calcium and magnesium (***Figure 6***); stochasticity in the firing structure (***Figure 7***), temperature and experimental conditions variations (***Supplemental files***). Where possible, we set parameters to values previously estimated from synaptic physiology and biochemistry experiments, and tuned the remainder within physiologically plausible ranges to reproduce ourtarget plasticity experiments (see ***Methods and Materials***).

**Figure 1.**
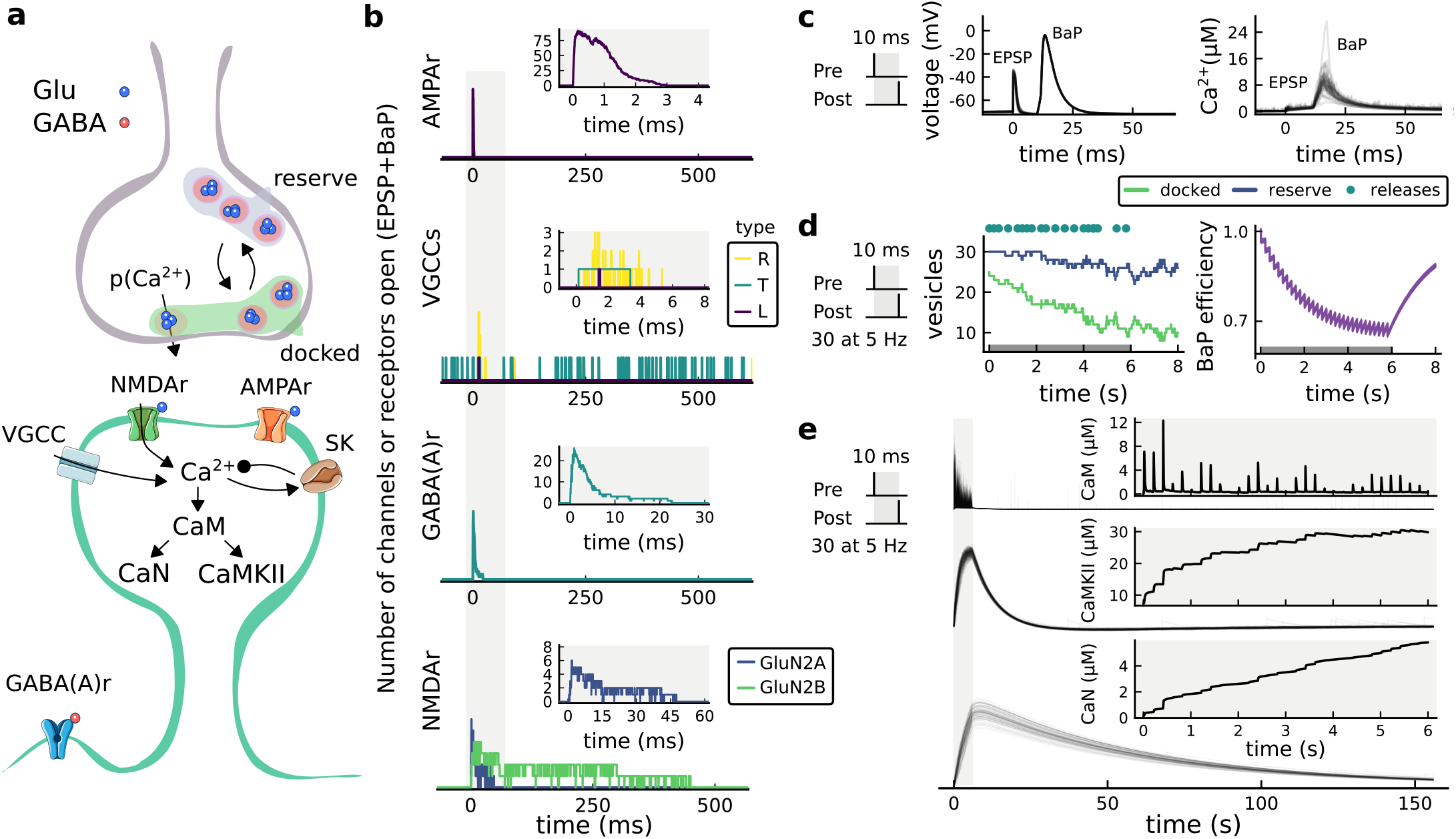
The synapse model, its timescales and mechanisms. **a**, Model diagram with the synaptic components including pre and postsynaptic compartments and inhibitory transmission (bottom left). **b**, Stochastic dynamics of the different ligand-gated and voltage-gated ion channels in the model. Plots show the total number of open channels as a function of time. AMPAr, NMDAr: AMPA- and NMDA-type glutamate receptors respectively; GABA(A)r: Type A GABA receptors; VGCC: R-, T- and L-type voltage-gated Ca2+ channels; SK: SK potassium channels. The insets show a zoomed time axis highlighting the difference in timescale of the activity among the channels. **c**, Dendritic spine membrane potential (left) and calcium concentration (right) as function of time for a single causal (1Pre1Post10) stimulus (EPSP: single excitatory postsynaptic potential, “1Pre”; BaP: single back-propagated action potential, “1Post”). **d**, Left: depletion of vesicle pools (reserve and docked) induced by 30 pairing repetitions delivered at 5 Hz (***Sterratt et al., 2011***), see ***Methods and Materials***. The same depletion rule is applied to both glutamate- and GABA-containing vesicles. Right: BaP efficiency as function of time. BaP efficiency phenomenologically captures the distance-dependent attenuation of BaP (***Buchanan and Mellor, 2007; Golding et al., 2001***), see ***Methods and Materials***. **e**, Concentration of active enzyme for CaM, CaN and CaMKII, as function of time triggered by 30 repetitions of 1Pre1Post10 pairing stimulations delivered at 5 Hz. The vertical grey bar is the duration of the stimuli, 6 s. The multiple traces in the graphs in panels **c** (right) and **e** reflect the run-to-run variabiltity due to the inherent stochasticity in the model.

**Figure 2.**
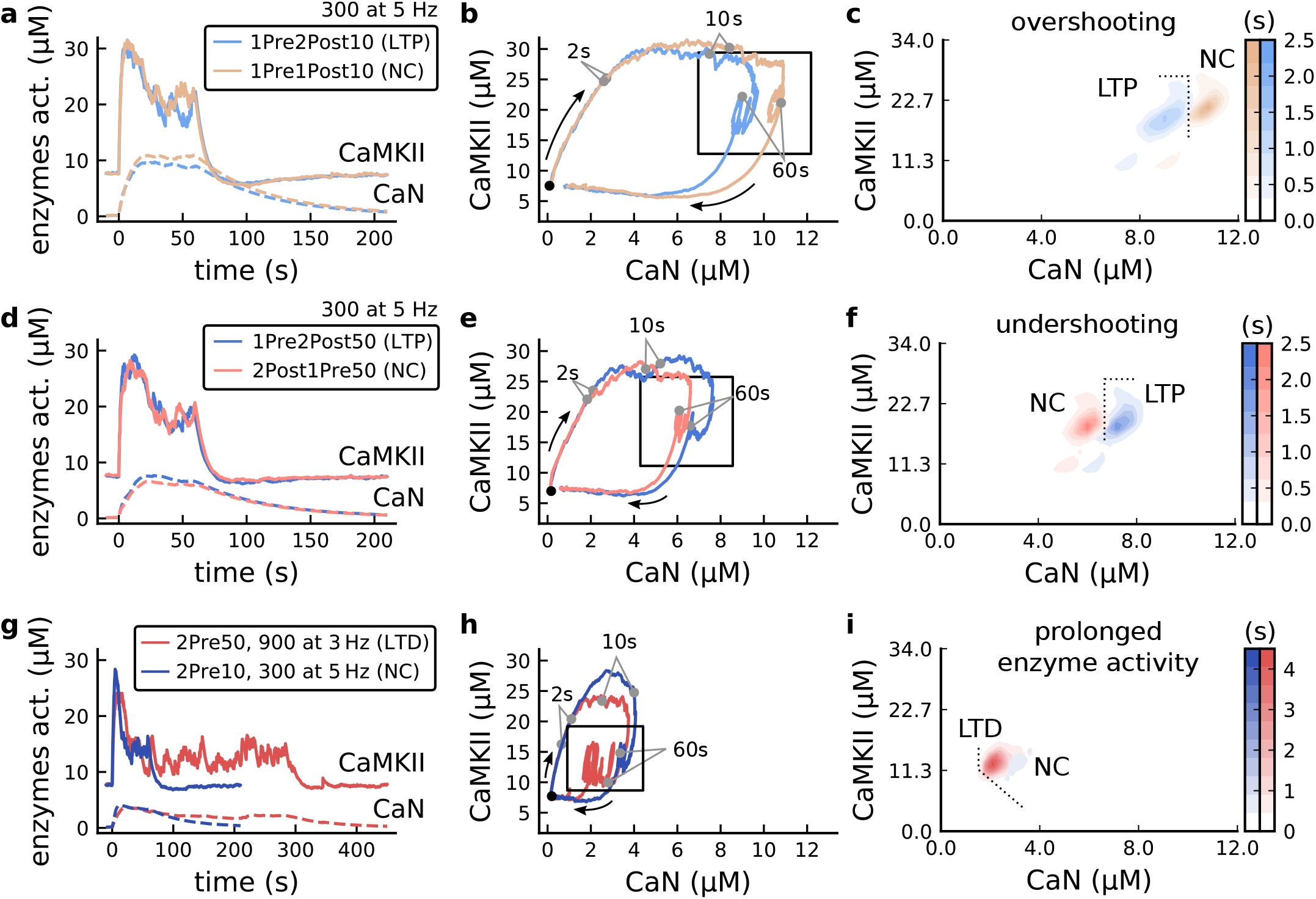
The duration and amplitude of the joint CaN-CaMKII activity differentiates plasticity protocols. **a**, Time-course of active enzyme concentration for CaMKII (solid line) and CaN (dashed line) triggered by two protocols consisting of 300 repetitions at 5 Hz of 1Pre2Post10 or 1Pre1Post10 stimulus pairings. Protocols start at time 0 s. Experimental data indicates that 1Pre2Post10 and 1Pre1Post10 produce LTP and no change (NC), respectively. **b**, Trajectories of joint enzymatic activity (CaN-CaMKII) as function of time for the protocols in panel **a**, starting at the initial resting state (filled black circle). The arrows show the direction of the trajectory and filled grey circles indicate the time points at 2,10 and 60 s after the beginning of the protocol represented as 2,10 and 60 s. The region of the CaN-CaMKII plane enclosed in the black square is expanded in panel **c**. **c**, Mean-time (colorbar) spent by the orbits in the CaN-CaMKII plane region expanded from panel **b** for each protocol (average of 100 samples). For panels **c, f** and **i** the heat maps were based on enzyme activity throughout the protocol plus a further 10 s after the stimulation ended. **d-f**, CaN-CaMKII activities for the protocols 1Pre2Post50 (LTP-inducing) and 2Post1Pre50 (NC) depicted in the same manner as in panels **a-c**. **g-i**, CaN-CaMKII activities for the LTD-inducing protocol 2Pre50 (900 repetitions at 3 Hz) and the NC protocol 2Pre10 (300 repetitions at 5 Hz) depicted in the same manner as in panels **a-c**.

**Figure 3.**
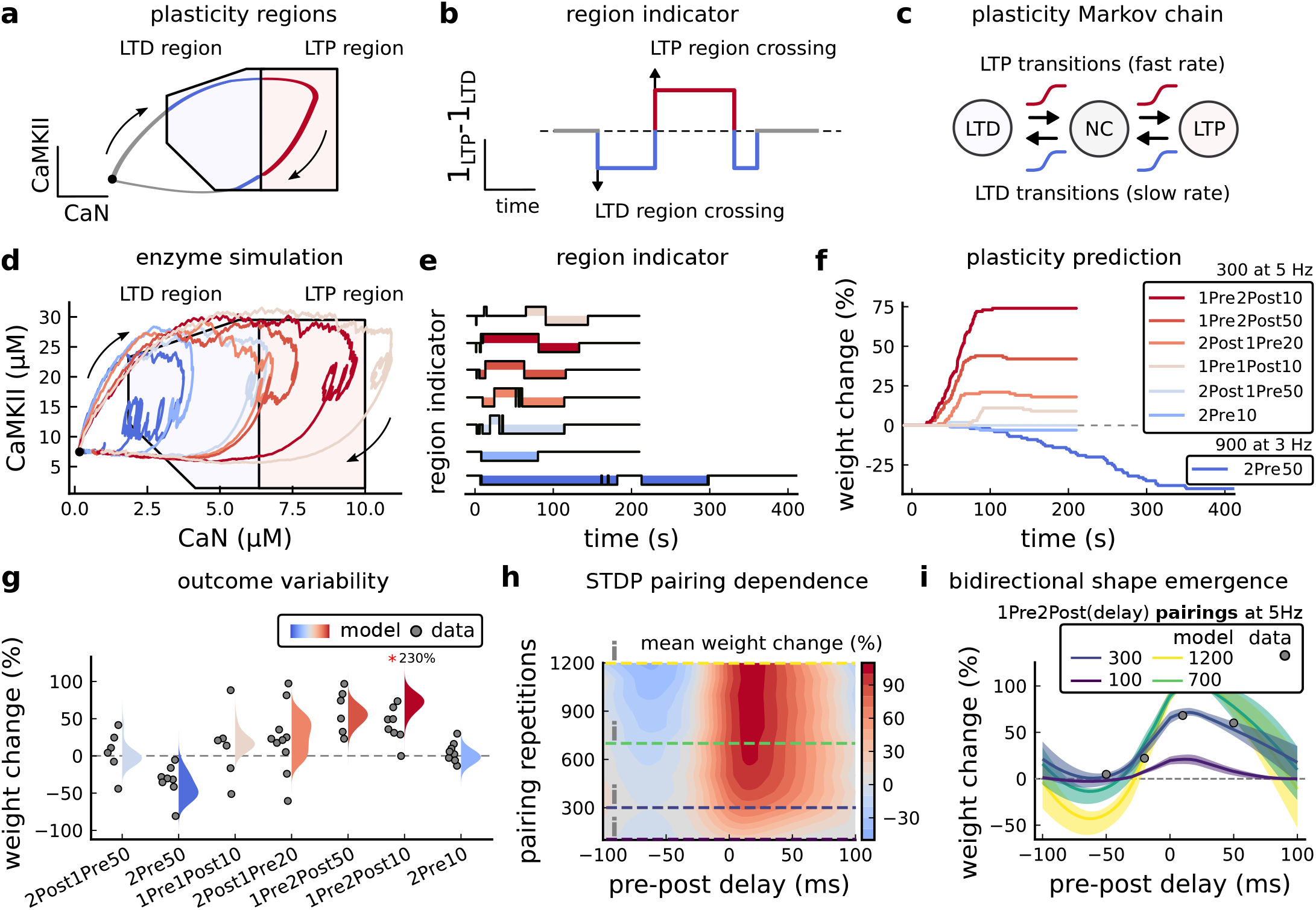
Read-out strategy to accurately model *Tigaret et al. (2016)* experiment. **a**, Illustration of the joint CaMKII and CaN activities crossing the plasticity regions. Arrows indicate the flow of time, starting at the filled black circle. Time is hidden so that changes in active enzyme concentrations are seen more clearly. **b**, Region indicator showing when the joint CaN and CaMKII activity crosses the LTD or LTP regions in panel **a**. For example, the LTP indicator is such that 1_*LTP*_(*x*) = 1 if *x* ∈ *LTP* and 0 otherwise. Leaving the region activates a leaking mechanism that keeps track of the accumulated time inside the region. Such leaking mechanism drives the transition rates used to predict plasticity (*Methods and Materials*). **c**, Plasticity Markov chain with three states: LTD, LTP and NC. There are only two transition rates which are functions of the plasticity region indicator (***Methods and Materials***). The LTP transition is fast whereas the LTD transition is slow, meaning that LTD change requires longer time inside the LTD region (panel **a**). The NC state starts with 100 processes. **d**, Joint CaMKII and CaN activity for all protocols in *Tigaret et al. (2016)* (shown in panel **f**). The stimulus ends when the trajectory becomes smooth. Trajectories correspond to those in ***Figure 2b,e and h***, at 60 s. **e**, Region indicator for the protocols in panel **f**. The upper square bumps are caused by the protocol crossing the LTP region, the lower square bumps when the protocol crosses the LTD region (as in panel **d**). **f**, Synaptic weight (%) as function of time for each protocol. The weight change is defined as the number (out of 100) of states in the LTP state minus the number of states in the LTD state (panel **c**). The trajectories correspond to the median of the simulations in panel **g**. **g**, Synaptic weight change (%) predicted by the model compared to data (EPSC amplitudes) from *Tigaret et al. (2016)* (100 samples for each protocol, also for panel **h** and **i**). The data (filled grey circles) was provided by ***Tigaret et al. (2016)*** (note an 230% outlier as the red asterisk). **h**, Predicted mean synaptic weight change (%) as a function of delay (ms) and number of pairing repetitions (pulses) for the protocol 1Pre2Post(delay), where delays are between −100 and 100 ms. LTD is induced by 2Post1Pre50 after at least 500 pulses. The mean weight change along each dashed line is reported in the STDP curves in panel **i**. **i**, Synaptic weight change (%) as a function of pre-post delay. Each plot corresponds to a different pairing repetition number (color legend). The solid line shows the mean, and the ribbons are the 2nd and 4th quantiles. The filled grey circles are the data means estimated in ***Tigaret et al. (2016)***, also shown in panel **g**. **Figure 3-Figure supplement 1.** Standard models comparison for predicting plasticity fail to account for the data from ***Tigaret et al. (2016)***. **Figure 3-Figure supplement 2.** Comparison showing different roles of stochasticity in the model. **Figure 3-Figure supplement 3.** Effects of blocking VGCCs. **Figure 3-Figure supplement 4.** Exclusively setting vertical boundaries (no CaMKII selectivity) fails to capture the correct outcome. **Figure 3-Figure supplement 5.** Varying *Tigaret et al. (2016)* experimental parameters.

**Figure 4.**
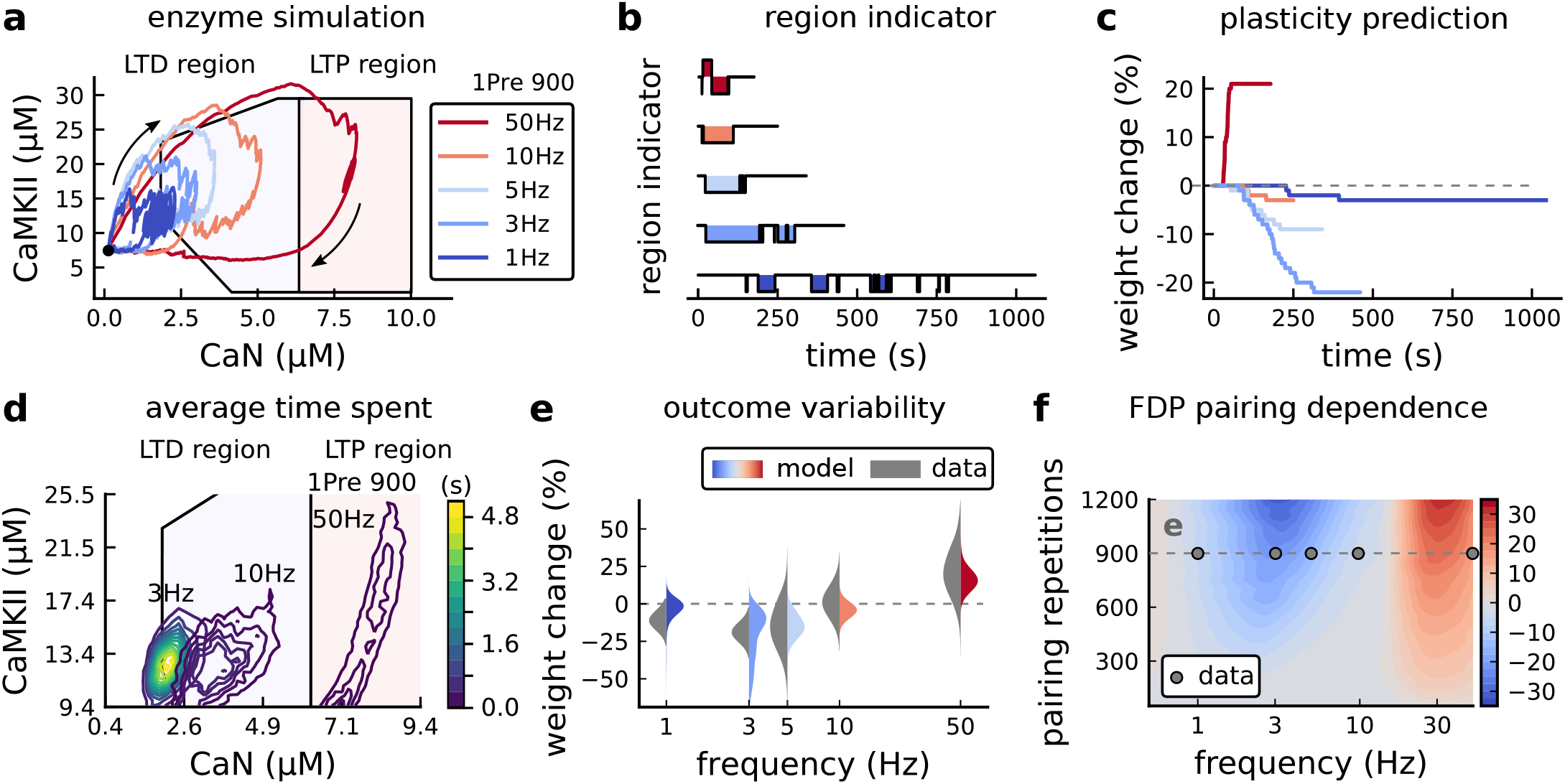
Frequency dependent plasticity, *Dudek and Bear (1992)* dataset. **a**, Example traces of joint CaMKII-CaN activity for each of ***Dudek and Bear (1992)*** protocol. **b**, Region indicator showing when the joint CaMKII-CaN activity crosses the LTD or LTP regions for each protocol in panel **a**. **c**, Synaptic weight change (%) as a function of time for each protocol, analogous to ***Figure 3c***. Trace colours correspond to panel **a**. The trajectories displayed were chosen to match the medians in panel **e**. **d**, Mean (100 samples) time spent (s) for protocols 1Pre for 900 pairing repetitions at 3,10 and 50 Hz. **e**, Comparison between data from ***Dudek and Bear (1992)*** and our model (1Pre 900p, 300 samples per frequency, see ***Table 1***). Data are represented as normal distributions with the mean and variance of the change in field EPSP slope taken from ***Dudek and Bear (1992)***. **f**, Prediction for the mean weight change (%) when varying the stimulation frequency and pulse number (24×38×100 data points, respectively pulse x frequency x samples). The filled grey circles show the ***Dudek and Bear (1992)*** protocol parameters and the corresponding results are shown in panel **e**. **Figure 4-Figure supplement 1.** Varying experimental parameters in ***Dudek and Bear (1992)*** and Poisson spike train during development.

**Figure 5.**
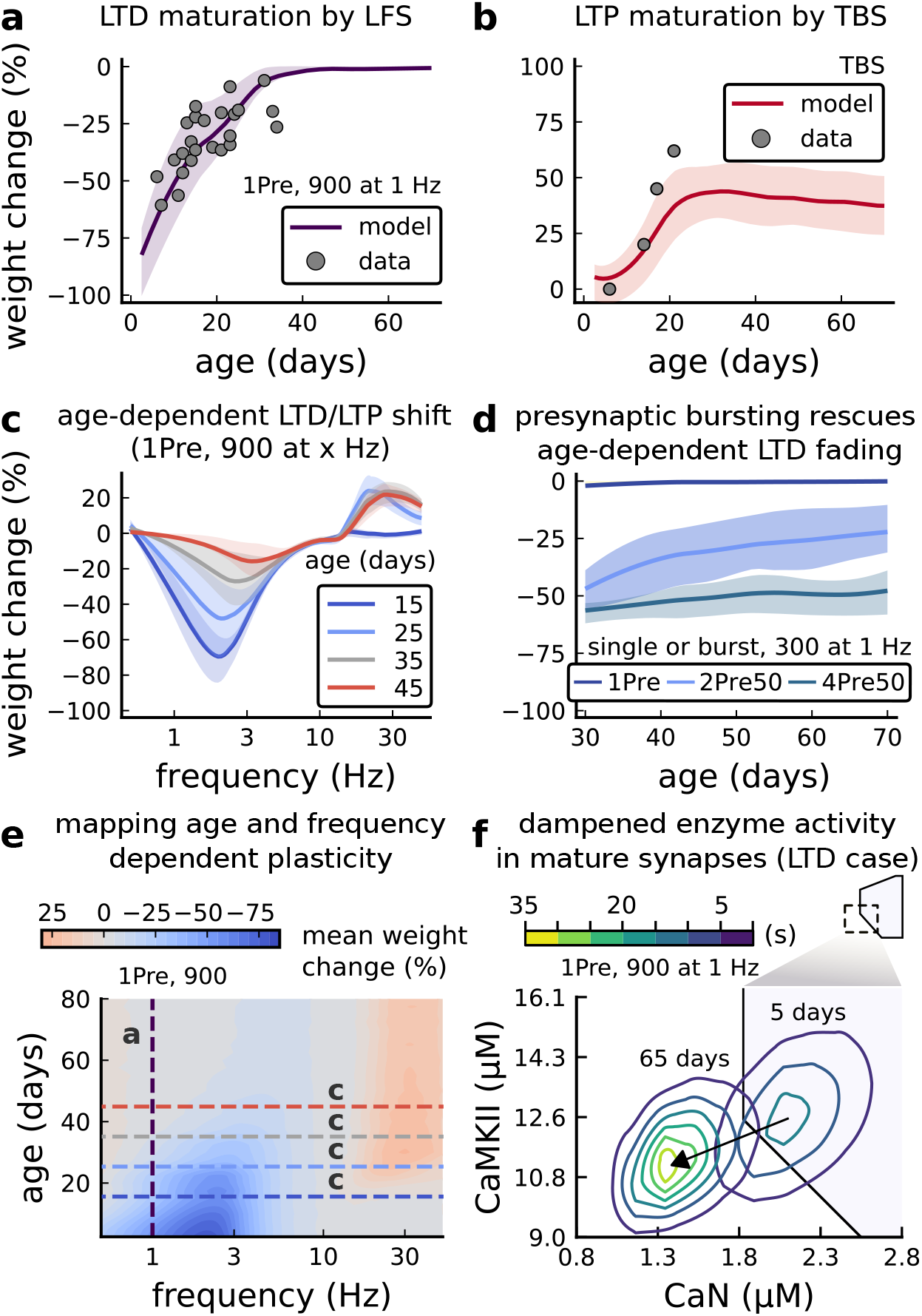
Age-dependent plasticity, *Dudek and Bear (1993)* dataset. **a**, Synaptic weight change for 1Pre, 900 at 1 Hz as in ***Dudek and Bear (1993)***. The solid line is the mean and the ribbons are the 2nd and 4th quantiles predicted by our model (same for panel **b**, **c** and **f**). **b**, Synaptic weight change for Theta Burst Stimulation (TBS – 4Pre at 100 Hz repeated 10 times at 5Hz given in 6 epochs at 0.1Hz, see ***Table 1***). **c**, Synaptic weight change as a function of frequency for different ages. BCM-like curves showing that, during adulthood, the same LTD protocol becomes less efficient. It also shows that high-frequencies are inefficient at inducing LTP before P15. **d**, Synaptic weight change as a function of age. Proposed protocol using presynaptic bursts to recover LTD at ? P35 with less pulses, 300 instead of the original 900 from ***Dudek and Bear (1993)***. This effect is more pronounced for young rats. ***Figure 5-Figure Supplement 1*** shows a 900 pulses comparison. **e**, Mean synaptic strength change (%) as a function of frequency and age for 1Pre 900 pulses (32×38×100, respectively, for frequency, age and samples). The protocols in ***Dudek and Bear (1993)*** (panel **a**) are marked with the yellow vertical line. The horizontal lines represent the experimental conditions of panel **c**. Note the P35 was used for ***Dudek and Bear (1992)*** experiment in ***Figure 4f***. **f**, Mean time spent for the 1Pre 1Hz 900 pulses protocol showing how the trajectories are left-shifted as rat age increases. **Figure 5-Figure supplement 1.** Duplets, triplets and quadruplets for FDP, perturbing developmental-mechanisms for LFS and HFS in ***Dudek and Bear (1993)***, and age-related changes in STDP experiments (***Inglebert et al., 2020; Tigaret et al., 2016; Meredith et al., 2003***).

**Figure 6.**
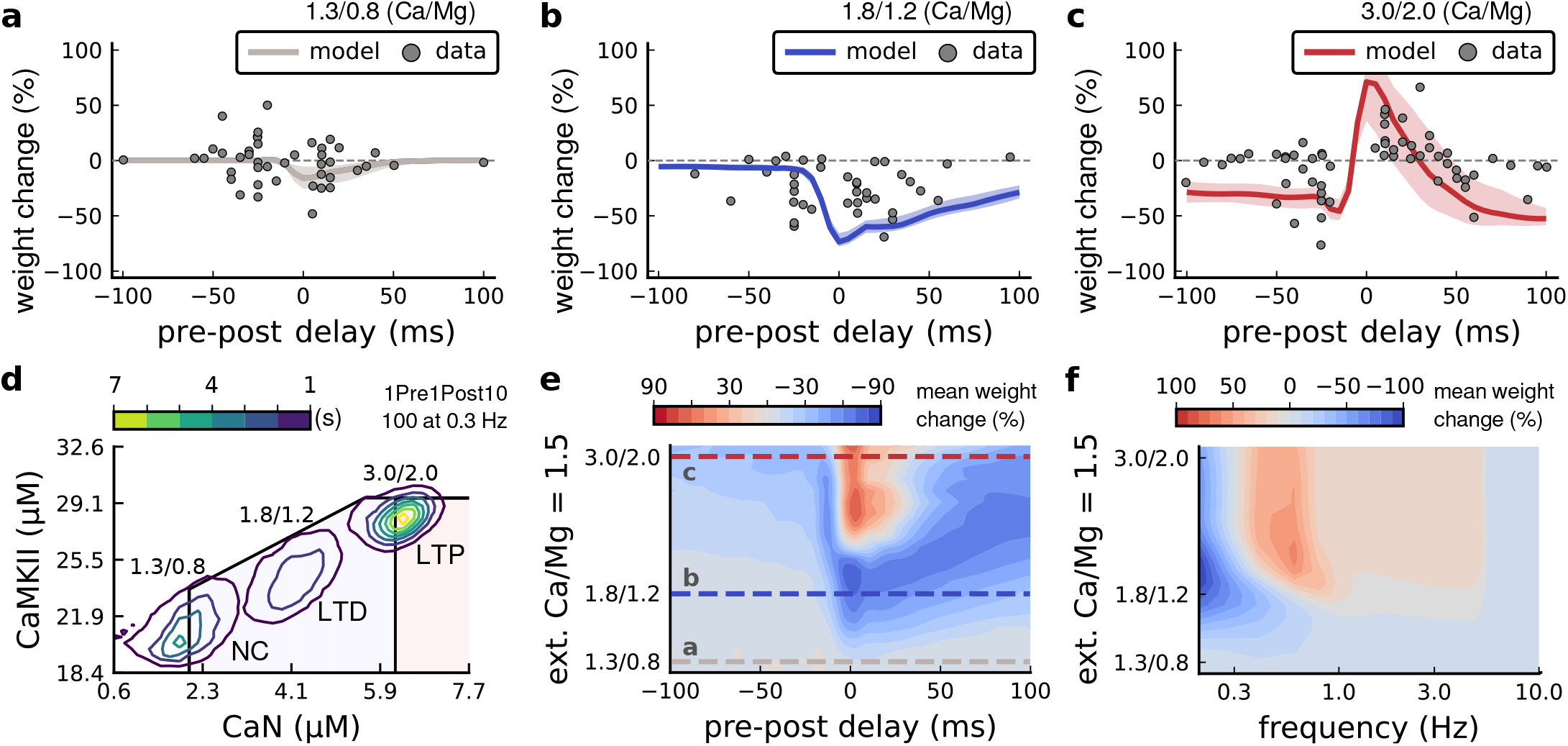
Effects of extracellular calcium and magnesium concentrations on plasticity. **a,** Synaptic weight (%) for a STDP rule with [Ca^2+^]_o_=1.3 mM (fixed ratio, Ca/Mg=1.5). According to the data extracted from ***Inglebert et al. (2020)***, the number of pairing repetitions for causal/positive (anti-causal/negative) delays is 100 (150), both delivered at 0.3 Hz. The solid line is the mean, and the ribbons are the 2nd and 4th quantiles predicted by our model (all panels use 100 samples). **b,** Same as **a**, but for [Ca^2+^]_o_ = 1.8 mM (Ca/Mg ratio = 1.5). **c,** Same as **a**, but for [Ca^2+^]_o_ = 3 mM (Ca/Mg ratio = 1.5). **d,** Mean time spent for causal pairing, 1Pre1Post10, at different Ca/Mg concentration ratios. The contour plots are associated with the panels **a**, **b** and **c**. **e,** Predicted effects of extracellular Ca/Mg on STDP outcome. Synaptic weight change (%) for causal (1Pre1Post10,100 at 0.3 Hz) and anticausal (1Post1Pre10,150 at 0.3 Hz) pairings varying extracellular Ca from 1.0 to 3 mM (Ca/Mg ratio = 1.5). The dashed lines represent the experiments in the panel **a**, **b** and **c**. We used 21×22×100 data points, respectively calcium x delay x samples. **f**, Predicted effects of varying frequency and extracellular Ca/Mg for an STDP protocol. Contour plot showing the mean synaptic weight (%) for a single causal pairing protocol (1Pre1Post10,100 samples) varying frequency from 0.1 to 10 Hz and [Ca^2+^]_o_ from 1.0 to 3 mM (Ca/Mg ratio = 1.5). We used 21×18×100 data points, respectively calcium x frequency x samples. **Figure 6-Figure supplement 1**. Effects of extracellular calcium and magnesium concentration on plasticity. **Figure 6-Figure supplement 2**. Temperature and age effects.

**Figure 7.**
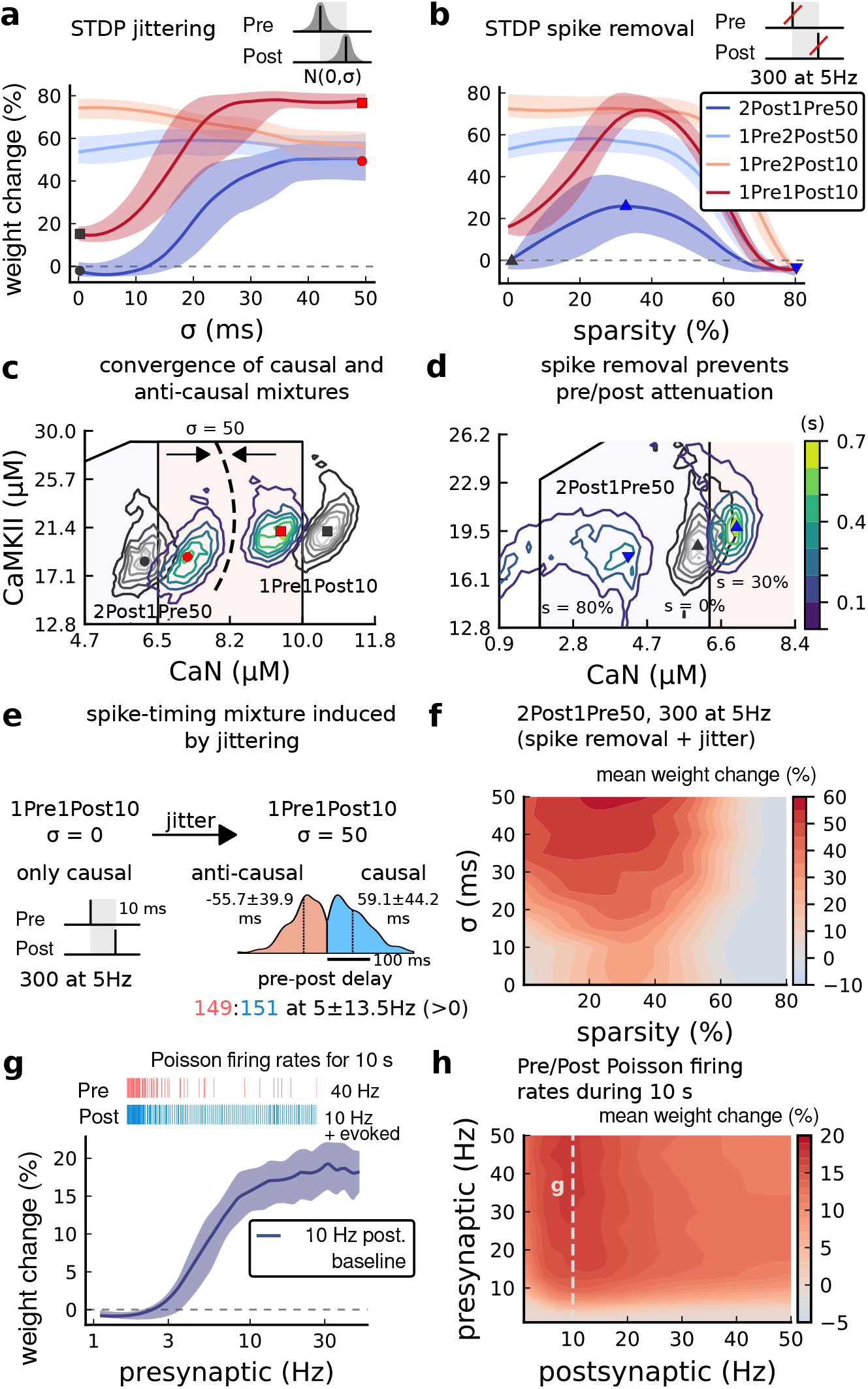
Jitter and spike dropping effects on STDP and Poisson spike trains. **a**, Mean weight (%) for the jittered STDP protocols (protocol color legend shown in **b**). The solid line is the mean, and the ribbons are the 2nd and 4th quantiles predicted by our model using 100 samples (same panels a, b and g). **b**, Mean weight (%) for the same (*Tigaret et al., 2016)* protocols used in panel **a** subjected to random spike removal (sparsity %). **c**, Mean time spent (s) varying jittering. Contour plot shows 2Post1Pre50 and 1Pre1Post10 (300 at 5 Hz) without (grey contour plot) and with jittering (coloured contour plot). The circles and squares correspond to the marks in panel **a**. **d**, Mean time spent (s) varying sparsity. Contour plot in grey showing 0% sparsity for 2Post1Pre50 300 at 5Hz (see ***Figure 2f***). The contour plots show the protocol with spike removal sparsities at 0% (NC), 30% (LTP), and 80% (NC). The triangles correspond to the same marks in panel **a**. **e**, Distribution of the 50 ms jittering applied to the causal protocol 1Pre1Post10, 300 at 5 Hz in which nearly half of the pairs turned into anticausal. The mean frequency is 5 ± 13.5 Hz making it to have a similar firing structure and position in the LTP region. The similar occurs for 2Post1Pre50 (panel c). **f**, Mean weight change (%) combining both jittering (panel **a**) and sparsity (panel **b**) for 2Post1Pre50, 300 at 5 Hz. **g**, Mean weight change (%) of pre and postsynaptic Poisson spike train delivered simultaneously for 10 s. The plot shows the plasticity outcome for different presynaptic firing rate (1000/frequency) for a fixed postsynaptic baseline at 10Hz. The upper raster plot depicts the released vesicles at 40 Hz and the postsynaptic baseline at 10Hz (including the AP evoked by EPSP). **h)**, Mean weight change (%) varying the rate of pre and postsynaptic Poisson spike train delivered simultaneously for 10 s. The heat map data along the vertical white dashed line is depicted in panel **g**.

The model components are schematized in ***Figure 1a*** (full details in ***Methods and Materials***). For glutamate release, we used a two-pool vesicle depletion and recycling system, which accounts for short-term presynaptic depression and facilitation. When glutamate is released from vesicles, it can bind to the postsynaptic α-amino-3-hydroxy-5-methyl-4-isoxazolepropionic acid and N-methyl-D-aspartate receptors (AMPArs and NMDArs, respectively), depolarizing the spine head by ~30 mV (***Kwon et al., 2017; Jayant et al., 2017; Beaulieu-Laroche and Harnett, 2018***). The dendritic spine membrane depolarization causes the activation of voltage-gated calcium channels (VGCCs) and removes magnesium ([Mg^2+^]_o_) block from NMDArs. Backpropagating action potentials (BaP) can also depolarize the spine membrane by up to ~60 mV (***Kwon et al., 2017; Jayant et al., 2017***). As an inhibitory component, we modelled a gamma-aminobutyric acid receptor (GABAr) synapse on the dendrite shaft (***Destexhe et al., 1998***). Calcium ions influx through VGCCs and NMDArs can activate SK potassium channels (***Adelman et al., 2012; Griffith et al., 2016***), which provide a tightly-coupled local negative feedback limiting spine depolarisation. Upon entering the spine, calcium ions also bind to calmodulin (CaM). Calcium-bound CaM in turn activates two major signalling molecules (***Fujii et al., 2013***): Ca^2+^/calmodulin-dependent protein kinase II (CaMKII) and calcineurin (CaN) phosphatase, also known as PP2B (***Saraf et al., 2018***). We included these two enzymes because of the overwhelming evidence that CaMKII activation is necessary for Schaffer-collateral LTP (***Giese et al., 1998; Chang et al., 2017***), while CaN activation is necessary for LTD (***O’Connor et al., 2005; Otmakhov et al., 2015***). Later, we show how we map the joint activity of CaMKII and CaN to LTP and LTD. Ligand-gated ion channels (ionotropic receptors) and voltage-gated ion channels have an inherent random behavior, stochastically switching between open and closed states (***Ribrault et al., 2011***). If the number of ion channels is large, then the variability of the total population activity becomes negligible relative to the mean (***O’Donnell and Van Rossum, 2014***). However individual hippocampal synapses contain only small numbers of receptors and ion channels, for example they contain ~10 NMDArs and <15 VGCCs (***Takumi et al., 1999; Sabatini and Svoboda, 2000; Nimchinsky et al., 2004***), making their total activation highly stochastic. Therefore, we modelled AMPAr, NMDAr, VGCCs and GABAr as stochastic processes. Presynaptic vesicle release events were also stochastic: glutamate release was an all-or-none event, and the amplitude of each glutamate pulse was drawn randomly, modelling heterogeneity in vesicle size (***Liu et al., 1999***). The inclusion of stochastic processes to account for an intrinsic noise in synaptic activation (***Deperrois and Graupner, 2020***) contrasts with most previous models in the literature, which either represent all variables as continuous and deterministic or add an external generic noise source (***Bhalla, 2004; Antunes and De Schutter, 2012; Bartol et al., 2015***).

The synapse model showed nonlinear dynamics across multiple timescales. For illustration, we stimulated the synapse with single simultaneous glutamate and GABA vesicle releases (***Figure 1b***). AMPArs and VGCCs open rapidly but close again within a few milliseconds. The dendritic GABAr closes more slowly, on a timescale of ~10 ms. NMDArs, the major calcium source, closes on timescales of ~50 ms and ~250 ms for the GluN2A and GluN2B subtypes, respectively.

To show the typical responses of the spine head voltage and Ca^2+^, we stimulated the synapse with a single presynaptic pulse (EPSP) paired 10 ms later with a single BaP (1Pre1Post10) (***Figure 1c left***). For this pairing, the arrival of a BaP at the spine immediately after an EPSP, leads to a large Ca^2+^ transient aligned with the BaP due to the NMDArs first being bound by glutamate then unblocked by the BaP depolarisation (***Figure 1c right***).

Single pre or postsynaptic stimulation pulses did not cause depletion of vesicle reserves or substantial activation of the enzymes. To illustrate these slower-timescale processes, we stimulated the synapse with a prolonged protocol: one presynaptic pulse followed by one postsynaptic pulse 10 ms later, repeated 30 times at 5 Hz (***Figure 1d-e***). The number of vesicles in both the docked and reserve pools decreased substantially over the course of the stimulation train (***Figure 1d left***), which in turn causes decreased vesicle release probability. Similarly, by the 30th pulse, the dendritic BaP amplitude had attenuated to ~85% (~70% BaP efficiency; ***Figure 1d right***) of its initial amplitude, modelling the effects of slow dendritic sodium channel inactivation (***Colbert et al., 1997; Golding et al., 2001***). Free CaM concentration rose rapidly in response to calcium transients but also decayed back to baseline on a timescale of ~500 ms (***Figure 1e top***). In contrast, the concentration of active CaMKII and CaN accumulated over a timescale of seconds, reaching a sustained peak during the stimulation train, then decayed back to baseline on a timescale of ~10 and ~120 s respectively, in line with experimental data (***Quintana et al., 2005; Fujii et al., 2013; Chang et al., 2017***) (***Figure 1e***).

The effects of the stochastic variables can be seen in ***Figure 1b-d***. The synaptic receptors and ion channels open and close randomly (***Figure 1b***). Even though spine voltage, calcium, and downstream molecules were modelled as continuous and deterministic, they inherited some randomness from the upstream stochastic variables. As a result, there was substantial trial-to-trial variability in the voltage and calcium responses to identical pre and postsynaptic spike trains (grey traces in ***Figure 1c***). This variability was also passed on to the downstream enzymes CaM, CaMKII and CaN, but was filtered and therefore attenuated by the slow dynamics of CaMKII and CaN. In summary, the model contained stochastic nonlinear variables acting over five different orders of magnitude of timescale, from ~1 ms to ~1 min, making it sensitive to both fast and slow components of input signals.

### Distinguishing between stimulation protocols using the CaMKII and CaN joint response

It has proven difficult for simple models of synaptic plasticity to capture the underlying rules and explain why some stimulation protocols induce plasticity while others do not. We tested the model’s sensitivity by simulating its response to a set of protocols used by ***Tigaret et al. (2016)*** in a recent *ex vivo* experimental study on adult (P50-55) rat hippocampus with blocked GABAr. We focused on three pairs of protocols (three rows in ***Figure 2***). For each of these pairs, one of the protocols experimentaly induced LTP or LTD, while the other subtly different protocol caused no change (NC) in synapse strength. Notably, three leading spike-timing and calcium-dependent plasticity models (***Song et al., 2000; Pfister and Gerstner, 2006; Graupner and Brunel, 2012***) could not fit these data (***Figure 3-Figure Supplement 1 a,b and c***). We thus asked if, by contrast, our new model could distinguish between each pair of protocols by assigning the correct plasticity outcome.

The first pair of protocols differed in intensity. A protocol which caused no plasticity consisted of 1 presynaptic spike followed 10 ms later by one postsynaptic spike repeated at 5 Hz for one minute (1Pre1Post10, 300 at 5Hz). The other protocol induced LTP, but differed only in that it included a postsynaptic doublet instead of a single spike (1Pre2Post10, 300 at 5Hz), implying a slightly stronger initial BaP amplitude. We first attempted to achieve separability by plotting CaMKII or CaN activities independently. As observed in the plots in ***Figure 2a***, it was not possible to set a single concentration threshold on either CaMKII or CaN that would discriminate between the protocols. This result was expected, at least for CaMKII, as recent experimental data demonstrates a fast saturation of CaMKII concentration in dendritic spines regardless of stimulation frequency (***Chang et al., 2017***).

To achieve better separability we set out to testa different approach, which was to combine the activity of the two enzymes, by plotting the joint CaMKII and CaN responses against each other on a 2D plane (***Figure 2b***). This innovative geometric plot is based on a mathematical concept of orbits from dynamical systems theory (***Meiss, 2007***). In this plot, the trajectories of two protocols can be seen to overlap for the initial part of the transient and then diverge. To quantify trial to trial variability, we also calculated contour maps showing the mean fraction of time the trajectories spent in each part of the plane during the stimulation (***Figure 2c***). Importantly, both the trajectories and contour maps were substantially non-overlapping between the two protocols, implying that they can be separated based on the joint CaN-CaMKII activity. We found that the 1Pre2Post10 protocol leads to a weaker response in both CaMKII and CaN, corresponding to the lower blue traces in ***Figure 2b***. The decreased response to the doublet protocol was due to the stronger attenuation of dendritic BaP amplitude over the course of the simulation (***Golding et al., 2001***), leading to reduced calcium influx through NMDArs and VGCCs (data not shown).

Using the second pair of protocols, we explored if this combined enzyme activity analysis could distinguish between subtle differences in protocol sequencing. We stimulated our model with one causal paring protocol (EPSP-BaP) involving a single presynaptic spike followed 50 ms stimulated our model with one causal paring protocol (EPSP-BaP) involving a single presynaptic spike followed 50 ms later by a doublet of postsynaptic spikes (1Pre2Post50, 300 at 5Hz), repeated at 5 Hz for one minute, which caused LTP in ***Tigaret et al. (2016)***. The other anticausal protocol involved the same total number of pre and postsynaptic spikes, but with the pre-post order reversed (2Post1Pre50, 300 at 5Hz). Experimentally the anticausal (BaP-EPSP) protocol did not induce plasticity (***Tigaret et al., 2016***). Notably, the only difference was the sequencing of whether the pre or postsynaptic neuron fired first, over a short time gap of 50 ms. Despite the activations being apparently difficult to distinguish (***Figure 2d***), we found that the LTP-inducing protocol caused greater CaN activation than the protocol that did not trigger plasticity. Indeed, this translated to a horizontal offset in both the trajectory and contour map (***Figure 2e-f***), demonstrating that another pair of protocols can be separated in the joint CaN-CaMKII plane.

The third pair of protocols differed in both duration and intensity. We thus tested the combined enzyme activity analysis in this configuration. In line with a previous study (***Isaac et al., 2009***), ***Tigaret et al. (2016)*** found that a train of doublets of presynaptic spikes separated by 50 ms repeated at a low frequency of 3 Hz for 5 minutes (2Pre50, 900 at 3Hz) induced LTD, while a slightly more intense but shorter duration protocol of presynaptic spike doublets separated by 10 ms repeated at 5 Hz for one minute (2Pre10, 300 at 5Hz) did not cause plasticity. When we simulated both protocols in the model (***Figure 2g-i***), both caused similar initial responses in CaMKII and CaN. In the shorter protocol, this activation decayed to baseline within 100 s of the end of the stimulation. However the slower and longer-duration 2Pre50 3Hz 900p protocol caused an additional sustained, stochastically fluctuating, plateau of activation of both enzymes (***Figure 2g***). This resulted in the LTD-inducing protocol having a downward and leftward-shifted CaN-CaMKII trajectory and contour plot, relative to the other protocol (***Figure 2h-i***). These results again showed that the joint CaN-CaMKII activity can predict plasticity changes.

### A geometrical readout mapping joint enzymatic activity to plasticity outcomes

The three above examples demonstrated that ploting the combined CaN-CaMKII activities in a 2D plane allowed us to distinguish between subtly different protocols with correct assignment of plasticity outcome. We found that the simulated CaN-CaMKII trajectories from the two LTP-inducing protocols (***Figure 2a*** and ***Figure 2d***) spent a large fraction of time near ~ 20 μM CaMKII and 7–10 μM CaN. In contrast, protocols that failed to trigger LTP had either lower (***Figure 2d and g***), or higher CaMKII and CaN activation (1Pre1Post10, ***Figure 2a***). The LTD-inducing protocol, by comparison, spent a longer period in a region of sustained but lower ~ *12μM* CaMKII and ~ 2*μ*M CaN and activation. The plots in ***Figure 2c, f and g***, show contour maps of histograms of the joint CaMKII-CaN activity, indicatingwhere in the plane the trajectories spent most time. ***Figure2c and f*** indicate that this measure can be used to predict plasticity, because the NC and LTP protocol histograms are largely non-overlapping. In ***Figure 2c***, the NC protocol response “overshoots” the LTP protocol response, whereas in ***Figure 2f*** the NC protocol response “undershoots” the LTP protocol response. In contrast, when we compared the response histograms for the LTD and NC protocols, we found a greater overlap (***Figure 2i***). This suggested that, in this case, the histogram alone was not sufficient to separate the protocols, and that protocol duration is also important. LTD induction (2Pre50) required a more prolonged activation than NC (2Pre10). We thus took advantage of these joint CaMKII-CaN activity maps to design a minimal readout mechanism connecting combined enzyme activity to LTP, LTD or no change (NC). We reasoned that this readout would need three key properties. First, since the CaMKII-CaN trajectories corresponding to LTP and LTD were not linearly separable, the readout requires nonlinear boundaries to activate the plasticity inducing components. Second, since LTD requires more prolonged activity than LTP, the readout should be sensitive to the timescale of the input. Third, a mechanism is required to convert the 2D LTP-LTD inducing signals into a synaptic weight change. After iterating through several designs, we satisfied the first property by designing “plasticity regions”: polygons in the CaN-CaMKII plane that would detect when trajectories pass through. We satisfied the second property by using two plasticity inducing components with different time constants which low-pass-filter the plasticity region signals. We satisfied the third property by feeding both the opposing LTP and LTD signals into a stochastic Markov chain which accumulated the total synaptic strength change. Overall this readout mechanism acts as a parsimonious model of the complex signalling cascade linking CaMKII and CaN activation to expression of synaptic plasticity (***He et al., 2015***). It can be considered as a two-dimensional extension of previous computational studies that applied analogous 1D threshold functions to dendritic spine calcium concentration (***Shouval et al., 2002; Karmarkar and Buonomano, 2002; Graupner and Brunel, 2012; Standage et al., 2014***).

We now elaborate on the readout design process. We first drew non-overlapping polygons of LTP and LTD “plasticity regions” in the CaN-CaMKII plane (***Figure 3a***). We positioned these regions over the parts of the phase space where the enzyme activities corresponding to the LTP- and LTD-inducing protocols were most different (***Methods and Materials***), as shown by trajectories in ***Figure 2***. When a trajectory enters in one of these plasticity regions, it activates LTD or LTP indicator variables (***Methods and Materials***) which encode the joint enzyme activities (trajectories in the phase plots) transitions across the LTP and LTD regions over time (***Figure 3b***). These indicator variables drove transition rates of a plasticity Markov chain used to predict LTP or LTD (***Figure 3c***), see ***Methods and Materials***. Intuitively, this plasticity Markov chain models the competing processes of insertion/deletion of AMPArs to the synapse, although this is not represented in the model. The LTD transition rates were slower than the LTP transition rates, to reflect studies showing that LTD requires sustained synaptic stimulation (***Yang et al., 1999; Mizuno et al., 2001; Wang et al., 2005***). The parameters for this plasticity Markov chain (***Methods and Materials***) were fit to the plasticity induction outcomes from different protocols (*Table 1*). At the beginning of the simulation, the plasticity Markov chain starts with 100 processes (***Destexhe et al., 1998***) in the state No Change (NC), with each variable representing 1% weight change, an abstract measure of synaptic strength that can be either EPSP, EPSC, or field EPSP slope depending on the experiment. Each process can transit stochastically between NC, LTP and LTD states. At the end of the protocol, the plasticity outcome is given by the difference between the number of processes in the LTP and the LTD states (***Methods and Materials***).

**Table 1.** Table with the parameters extracted from the respective publications. To fit the data associated to publications displaying a parameter interval (e.g. 70 to 100) we used a value within the provided limits. Otherwise, we depict in parentheses the value used to fit to the data. For complete data structure on these publications and the ones used for method validation see github code. We allowed the AP to be evoked by EPSPs for these protocols: ***Mizuno et al. (2001)***, ***Dudek and Bear (1992) Dudek and Bear (1993)***. Note that ***Tigaret et al. (2016)*** used GABA(A)r blockers, which we modelled by setting the GABAr conductance to zero. Also, ***Mizuno et al. (2001)*** LTD protocol used partial NMDA blocker modelled by reducing NMDA conductance by 97 %. **Table 1 – Table Supplement 1.** Comparison of recent computational models for plasticity. **Table 2 – Table Supplement 2.** Comparison of the experimental conditions for the different reproduced datasets in recent computational models for plasticity.

| Experiment | Paper | Repetitions | Freq (Hz) | Age (days) | Temp. (°C) | [Ca<sup>2+</sup>]<sub>o</sub>(mM) | [Mg<sup>2+</sup>]<sub>o</sub>(mM) |
| --- | --- | --- | --- | --- | --- | --- | --- |
| STDP | Tigaret et al. (2016) | 300 | 5 | 56 | 35 | 2.5 | 1.3 |
| STDP | Inglebert et al. (2020) | 100, positive delays<br>150, negative delays | 0.3 | 14—20<br>(21 for LTP) | 30<br>(30.45 for LTP) | 1.3—3 | Ca/1.5 |
| STDP | Meredith et al. (2003) | 20 | 0.2 | 9—45 | 24—28 | 2 | 2 |
| STDP | Wittenberg and Wang (2006) | 70—100 | 5 | 14—21 | 24—30<br>(22.5-23) | 2 | 1 |
| pre-burst | Tigaret et al. (2016) | 300 and 900 | 3 and 5 | 56 | 35 | 2.5 | 1.3 |
| FDP | Dudek and Bear (1992) | 900 | 1—50 | 35 | 35 | 2.5 | 1.5 |
| FDP | Dudek and Bear (1993) | 900 | 1 | 7—35 | 35 | 2.5 | 1.5 |
| TBS | Dudek and Bear (1993) | 3—4 (5) epochs | 4Pre at 100 Hz<br>(10x at 5Hz) | 6, 14 and 17 | 35 | 2.5 | 1.5 |
| LFS | Mizuno et al. (2001) | 1—600 | 1 | 12—28 | 30<br>(26.5-31) | 2.4 | 0 |

In ***Figure 3d***, we plot the model’s responses to seven different plasticity protocols used by ***Tigaretet al. (2016)*** by overlaying example CaMKII-CaN trajectories for each protocol with the LTP and LTD regions. The corresponding region indicators are plotted as function of time in ***Figure 3e***, and long-term alterations in the synaptic strength are plotted as function of time in ***Figure 3f***. The three protocols that induced LTP in the ***Tigaret et al. (2016)*** experiments spent substantial time in the LTP region, and so triggered potentiation. In contrast, the 1Pre1Post10 overshoots both regions, crossing them only briefly on its return to baseline, and so resulted in little weight change. The protocol that induced LTD (2Pre50, purple trace) is five times longer than other protocols, spending sufficient time inside the LTD region (***Figure 3f***). In contrast, two other protocols that spent time in the same LTD region of the CaN-CaMKII plane (2Post1Pre50 and 2Pre10) were too brief to induce LTD. These protocols were also not strong enough to reach the LTP region, so resulted in no net plasticity, again consistent with ***Tigaret et al. (2016)*** experiments.

We observed run-to-run variability in the amplitude of the predicted plasticity, due to the inherent stochasticity in the model. To ensure that stochastic components are necessary for adequate model behaviour, we compared stochastic and deterministic versions of the model with and without discrete presynaptic release and found that adding stochastic components indeed modified the model’s behaviour (***Figure 3-Figure Supplement 2***). Also, we confirmed that VGCCs are necessary for accurate modelling of ***Tigaret et al. (2016)*** data as blocking these channels reproduced the data obtained in VGCC blockers by Tigaret i.e. no potentiation could be elicited (***Figure 3-Figure Supplement 3***). Finally, we stress in ***Figure 3-Figure Supplement 4*** that the horizontal boundaries (related to CaMKII activity) are indeed necessary.

In ***Figure 3g***, we plot the distribution of the predicted plasticity from all the protocols (colours) of Tigaret alongside the experimental data (***Tigaret et al., 2016***). We find a very good correspondence between the model and experiments. Of note, data fitting of the experiments in ***Tigaret et al. (2016)*** (***Figure 3g***) was more accurate with our model than the fitting obtained with existing leading spike- or calcium-based STDP models (***Song et al., 2000; Pfister and Gerstner, 2006; Graupner and Brunel, 2012***), see ***Figure 3-Figure Supplement 1***.

Experimentally, LTP can be induced by few pulses while LTD usually requires stimulation protocols of longer duration (***Yang et al., 1999; Mizuno et al., 2001; Wang et al., 2005***). We incorporated this effect into the geometrical readout model by letting LTP have faster transition rates than LTD (***Figure 3c***). ***Tigaret et al. (2016)*** found that 300 repetitions of anticausal post-before-pre pairings did not cause LTD, in contrast to the canonical spike-timing-dependent plasticity curve (***Bi and Poo, 1998***). We hypothesized that LTD might indeed appear with the anticausal protocol (*Table 1*) if stimulation duration was increased. To explore this possibility in our model, we systematically varied the number of paired repetitions from 100 to 1200, and also co-varied the pre-post delay from −100 to 100 ms. ***Figure 3h*** shows a contour plot of the predicted mean synaptic strength change across for the 1Pre2Post(delay) stimulation protocol for different numbers of pairing repetitions. In ***Figure 3h***, a LTD window appears after ~500 pairing repetitions for some anticausal pairings, in line with our hypothesis. The magnitude of LTP also increases with pulse number, for causal positive pairings. For either 100 or 300 pairing repetitions, only LTP or NC is induced (***Figure 3i***). The model also made other plasticity predictions by varying ***Tigaret et al. (2016)*** experimental conditions (***Figure 3-Figure Supplement 5***). In summary, our geometrical readout reveals that the direction and magnitude of the change in synaptic strength can be predicted from the joint CaMKII-CaN activity in the LTP and LTD regions.

### Frequency-dependent plasticity

The stimulation protocols used by ***Tigaret et al. (2016)*** explored how subtle variations in pre and postsynaptic spike timing influenced the direction and magnitude of plasticity (see ***Table 1**) for experimental differences). In contrast, traditional synaptic plasticity protocols exploring the role of presynaptic stimulation frequency did not measure the timing of co-occurring postsynaptic spikes (***Dudek and Bear, 1992; Wang and Wagner, 1999; Kealy and Commins, 2010***). These studies found that long-duration low-frequency stimulation induces LTD, whereas short-duration high-frequency stimulation induces LTP, with a cross-over point of zero change at intermediate stimulation frequencies. In addition to allowing us to explore frequency-dependent plasticity (FDP), this stimulation paradigm also gave us further constraints to define the LTD polygon region in the model since in ***Tigaret et al. (2016)***, only one LTD case was available. For FDP, we focused on modelling the experiments from ***Dudek and Bear (1992)***, who stimulated Schaffer collateral projections to pyramidal CA1 neurons with 900 pulses in frequencies ranging from 1 Hz to 50 Hz. In addition to presynaptic stimulation patterns, the experimental conditions differed from ***Tigaret et al. (2016)*** in two other aspects: animal age and control of postsynaptic spiking activity (see **Table 1*** legend). We incorporated both age-dependence and EPSP-evoked-BaPs in our model (***Methods and Materials***). Importantly, the geometrical readout mechanism mapping joint CaMKII-CaN activity to plasticity remained identical for all experiments in this work.

***Figure 4a*** shows the joint CaMKII-CaN activity when we stimulated the model with 900 presynaptic spikes at 1,3, 5,10 and 50 Hz (***Dudek and Bear, 1992***). Higher stimulation frequencies drove stronger responses in both CaN and CaMKII activities (***Figure 4a***). ***Figure4b,c*** showthe corresponding plasticity region indicator for the LTP/LTD region threshold crossings and the synaptic strength change. From this set of five protocols, only the 50 Hz stimulation drove a response strong enough to reach the LTP region of the plane (***Figure 4a and d***). Although the remaining four protocols drove responses primarily in the LTD region, only the 3 and 5 Hz stimulations resulted in substantial LTD. The 1 Hz and 10 Hz stimulations resulted in negligible LTD, but for two distinct reasons. Although the 10 Hz protocol’s joint CaMKII-CaN activity passed through the LTD region of the plane (***Figure 4a and d***), it was too brief to activate the slow LTD mechanism built into the readout (***Methods and Materials***). The 1 Hz stimulation, on the other hand, was prolonged, but its response was too weak to reach the LTD region, crossing the threshold only intermittently (***Figure 4b, bottom trace***). Overall the model matched well the mean plasticity response found by ***Dudek and Bear (1992)***, see ***Figure 4e***, following a classic BCM-like curve as function of stimulation frequency (***Abraham et al., 2001; Bienenstock et al., 1982***).

We then used the model to explore the stimulation space in more detail by varying the stimulation frequency from 0.5 Hz to 50 Hz, and varying the number of presynaptic pulses from 50 to 1200. ***Figure 4f*** shows a contour map of the mean synaptic strength change (%) in this 2D frequency–pulse number space. Under ***Dudek and Bear (1992)*** experimental conditions, we found that LTD induction required at least ~300 pulses, at frequencies between 1Hz and 3Hz. In contrast, LTP could be induced using ~50 pulses at ~20Hz or greater. The contour map also showed that increasing the number of pulses (vertical axis in ***Figure 4e***) increases the magnitude of both LTP and LTD. This was paralleled by a widening of the LTD frequency range, whereas the LTP frequency threshold remained around ~20Hz, independent of pulse number.

The pulse-dependent amplification of synaptic weight predicted in ***Figure 4*** is also valid for ***Tigaret et al. (2016)*** experiment shown in ***Figure 3h***.

*Ex vivo* experiments in ***Dudek and Bear (1992)*** were done at 35°*C*. However, lower temperatures are more widely used for *ex vivo* experiments because they extend brain slice viability. We performed further simulations testing temperature modifications for ***Dudek and Bear (1992)*** experiment, predicting a strong effect on plasticity outcomes (***Figure 4-Figure Supplement 1d-f***).

### Variations in plasticity induction with developmental age

The rules for induction of LTP and LTD change during development (***Dudek and Bear, 1993; Cao and Harris, 2012***), so a given plasticity protocol can produce different outcomes when delivered to synapses from young animals versus mature animals. For example, when ***Dudek and Bear (1993)*** tested the effects of low-frequency stimulation (1 Hz) on CA3-CA1 synapses from rats of different ages, they found that the magnitude of LTD decreases steeply with age from P7 until becoming minimal in mature animals >P35 (***Figure 5a***, circles). Across the same age range, they found that a theta-burst stimulation protocol induced progressively greater LTP magnitude with developmental age (***Figure 5b***, circles). Paralleling this, multiple properties of neurons change during development: the NMDAr switches its dominant subunit expression from GluN2B to GluN2A (***Sheng et al., 1994; Popescu et al., 2004; Iacobucci and Popescu, 2017)***, the reversal potential of the receptor (GABAr) switches from depolarising to hyperpolarizing (***Rivera et al., 1999; Meredith et al., 2003; Rinetti-Vargas et al., 2017***), and the action potential backpropagates more efficiently with age (***Buchanan and Mellor, 2007***). These mechanisms have been proposed to underlie the developmental changes in synaptic plasticity rules because they are key regulators of synaptic calcium signalling (***Meredith et al., 2003; Buchanan and Mellor, 2007***). However, their sufficiency and individual contributions to the age-related plasticity changes are unclear and this has not been taken into account in any previous model. We incorporated these mechanisms in the model (***Methods and Materials***) by parameterizing each of the three components to vary with the animal’s postnatal age, to test if they could account for the age-dependent plasticity data.

We found that elaborating the model with age-dependent changes in NMDAr composition, GABAr reversal potential, and BaP efficiency, while keeping the same plasticity readout parameters, was sufficient to account for the developmental changes in LTD and LTP observed by ***Dudek and Bear (1993)*** (*Figure 5a,b*). We then explored the model’s response to protocols of various stimulation frequencies, from 0.5 to 50 Hz, across ages from P5 to P80 (***Figure 5c,e***). ***Figure 5c*** shows the synaptic strength change as function of stimulation frequency for ages P15, P25, P35 and P45. The magnitude of LTD decreases with age, while the magnitude of LTP increases with age. ***Figure 5e*** shows a contour plot of the same result, covering the age-frequency space.

The 1Hz presynaptic stimulation protocol in Dudek and Bear (1993) did not induce LTD in adult animals (Dudek and Bear, 1992). We found that the joint CaN-CaMKII activity trajectories for this stimulation protocol underwent an age-dependent leftward shift beyond the LTD region (***Figure 5f***). This implies that LTD is not induced in mature animals by this conventional LFS protocol due to insufficient activation of enzymes. In contrast, ***Tigaret et al. (2016)*** and ***Isaac et al. (2009)*** were able to induce LTD in adult rat tissue by combining LFS with presynaptic spike pairs repeated 900 times at 3 Hz. Given these empirical findings and our modelling results, we hypothesized that LTD induction in adult animals requires that the stimulation protocol: 1) causes CaMKII and CaN activity to stay more in the LTD region than the LTP region, and 2) is sufficiently long to activate the LTD readout mechanism. With experimental parameters used by ***Dudek and Bear (1993)***, this may be as short as 300 pulses when multi-spike presynaptic protocols are used since the joint CaMKII-CaN activity can reach the LTD region more quickly than for single spike protocols. We simulated two such potential protocols as predictions: doublet and quadruplet spike groups delivered 300 times at 1 Hz, with 50 ms between each pair of spikes in the group (***Figure 5d***). The model predicted that both these protocols induce LTD in adults, whereas as shown above, the single pulse protocol did not cause LTD. These findings suggest that the temporal requirements for inducing LTD may not be as prolonged as previously assumed, since they can be reduced by varying stimulation intensity. See ***Figure 5-Figure Supplement 1*** for frequency versus age maps for presynaptic bursts.

***Dudek and Bear (1993)*** also performed theta-burst stimulation (TBS, ***Table 1***) at different developmental ages, and found that LTP is not easily induced in young rats (***Cao and Harris, 2012***), as depicted in *Figure 5b*. The model qualitatively matches this trend, and also predicts that TBS induces maximal LTP around P21, before declining further during development (***Figure 5b***, green curve). Similarly, we found that high-frequency stimulation induces LTP only for ages >P15, peaks at P35, then gradually declines at older ages (***Figure 5e***). Note that in ***Figure 5b***, we used 6 epochs instead of 4 used by ***Dudek and Bear (1993)*** to increase LTP outcome which is known to washout after one hour for young rats (***Cao and Harris, 2012***).

In contrast to ***Dudek and Bear (1993)*** findings, other studies have found that LTP can be induced in hippocampus in young animals (<P15) with STDP. For example, Meredith et al. (2003) found that, at room temperature, 1Pre1Post10 induces LTP in young rats, whereas 1Pre2Post10 induces NC. This relationship was inverted for adults, with 1Pre1Post inducing no plasticity and 1Pre2Post10 inducing LTP (***Figure 5-Figure Supplement 7***).

Together, these results suggest that not only do the requirements for LTP/LTD change with age, but also that these age-dependencies are different for different stimulation patterns. Finally, we explore which mechanisms are responsible for plasticity induction changes across development in the FDP protocol (***Figure 5-Figure Supplement 1***) by fixing each parameter to young or adult values for the FDP paradigm. Our model analysis suggests that the NMDAr switch (***Iacobucci and Popescu, 2017***) is a dominant factor affecting LTD induction, but the maturation of BaP (***Buchanan and Mellor, 2007***) is the dominant factor affecting LTP induction, with GABAr shift having only a weak influence on LTD induction for ***Dudek and Bear (1993)*** FDP.

Plasticity requirements during development do not necessarily follow the profile in ***Dudek and Bear (1993)*** as shown by ***Meredith et al. (2003)*** STDP experiment. Our model shows that multiple developmental profiles are possible when experimental conditions vary within the same stimulation paradigm. This is illustrated in **Figure 6-Figure Supplement 2** a-c by varying the age of STDP experiments done in different conditions. We fitted well the data from ***Wittenberg and Wang (2006)*** by adapting the model with appropriate age and temperature.

### Effects of extracellular calcium and magnesium concentrations on plasticity outcome

The canonical STDP rule (***Bi and Poo, 1998***), measured in cultured neurons with high extracellular calcium ([Ca^2+^]_o_) and at room temperature, was recently found not to be reproducible at physiological [Ca^2+^]_o_ in CA1 brain slices (***Inglebert et al., 2020***). Instead, by varying the [Ca^2+^]_o_ and [Mg^2+^]_o_, ***Inglebert et al. (2020)*** found a spectrum of STDP rules with either no plasticity or full-LTD for physiological [Ca^2+^]_o_ conditions ([Ca^2+^]_o_< 1.8 mM) and a bidirectional rule for high [Ca^2+^]_o_ ([Ca^2+^]_o_> 2.5 mM), shown in ***Figure 6a-c***.

We attempted to reproduce ***Inglebert et al. (2020)*** findings by varying [Ca^2+^]_o_ and [Mg^2+^]_o_ with the following consequences for the model mechanisms (***Methods and Materials***). On the presynaptic side, [Ca^2+^]_o_ modulates vesicle release probability. On the postsynaptic side, high [Ca^2+^]_o_ reduces NMDAr conductance (***Maki and Popescu, 2014***), whereas [Mg^2+^]_o_ affects the NMDAr Mg^2+^ block (***Jahr and Stevens, 1990***). Furthermore, spine calcium influx activates SK channels, which hyperpolarize the membrane and indirectly modulate NMDAr activity (***Ngo-Anh et al., 2005; Griffith et al., 2016***).

***Figure 6a-c*** compares our model to ***Inglebert et al. (2020)*** STDP data at different [Ca^2+^]_o_ and [Mg^2+^]_o_. Note that ***Inglebert et al. (2020)*** used 150 pairing repetitions for the anti-causal stimuli and 100 pairing repetitions for the causal stimuli both delivered at 0.3 Hz. At [Ca^2+^]_o_=1.3 mM, ***Figure 6a*** shows that the STDP rule induced weak LTD for brief causal delays. At [Ca^2+^]_o_ = 1.8 mM, in ***Figure 6b***, the model predicted a full-LTD window. At [Ca^2+^]_o_ = 3 mM, in ***Figure 6c***, it predicted a bidirectional rule with a second LTD window for long causal pairings, previously theorized by ***Rubin et al. (2005)***.

***Figure 6d*** illustrates the time spent by the joint CaN-CaMKII activity for 1Pre1Post10 using ***Inglebert et al. (2020)*** experimental conditions. Each density plot corresponds to a specific specific Ca/Mg ratio as in ***Figure 6a-c***. The response under low [Ca^2+^]_o_ spent most time inside the LTD region, but high [Ca^2+^]_o_ shifts the trajectory to the LTP region. ***Figure 6-Figure Supplement 1a*** presents density plots for the anti-causal protocols.

***Inglebert et al. (2020)*** fixed the Ca/Mg ratio at 1.5, although aCSF formulations in the literature differ (see ***Table 1***). Figure 6-Figure Supplement 1d shows that varying the Ca/Mg ratio and [Ca^2+^]_o_ for ***Inglebert et al. (2020)*** experiments restrict LTP to Ca/Mg>1.5 and [Ca^2+^]_o_>1.8 mM.

Our model can also identify the transitions between LTD and LTP depending on Ca/Mg. ***Figure 6e*** shows a map of plasticity as function of pre-post delay and Ca/Mg concentrations and the parameters where LTP is induced for the 1Pre1Post10 protocol. Since plasticity rises steeply at around [Ca^2+^]_o_ = 2.2 mM (see ***Figure 6e***), small fluctuations in [Ca^2+^]_o_ near this boundary could cause qualitative transitions in plasticity outcomes. For anti-causal pairings, increasing [Ca^2+^]_o_ increases the magnitude of LTD (***Figure 6-Figure Supplement 1b*** illustrates this with ***Inglebert et al. (2020)*** data).

***Inglebert et al. (2020)*** also found that increasing the pairing frequency to 5 or 10 Hz results in a transition from LTD to LTP for 1Pre1Post10 at [Ca^2+^]_o_ = 1.8 mM (***Figure 6-Figure Supplement 1c***), similar frequency-STDP behaviour has been reported in the cortex (***Sjöström et al., 2001***). In ***Figure 6f***, we varied both the pairing frequencies and [Ca^2+^]_o_ and we observe similar transitions to ***Inglebert et al. (2020)***. However, the model’s transition for [Ca^2+^]_o_ = 1.8 mM was centred around 0.5 Hz, which was untested by ***Inglebert et al. (2020)***. The model predicted no plasticity at higher frequencies, unlike the data, that shows scattered LTP and LTD (see ***Figure 6-Figure Supplement 1c***). Another frequency dependent comparison, ***Figure3-Figure Supplement 5c*** and ***Figure 6-Figure Supplement 1h***, show that ***Tigaret et al. (2016)*** burst-STDP and ***Inglebert et al. (2020)*** STDP share a similar transition structure, different from ***Dudek and Bear (1992)*** FDP.

In contrast to ***Inglebert et al. (2020)*** results, we found that setting low [Ca^2+^]_o_ for ***Tigaret et al. (2016)*** burst-STDP abolishes LTP, and does not induce strong LTD (***Figure 3-Figure Supplement 5d***). For ***Dudek and Bear (1992)*** experiment, ***Figure 4-Figure Supplement 1d*** [Mg^2+^]_o_ controls a sliding threshold between LTD and LTP but not [Ca^2+^]_o_ (***Figure 4-Figure Supplement 1b***). For another direct stimulation experiment, ***Figure 6-Figure Supplement 1c*** shows that in an Mg-free medium, LTP expression requires fewer pulses (***Mizuno et al., 2001***).

Despite exploring physiological [Ca^2+^]_o_ and [Mg^2+^]_o_ Inglebert (***Inglebert et al., 2020***) use a non-physiological temperature (30°*C*) which extends T-type VGCC closing times and modifies the CaN-CaMKII baseline (***Figure 6-Figure Supplement 2i***). ***Figure 6-Figure Supplement 2g,h*** show comparable simulations for physiological temperatures. In summary, our model predicts that temperature can change STDP rules in a similar fashion to [Ca^2+^]_o_ (***Figure 6-Figure Supplement 1a,b***). Overall, we confirm that plasticity is highly sensitive to variations in extracellular calcium, magnesium, and temperature (***Figure 3-Figure Supplement 5a, Figure 6-Figure Supplement 2d-f***).

### In vivo-like spike variability affects plasticity

In the above sections, we used highly regular and stereotypical stimulation protocols to replicate typical *ex vivo* plasticity experiments. In contrast, neural spiking in hippocampus *in vivo* is irregular and variable (***Fenton and Muller, 1998; Isaac et al., 2009***). Previous studies that asked how natural firing variability affects the rules of plasticity induction used simpler synapse models (***Rackham et al., 2010; Graupner et al., 2016; Cui et al., 2018***). We explored this question in our synapse model using simulations with three distinct types of additional variability: 1) spike time jitter, 2) failures induced by dropping spikes, 3) independent pre and postsynaptic Poisson spike trains (***Graupner et al., 2016***).

We introduced spike timing jitter by adding zero-mean Gaussian noise (s.d. *σ*) to pre and postsynaptic spikes, changing spike pairs inter-stimulus interval (ISI). In ***Figure 7a***, we plot the LTP magnitude as function of jitter magnitude (controlled by *σ*) for protocols taken from ***Tigaret et al. (2016)***. With no jitter, *σ* = 0, these protocols have different LTP magnitudes (corresponding to ***Figure 3***) and become similar once *σ* increases. The three protocols with a postsynaptic spike doublet gave identical plasticity for *σ* = 50 ms.

To understand the effects of jittering, we plotted the trajectories of joint CaN-CaMKII activity (***Figure 7c***). 2Post1Pre50 which “undershoots” the LTP region shifted into the LTP region for jitter *σ* = 50 ms. In contrast, 1Pre1Post10 which “overshoots” the LTP region shifted to the opposite direction towards the LTP region.

Why does jitter cause different spike timing protocols to yield similar plasticity magnitudes? Increasing jitter causes a fraction of pairings to invert causality. Therefore, the jittered protocols became a mixture of causal and anticausal pairings (***Figure 7c***). This situation occurs for all paired protocols. So any protocol with the same number spikes will produce a similar outcome if the jitter is large enough. Note that despite noise the mean frequency was conserved at 5 ± 13.5 Hz (see ***Figure 7e***).

Next, we studied the effect of spike removal. In the previous sections, synaptic release probability was ~60% (for [Ca^2+^]_o_ = 2.5 mM) or lower, depending on the availability of docked vesicles (***Methods and Materials***). However, baseline presynaptic vesicle release probability is heterogeneous across CA3-CA1 synapses, ranging from ~ 10 – 90% (***Dobrunz et al., 1997; Enoki et al., 2009***) and likely lower on average *in vivo* (***Froemke and Dan, 2002; Borst, 2010***). BaPs are also heterogeneous with random attenuation profiles (***Golding et al., 2001***) and spike failures (***Short et al., 2017***). To test the effects of pre and postsynaptic failures on plasticity induction, we performed simulations where we randomly removed spikes, altering the regular attenuation observed in ***Tigaret et al. (2016)*** protocols.

In ***Figure 7b*** we plot the plasticity magnitude as function of sparsity (percentage of removed spikes). The sparsity had different specific effects for each protocol. 1Pre2Post10 and 1Pre2Post50 which originally produced substantial LTP were robust to spike removal until ~ 60% sparsity. In contrast, the plasticity magnitude from both 1Pre1Post10 and 2Post1Pre50 showed a non-monotonic dependence on sparsity, first increasing then decreasing, with maximal LTP at ~40% sparsity.

To understand how sparsity causes this non-monotonic effect on plasticity magnitude, we plotted the histograms of time spent in the CaN-CaMKII plane for 2Post1Pre50 for three levels of sparsity: 0%, 30% and 80% (***Figure 7d** these protocols have diff*). For 0% sparsity, the activation spent most time at the border between the LTP and LTD regions, resulting in no change. Increasing sparsity to 30% caused the activation to shift rightward into the LTP region because there was less attenuation of pre and postsynaptic resources. In contrast, at 80% sparsity, the activation moved into the LTD region because there were not enough events to substantially activate CaMKII and CaN. Since LTD is a slow process and the protocol duration is short (60s), there was no net plasticity. Therefore for this protocol, high and low sparsity caused no plasticity for distinct reasons, whereas intermediate sparsity enabled LTP by balancing resource depletion with enzyme activation.

Next we tested the interaction of jitter and spike removal. ***Figure 7f*** shows a contour map of weight change as a function of jitter and sparsity for the 2Post1Pre50 protocol, which originally induced no plasticity (***Figure 2***). Increasing spike jitter enlarged the range of sparsity inducing LTP. In summary, these simulations (***Figure 7a,b,f and h***) show that different STDP protocols have different degrees of sensitivity to noise in the firing structure, suggesting that simple plasticity rules derived from regular *ex vivo* experiments may not predict plasticity *in vivo.*

How does random spike timing affect rate-dependent plasticity? We stimulated the model with pre and postsynaptic Poisson spike trains for 10s, under Dudek and Bear (1992) experimental conditions. We systematically varied both the pre and postsynaptic rates (***Figure 7h***). The 10s stimulation protocols induced only LTP, since LTD requires a prolonged stimulation (***Mizuno et al., 2001***). LTP magnitude monotonically increased with the presynaptic rate (***Figure 7g and h***). In contrast, LTP magnitude varied non-monotonically as a function of postsynaptic rate, initially increasing until a peak at 10 Hz, then decreasing with higher stimulation frequencies. This non-monotonic dependence on post-synaptic rate is inconsistent with classic rate-based models of Hebbian plasticity. We also investigated how this plasticity dependence on pre- and postsynaptic Poisson firing rates varies with developmental age (***Figure 4-Figure Supplement 1g-i***). We found that at P5 no plasticity is induced, at P15 a LTP region appears at around 1 Hz postsynaptic rate, and at P20 plasticity becomes similar to the mature age, with a peak in LTP magnitude at 10 Hz postsynaptic rate.

## Discussion

We built a model of a rat CA3-CA1 hippocampal synapse, including key electrical and biochemical components underlying synaptic plasticity induction (***Figure 1***). We developed a novel geometric readout of combined CaN-CaMKII dynamics (***Figure 2–Figure 4***) to predict the outcomes from a range plasticity experiments with heterogeneous conditions: animal developmental age (***Figure 5***), aCSF composition (***Figure 6***), temperature (***Supplemental files***), and *in vivo*-like firing variability (***Figure 7***). This readout provides a simple and intuitive window into the dynamics of the synapse during plasticity. Our model is thus based on the joint activity of these two key postsynaptic enzymes at both fast and slow time scales and considers the stochastic and adaptable dynamics of their activities dictated by the upstream calcium-dependent components at both the pre- and postsynapse. On this basis alone, our model is akin to biological processes where the outcome is jointly determined by several stochastic signaling components and a combination of multiple enzyme activities in time and space, i.e., are multi-dimensional. Our model is scalable, as it gives the possibility for the readout to be extended to dynamics of *n* different molecules, using *n*-dimensional closed regions. It is abstract in the sense that we do not identify the readout components with specific synaptic molecules. Nevertheless, we anticipate that simple biochemical networks could implement the readout’s functional mapping (***Alon, 2019***).

In addition to providing a new model of CA3-CA1 synapse biophysics, the main contribution of this work is the novel readout mechanism mapping synaptic enzymes to plasticity outcomes. This readout was built based on the concept that the full temporal activity of CaN-CaMKII over the minutes-timescale stimulus duration, and not their instantaneous levels, is responsible for changes in synaptic efficacy (***Fujii et al., 2013***). The readaout follows the measurements of CaMKII and CaN molecular dynamics made using FRET imaging (***Fujii et al., 2013***). CaMKII and CaN were chosen because they act upstream of several biochemical pathways implicated in the expression of plasticity and their inhibition blocks LTP and LTD, respectively (O'Connor et al., 2005). We expect that future studies using high temporal resolution measurements such as those provided by recent FRET tools available for CaMKII (***Chang et al., 2017, 2019***) will bring refinements to our model with the possibility to further test our readout predictions. In contrast, previous models assume that plasticity is explainable in terms of synaptic calcium or enzyme response to single BAP-EPSP pairings (***Shouval et al., 2002; Karmarkar and Buonomano, 2002***). We instantiated this concept by analyzing the joint CaN-CaMKII activity in the two-dimensional plane and designing polygonal plasticity readout regions (***Figure 3a***). In doing so, we generalised previous work with plasticity induction based on single threshold and a slow variable (***Badoual et al., 2006; Rubin et al., 2005; Clopath and Gerstner, 2010; Graupner and Brunel, 2012***) Given the high number of parameters in the model, we do not expect that the specific readout parameters we fit are unique. The addition of new datasets could better constrain the model fit. Here, we used only a two-dimensional readout, but anticipate a straightforward generalisation to higher-dimensions. The central discovery is that these trajectories, despite being stochastic, can be separated in the plane as a function of the stimulus (***Figure 3***). This is the basis of our new synaptic plasticity rule.

Let us describe the intuition behind our model more concisely. First, we abstracted away the sophisticated cascade of plasticity expression. Second, the plasticity regions, crossed by the trajectories, are described with a minimal set of parameters. Importantly, their tuning is quite straightforward and done only once, even when the joint activity is stochastic. The tuning of the model is possible thanks to the decoupling of the plasticity process from the spine biophysics which acts as a feedforward input to the plasticity Markov chain and from the distributions of the different trajectories, which are well separated. It is expected that one could find other versions of this model (parameters or conceptual) instantiating our multidimensional readout concept that also match the data well. The separability afforded by the geometrical readout, along with the model flexibility via fitting the plasticity regions, enabled us to reproduce data from nine different experiments using a single fixed set of model parameters. In contrast, we found that classic spike-timing (***Song et al., 2000; Pfister and Gerstner, 2006***) or calcium-threshold (***Graupner and Brunel, 2012***) models could not reproduce the range of protocols from ***Tigaret et al. (2016)*** (***Figure 3-Figure Supplement 1***). More complicated molecular-cascade models have been shown to account for individual plasticity experiments (***Antunes et al., 2016; Jędrzejewska-Szmek et al., 2017; Mäki-Marttunen et al., 2020; Bhalla, 2017***), but have not been demonstrated to reproduce the wide range of protocols presented here while considering experimental heterogeneity.

For some protocols, the CaMKII-CaN trajectories overshot the plasticity regions (e.g. ***Figure 3d***). Although abnormally high and prolonged calcium influx to cells can trigger cell death (***Zhivotovsky and Orrenius,2011***), the effects of high calcium concentrations at single synapses are poorly understood. Notably, a few studies have reported evidence consistent with an overshoot, where strong synaptic calcium influx does not induce LTP (***Yang et al., 1999; Tigaret et al., 2016; Pousinha et al., 2017***).

Our model included critical components for plasticity induction at CA3-CA1 synapses: those affecting dendritic spine voltage, calcium signalling, and enzymatic activation. We were able to use our model to make quantitative predictions, because its variables and parameters corresponded to biological components. This property allowed us to incorporate the model components’ dependence on developmental age, external Ca/Mg levels, and temperature to replicate datasets across a range of experimental conditions. The model is relatively fast to simulate, taking ~1 minute of CPU time to run 1 minute of biological time. These practical benefits should enable future studies to make experimental predictions on dendritic integration of multiple synaptic inputs (***Blackwell et al., 2019; Oliveira et al., 2012; Ebner et al., 2019***) and on the effects of synaptic molecular alterations in pathological conditions. In contrast, abstract models based on spike timing (***Song et al., 2000; Pfister and Gerstner, 2006; Clopath and Gerstner, 2010***) or simplified calcium dynamics (***Shouval et al., 2002; Graupner and Brunel, 2012***) must rely on ad hoc adjustment of parameters with less biological interpretability.

Intrinsic noise is an essential component of the model. How can the synapse reliably express plasticity but be noisy at the same time (***Yuste et al., 1999; Ribrault et al., 2011***)? Noise can be reduced either by redundancy or by averaging across time, also called ergodicity (***Sterling and Laughlin, 2015***). However redundancy requires manufacturing and maintaining more components, and therefore costs energy. We propose that, instead, plasticity induction is robust due to temporal averaging by slow-timescale signalling and adaptation processes. These slow variables display reduced noise levels by averaging the faster timescale stochastic variables. This may be a reason why CaMKII uses auto-phosphorylation to sustain its activity and slow its decay time (***Chang et al., 2017, 2019***). In summary, this suggests that the temporal averaging by slow variables, combined with the separability afforded by the multidimensional readout, allows synapses to tolerate noise while remaining energy-efficient.

A uniqueness of our model is that it simultaneously incorporates biological variables such as electrical components at pre and postsynaptic sites some with adaptive functions such as attenuation, age and temperature, stochastic noise and fast and slow timescales. Some of these variables have been modelled by other groups, e.g. stochasticity, BaP attenuation or pre-synaptic plasticity (***Cai et al., 2007; Shouval and Kalantzis, 2005; Zeng and Holmes, 2010; Miller et al., 2005; Yeung et al., 2004; Shah et al., 2006; Deperrois and Graupner, 2020; Costa et al., 2015***), but generally independently from each other. To position the uniqueness of our model in this broader context, we also provide a direct comparison of our model with some of the most recent leading models of excitatory synapse plasticity and the experimental work they reproduce (***Table 1-Table Supplement 1 and Table 1-Table Supplement 2***).

We identified some limitations of the model. First, we modelled only a single postsynaptic spine attached to a two-compartment neuron (soma and dendrite), see Model Compartments in Online Methods. Second, the model abstracted the complicated process of synaptic plasticity expression. Indeed, even if this replicated the early phase of LTP/LTD expression in the first 30–60 minutes after induction, we did not take into account slower protein-synthesis-dependent processes, maintenance processes, and synaptic pruning proceed at later timescales (***Bailey et al., 2015***). Third, like most biophysical models, ours contained many parameters (***Methods and Materials***). Although we set these to physiologically plausible values and then tuned to match the plasticity data, other combinations of parameters may fit the data equally well (***Marder and Taylor, 2011; Mäki-Marttunen et al., 2020***) due to the ubiquitous phenomenon of redundancy in biochemical and neural systems (***Gutenkunst et al., 2007; Marder, 2011***). Indeed synapses are quite heterogeneous in receptor and ion channel counts (***Takumi et al., 1999; Sabatini and Svoboda, 2000; Racca et al., 2000; Nimchinsky et al., 2004***), protein abundances (***Shepherd and Harris, 1998; Sugiyama et al., 2005***), and spine morphologies (***Bartol et al., 2015; Harris and Stevens, 1989***), even within the subpopulation of CA1 pyramidal neuron synapses that we modelled here. It remains to be discovered how neurons tune their synaptic properties in this vast parameter space to achieve functional plasticity rules, or implement meta-plasticity (***Huang et al., 1992; Deisseroth et al., 1995; Abraham, 2008***). Fourth, the activation of clustered synapses could influence the plasticity outcome, and the number of synapses activated during plasticity induction can be difficult to control experimentally. Our model concerns plasticity at a single synapse, which is also important during synaptic cluster activation (***Ujfalussy and Makara, 2020***). We drew from data in ***Tigaret et al. (2016)*** where there is little indication of simultaneous clustered synaptic activation. Furthermore, our simulations are in good agreement with plasticity experiments using local field potential recordings (***Dudek and Bear, 1993***) where the number of activated synapses is uncertain. This indicates that the model proposed here can account for this aspect of synaptic plasticity heterogeneity. Finally, our readout model does not correspond to a specific molecular cascade beyond CaN and CaMKII activations. However, we anticipate that the same mapping could be implemented by simple biochemical reaction networks, with for example, transition rates based on Hill functions for the plasticity boundaries. Future work could try to match this readout to known synaptic molecules.

Several predictions follow from our results. Since the model respected the stochasticity of vesicle release (***Rizzoli and Betz, 2005; Alabi and Tsien, 2012***), NMDAr (***Nimchinsky et al., 2004; Popescu et al., 2004; Iacobucci and Popescu, 2017; Sinclair et al., 2016***), and VGCC opening (***Magee and Johnston, 1995; Sabatini and Svoboda, 2000; Iftinca et al., 2006***), the magnitude of plasticity varied from simulation trial to trial (***Methods and Materials, Figure 3g and Figure 4e***). This suggests that the rules of plasticity are inherently stochastic (***Bhalla, 2004; Antunes et al., 2016***) and that the variability observed in these experiments (***Inglebert et al., 2020; Tigaret et al., 2016; Dudek and Bear, 1992, 1993; Mizuno et al., 2001; Meredith et al., 2003; Wittenberg and Wang, 2006***) is partly due to stochastic signalling, in addition to the previously-documented heterogeneity in synapse properties (***Nusser, 2018***) that we did not study here. By running extensive simulations over the space of protocols beyond those tested experimentally (***Figure 3h,i; Figure 4f; Figure 5c,e and f; Figure 6e,f***), we made testable predictions for plasticity outcomes. For example, ***Tigaret et al. (2016)*** did not find LTD when using classic post-before-pre stimulation protocols, but the model predicted that LTD could be induced if the number of pairing repetitions was extended (***Figure 3h,i***). The model also predicts that the lack of LTD induced by FDP in adults can be recovered using doublets or quadruplet spike protocols (***Figure 5d***). We tested the model’s sensitivity to spike time jitter and spike failure in the stimulation protocols (***Figure 7***). Our simulations predicted that this firing variability can alter the rules of plasticity, in the sense that it is possible to add noise to cause LTP for protocols that did not otherwise induce plasticity.

What do these results imply about the rules of plasticity *in vivo?* First, we noticed that successful LTP or LTD induction required a balance between two types of slow variables: those that attenuate, such as presynaptic vesicle pools and dendritic BaP, versus those that accumulate, such as slow enzymatic integration (***Cai et al., 2007; Mizusaki et al., 2018; Deperrois and Graupner, 2020***). This balance is reflected in the inverted-U shaped magnitude of LTP seen as a function of post-synaptic firing rate (***Figure 7h***). Second, although spike timing on millisecond timescales can in certain circumstances affect the direction and magnitude of plasticity (***Figure 3***), in order to drive sufficient activity of synaptic enzymes, these patterns would need to be repeated for several seconds. However, if these repetitions are subject to jitter or failures, as observed in hippocampal spike trains *in vivo* (***Fenton and Muller, 1998; Wierzynski et al., 2009***), then the millisecond-timescale information will be destroyed as it gets averaged out across repetitions by the slow integration processes of CaMKII and CaN (***Figure 7a-d***). The net implication is that millisecond-timescale structure of individual spike pairs is unlikely to play an important role in determining hippocampal synaptic plasticity *in vivo* (***Froemke and Dan, 2002; Sadowski et al., 2016; Graupner et al., 2016***).

In summary, we presented a new type of biophysical model for plasticity induction at the rat CA3-CA1 glutamatergic synapse. Although the model itself is specific to this synapse type, the study’s insights may generalise to other synapse types, enabling a deeper understanding of the rules of synaptic plasticity and brain learning.

## Methods and Materials

### Data and code availability

All simulations were performed in the Julia programming language (version 1.4.2). This choice was dictated by simplicity and speed (***Perkel, 2019***). The code for the Markov chains is mostly automatically generated from reactions, and could be exported to an SBML representation for porting to other languages.

Simulating the synapse model is equivalent to sampling a piecewise deterministic Markov process, and this relies on the thoroughly tested Julia package PiecewiseDeterministicMarkovProcesses.jl. These simulations are event-based, and no approximation is made beyond the ones required to integrate the ordinary differential equations by the LSODA method (Livermore Solver for Ordinary Differential Equations). We ran the parallel simulations in the Nef cluster operated by Inria.

### Notation

We write **1**_*A*_ for the indicator of a set *A*, meaning that **1**_*A*_(*x*) = 1 if *x* belongs to *A* and zero otherwise.

### Vesicle release and recycling

Vesicle-filled neurotransmitters from the presynaptic terminals stimulate the postsynaptic side when successfully released. We derived a vesicle release Markov chain model based on a deterministic approach described in ***Sterratt et al. (2011)***. We denote by (*t*_1_, –, *t_n_*) the arrival times of the presynaptic spikes.

Vesicles can be in two states, either belonging to the docked pool (with cardinal *D*) with fast emptying, or to the reserve pool (with cardinal *R*) which replenishes *D* (***Rizzoli and Betz, 2005***). Initially the docked and reserve pools have *D*_0_ and *R*_0_ vesicles, respectively. The docked pool loses one vesicle each time a release occurs (***Rudolph et al., 2015***), with transition *D* → *D* – 1 (***Figure 8***). The reserve pool replenishes the docked pool with transition (*R, D*) → (*R* – 1, *D* + 1). Finally, the reserve pool is replenished with rate 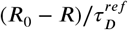 with the transition (*R,D*) → (*R* + 1,*P*).

**Figure 8.**
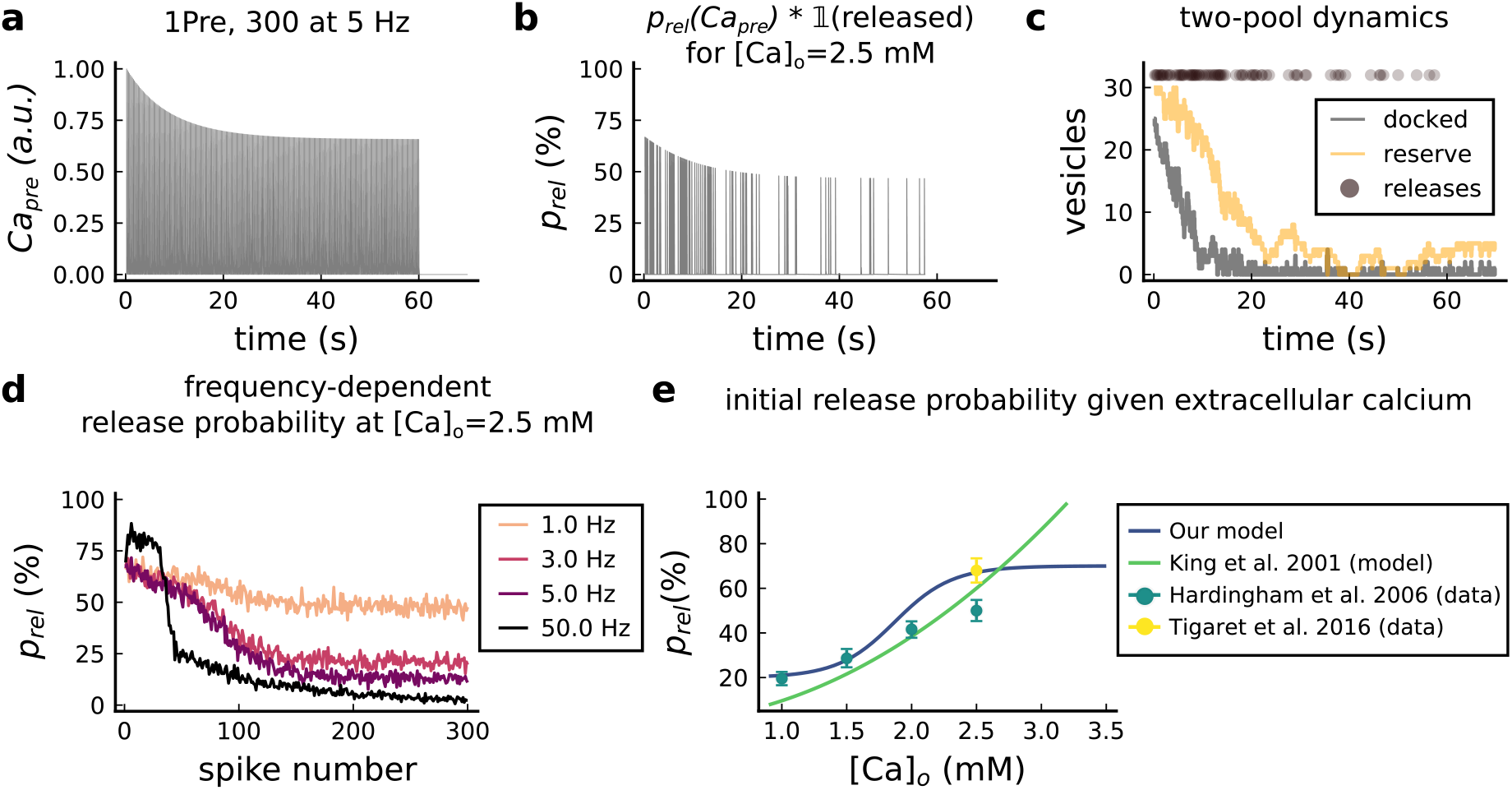
Presynaptic release. **a**, Presynaptic calcium in response to the protocol 1Pre, 300 at 5 z displaying adaptation. **b**, Release probability for the same protocol as panel A but subjected to the docked vesicles availability. **c**, Number of vesicles in the docked and reserve pools under depletion caused by the stimulation from panel **a**. **d**, Plot of the mean (300 samples) release probability (%) for different frequencies for the protocol 1Pre 300 pulses at [Ca^2+^]_o_ = 2.5 mM. **e**, Release probability (%) for a single presynaptic spike as a function of [Ca^2+^]_o_. Note that *King et al. (2001)* model was multiplied by the experimentally measured release probability at [Ca^2+^]_o_ = 2 mM since their model has this calcium concentration as the baseline. Our model also does not cover the abolishing of release probability at [Ca^2+^]_o_ = 0.5 mM which can also be difficult to measure experimentally given the rarity of events (***Hardingham et al., 2006***).

In addition to the stochastic dynamics in ***Table 2***, each spike *t_t_* triggers a vesicle release *D* → *D* – 1 with probability *p_rel_*:

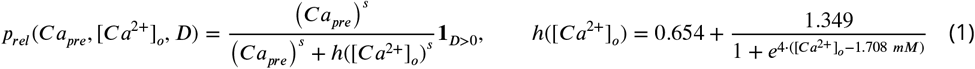

which is a function of presynaptic calcium *Ca_pre_* and extracellular calcium concentration [Ca^2+^]_o_ through the threshold *h*([*Ca*^2+^]_*0*_). To decide whether a vesicle is released for a presynaptic spike *t_i_,* we use a phenomenological model of *Ca_pre_* (see ***Figure 8a***) based on a resource-use function (***Tsodyks and Markram, 1997***):

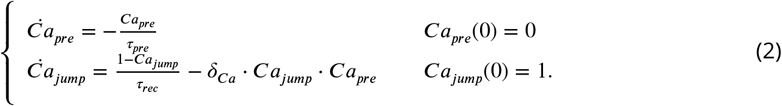

**Table 2.** Stochastic transitions used in the pool dynamics. Note that the rates depend on the pool’s cardinal (Pyle et al., 2000).

| Transition | Rate | Initial Condition |
| --- | --- | --- |
| $(R, D) \rightarrow (R - 1, D + 1)$ | $(D_0 - D) \cdot R / \tau_D$ | $D(0) = D_0$ |
| $(R, D) \rightarrow (R + 1, D - 1)$ | $(R_0 - R) \cdot D / \tau_R$ | $R(0) = R_0$ |
| $(R, D) \rightarrow (R + 1, D)$ | $(R_0 - R) / \tau_R^{ref}$ | |

Upon arrival of the presynaptic spikes, *t* ∈(*t*_1_,..., *t_n_*), we update *Ca_pre_* according to the deterministic jump:

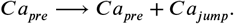

Finally, after *Ca_pre_* has been updated, a vesicle is released with probability *p_rel_* (***Figure 8b***).

Parameters for the vesicle release model are given in ***Table 3***. The experimental constraints to devise a release probability model are given by ***Hardingham et al. (2006)*** and ***Tigaret et al. (2016)***. Because [Ca^2+^]_o_ modifies the release probability dynamics (***King et al., 2001***), we fixed an initial release probability to 68 % for [Ca^2+^]_o_ = 2.5 mM as reported by ***Tigaret et al. (2016)*** (initial value in ***Figure 8b,d***). Additionally, ***Hardingham et al. (2006)*** reports a 38% reduction in the initial release probability when changing [Ca^2+^]_o_ from 2.5 mM to 1 mM. Taking these into account, the decreasing sigmoid function in the Figure 8e depicts our [Ca^2+^]_o_-dependent release probability model (*p_rel_*).

**Table 3.** Parameter values used in the presynaptic model. Our model does not implement a larger pool called “resting pool” containing ~ 180 vesicles (CA3-CA1 hippocampus) (***Alabi and Tsien, 2012***). **Terminology note**: In other works, the larger pool with ~ 180 vesicles can be found with different nomenclatures such as “reserve pool” (***Südhof, 2000***) or “resting pool” (***Alabi and Tsien, 2012***). Furthemore, the nomenclature used in our model for the reserve pool is use in other studies as the “recycling pool”, e.g. ***Rizzoli and Betz (2005)*** and ***Alabi and Tsien (2012)***.

| Name | Value | Reference |
| --- | --- | --- |
| **Vesicle release model (stochastic part)** |  |  |
| initial number of vesicles at D | $D_0 = 25$ | 5 to 20 (Rizzoli and Betz, 2005; Alabi and Tsien, 2012) |
| initial number of vesicles at R | $R_0 = 30$ | 17 to 20 vesicles (Alabi and Tsien, 2012) |
| time constant $R \rightarrow D$<br>(D recycling) | $\tau_D = 5 \text{ s}$ | 1 s (Rizzoli and Betz, 2005) |
| time constant $D \rightarrow R$<br>(R mixing) | $\tau_R = 45 \text{ s}$ | 20 s (when depleted) to 5 min (hypertonic shock)<br>(Rizzoli and Betz, 2005; Pyle et al., 2000) |
| time constant $1 \rightarrow R$<br>(R recycling) | $\tau_R^{ref} = 40 \text{ s}$ | 20 to 30 s (Rizzoli and Betz, 2005) |
| release probability half-activation curve | $h$ | see Equation 1 |
| release probability sigmoid slope | $s = 2$ | fixed for all $[\text{Ca}^{2+}]_o$ |
| **Vesicle release model (deterministic part)** |  |  |
| $Ca_{pre}$ attenuation recovery | $\tau_{pre} = 20 \text{ ms}$ | 50 - 500 ms with dye (Maravall et al., 2000)<br>therefore < 50 to 500 ms without dye |
| deterministic jump attenuation recovery | $\tau_{rec} = 20 \text{ s}$ | $\sim 20 \text{ s}$ (Rizzoli and Betz, 2005) |
| deterministic jump attenuation fraction | $\delta_{ca} = 0.0004$ | (Forsythe et al., 1998) |

Figure 8e shows that our *p_rel_* function is in good agreement with a previous analytical model suggesting that *p_rel_*([Ca^1+^]_*0*_) < ([*Ca*^2+^]_o_)^2^ *mM*^-2^ (***King et al., 2001***). Our model also qualitatively reproduces the vanishing of calcium dye fluorescence levels after 20 s of theta trains from ***Tigaret et al. (2016)*** (in their Supplementary Materials). We interpret their fluorescence measurements as an effect of short-term depression (see ***Figure 8b***).

Despite our model agreeing with previous works, it is a simplified presynaptic model that does not encompass the highly heterogeneous nature of vesicle release. Vesicle release dynamics are known to be sensitivity to various experimental conditions such as temperature (***Fernández-Alfonso and Ryan, 2004***), the age for some brain regions (***Rudolph et al., 2015***) or magnesium concentration (***Hardingham et al., 2006***). Furthermore, since our model of vesicle dynamics is simple, *τ_rec_* in Equation 2 has two roles: to delay the *p_rel_* recovery caused by *Ca_pre_* inactivation (enforced by *δ_Ca_* in ***Equation 2***) and to prevent vesicle release after HFS-induced depression (***King et al., 2001; Rizzoli and Betz, 2005***). Later, we incorporate a higher number of experimental parameters (age, temperature, [Ca^2+^]_o_, [Mg^2+^]_o_) with our NMDAr model, the main postsynaptic calcium source.

### Model compartments

Our model is built over three compartments, a spherical dendritic spine linked by the neck to a cylindrical dendrite connected to a spherical soma. The membrane potential of these compartments satisfy the equations below (parameters in ***Table 4***). Since the dendrite is a single compartment, the precise spine location is undefined. For more detailed morphological simulations to predict plasticity see ***Ebner et al. (2019), Chindemi et al. (2020)*** and ***Jędrzejewska-Szmek et al. (2017)***. The distance from the soma to the spine functionally mimics the BaP attenuation as shown in ***Golding et al. (2001)***, and it is set to 200 μm for all simulations, except in ***Figure 3-Figure Supplement 6c*** and ***Figure 3-Figure Supplement 5e***. In these panels, we modified this distance as described in the graph y-axis to model ***Ebner et al. (2019)*** data. The different currents in the soma, dendrite and spine are described as follows.

**Table 4.** Parameters for the neuron electrical properties. * The membrane leak conductance in the spine is small since the spine resistance is so high that is considered infinite (*>* 10^6^*M*Ω) (*Koch andZador, 1993).* The current thus mostly leaks axially through the neck cytoplasm. The dendrite leak conductance is also small in order to control the distance-dependent attenuation by the axial resistance term 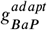 in Equation 4 and ***Equation 5***.

| Name | Value | Reference |
| --- | --- | --- |
| **Passive cable** |  |  |
| leak reversal potential | $E_{leak} = -70 mV$ | 69mV (Spigelman et al., 1996) |
| membrane leak conductance (for spine and passive dendrite) | $g_{leak} = 4 \cdot 10^{-6} nS/\mu m^2$ | * see table legend (Koch and Zador, 1993) |
| membrane leak conductance (only soma) | $g_{soma} = 5.31 \cdot 10^{-3} nS/\mu m^2$ | $3 \cdot 10^{-4}$ to $1.3 \cdot 10^{-3} nS/\mu m^2$ (Fernandez and White, 2010)<br>47 to $2.1 \cdot 10^3 nS$ (NeuroElectro:CA1) |
| membrane capacitance | $C_m = 6 \cdot 10^{-3} pF/\mu m^2$ | $1 \cdot 10^{-2} pF/\mu m^2$ (Hines and Carnevale, 1997)<br>17 to 177 pF (NeuroElectro:CA1) |
| axial resistivity of cytoplasm | $R_a = 1 \cdot 10^{-2} G\Omega\mu m$ | $2 \cdot 10^{-3} G\Omega\mu m$ (Golding et al., 2001) |
| **Dendrite** |  |  |
| dendrite diameter | $D_{dend} = 2 \mu m$ | same as Yi et al. (2017) |
| dendrite length | $L_{dend} = 1400 \mu m$ | apical dendrites, 1200 to 1600 $\mu m$ (Mendoza et al., 2018) |
| dendrite surface area | $A_{dend} = 8.79 \cdot 10^3 \mu m^2$ | $\pi \cdot D_{dend} \cdot L_{dend}$ |
| dendrite volume | $Vol_{dend} = 4.4 \cdot 10^3 \mu m^3$ | $\pi \cdot (D_{dend}/2)^2 \cdot L_{dend}$ |
| dendritic membrane capacitance | $C_{dend} = 52.77 pF$ | $C_m \cdot A_{dend}$ |
| dendrite leak reversal potential | $g_{leakdend} = 3.51 \cdot 10^{-2} nS$ | $g_{leak} \cdot A_{dend}$ |
| dendrite axial conductance | $g_{diff} = 50 nS$ | $R_a \cdot A_{dend}$ |
| **Soma** |  |  |
| soma diameter | $D_{soma} = 30 \mu m$ | 21 $\mu m$ (Stuart et al., 2016) page 3 |
| soma area (sphere) | $A_{soma} = 2.82 \cdot 10^3 \mu m^2$ | $(4\pi/3) \cdot (D_{soma}/2)^3$; $2.12 \cdot 10^3 \mu m^2$ (Zhuravleva et al., 1997) |
| soma membrane capacitance | $C_{soma} = 16.96 pF$ | $C_m \cdot A_{soma}$ |
| soma leaking conductance | $g_{leaksoma} = 15 nS$ | $g_{soma} \cdot A_{soma}$ (Fernandez and White, 2010) |
| **Dendritic spine** |  |  |
| spine head volume | $Vol_{sp} = 0.03 \mu m^3$ | Bartol et al. (2015) |
| spine head surface | $A_{sp} = 4.66 \cdot 10^{-1} \mu m^2$ | $4\pi \cdot (3Vol_{sp}/4\pi)^{2/3}$ |
| spine membrane capacitance | $C_{sp} = 2.8 \cdot 10^{-3} pF$ | $C_m \cdot A_{sp}$ |
| spine head leak conductance | $g_{leaksp} = 1.86 \cdot 10^{-6} nS$ | $g_{leak} \cdot A_{sp}$ |
| **Dendritic spine neck** |  |  |
| spine neck diameter | $D_{neck} = 0.1 \mu m$ | 0.05 to 0.6 $\mu m$ (Harris et al., 1992) |
| neck length | $L_{neck} = 0.2 \mu m$ | $0.7 \pm 0.6 \mu m$ (Adrian et al., 2017) |
| neck cross sectional area | $CS_{neck} = 7.85 \cdot 10^{-3} \mu m^2$ | $\pi \cdot (D_{neck}/2)^2$ |
| neck resistance | $g_{neck} = 3.92 nS \approx 255.1 M\Omega$ | $CS_{neck}/(L_{neck} \cdot R_a)$<br>50 to 550 $M\Omega$ ($275 \pm 27 M\Omega$) (Popovic et al., 2015) |

### Membrane potential and currents

The membrane potential of these compartments satisfy the equations below (parameters in ***Table 4***). The different currents are described in the following sections.

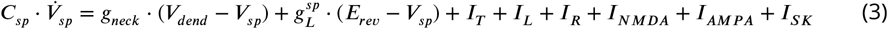

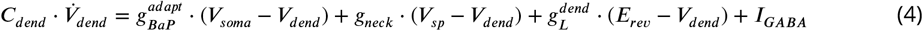

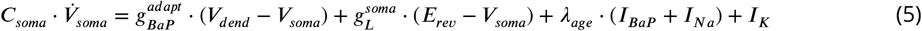

### Action-potential backpropagation (BaP)

#### Postsynaptic currents

The postsynaptic currents are generated in the soma, backpropagated to the dendritic spine and filtered by a passive dendrite. The soma generates BaPs using a version of the Na^+^ and K^+^ channel models developed by ***Migliore et al. (1999)***. The related parameters are described in ***Table 5*** (the voltage unit is mV).

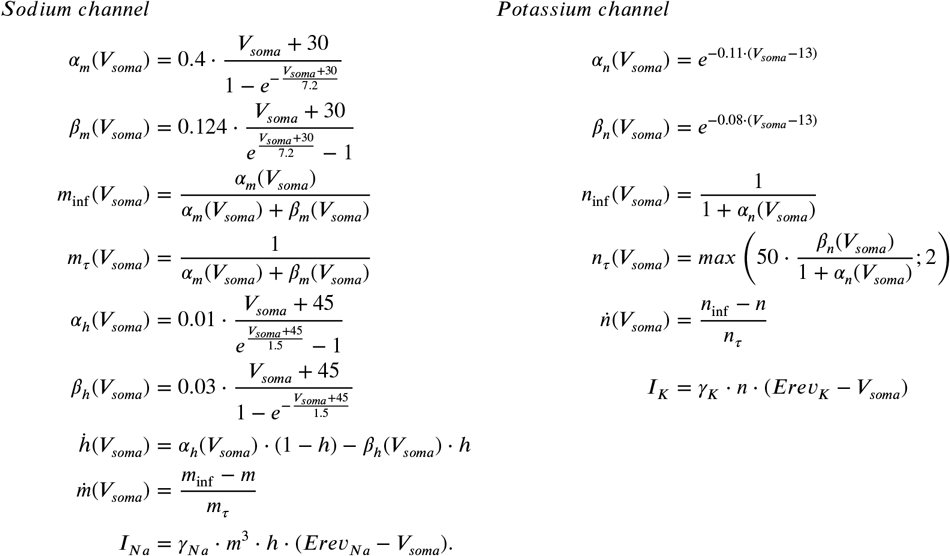

**Table 5.** The Na+ and K+ conductances intentionally do not match the reference because models with passive dendrite need higher current input to initiate action potentials (***Levine and Woody, 1978***). Therefore we set it to achieve the desired amplitude on the dendrite and the dendritic spine according to the predictions of ***Golding et al. (2001)*** and ***Kwon et al. (2017)***.

| Name | Value | Reference |
| --- | --- | --- |
| **Soma parameters for Na<sup>+</sup> and K<sup>+</sup> channel** |  |  |
| sodium conductance | $\gamma_{Na} = 8 \cdot 10^2 \text{ nS}$ | 0.32 nS/$\mu\text{m}^2$ (*Migliore et al., 1999*)<br>see legend commentary |
| potassium conductance | $\gamma_K = 40 \text{ nS}$ | 0.48 nS/$\mu\text{m}^2$ (*Migliore et al., 1999*)<br>see legend commentary |
| reversal potential sodium | $E_{rev_{Na}} = 50 \text{ mV}$ | *Migliore et al. (1999)* |
| reversal potential potassium | $E_{rev_K} = -90 \text{ mV}$ | *Migliore et al. (1999)* |
| **BaP attenuation parameters** |  |  |
| attenuation step factor (age) | $\delta_{age}$ | see *Equation 7* and *Figure 9b,c bottom*<br>*Buchanan and Mellor (2007); Golding et al. (2001)* |
| attenuation step factor | $\delta_{decay} = 1.727 \cdot 10^{-5}$ | adjusted to fit<br>*Buchanan and Mellor (2007); Golding et al. (2001)* |
| auxiliary attenuation step factor | $\delta_{aux} = 2.304 \cdot 10^{-5}$ | adjusted to fit<br>*Buchanan and Mellor (2007); Golding et al. (2001)* |
| recovery time for the attenuation factor | $\tau_{rec} = 2 \text{ s}$ | adjusted to fit<br>*Buchanan and Mellor (2007); Golding et al. (2001)* |
| recovery time for the age attenuation factor | $\tau_{rec}^{age} = 0.5 \text{ s}$ | adjusted to fit<br>*Buchanan and Mellor (2007); Golding et al. (2001)* |
| **AP evoked by EPSP** |  |  |
| decay time for $V_{evoke}$ | $\tau_v = 40 \text{ ms}$ | *Hines and Carnevale (1997)* |
| delay AP evoked by EPSP | $\delta_{delay-AP} = 15 \text{ ms}$ | *Fricker and Miles (2000)* |

To trigger a BaP, an external current *I_BaP_* is injected in the soma at times *t* ∈ {*t*_1_,..., *t_n_* (postsynaptic input times) for a chosen duration *δ_inj_* with amplitude *I_amp_* (*nA*), considering *H* as the Heaviside function this is expressed as:

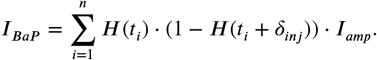

The current injected in the soma is filtered in a distance-dependent manner by the dendrite before it reaches the dendritic spine. Biologically, BaP adaptation is caused by the inactivation of sodium channels and the difference of sodium and potassium channel expression along the dendrite (***Jung et al., 1997; Golding et al., 2001***). We used a phenomenological model, implementing distant-dependent BaP amplitude attenuation by modifying the axial resistance 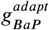 (see ***Equation 4 and Equation 5***) between the dendrite and the soma as follows (***Figure 9c top***):

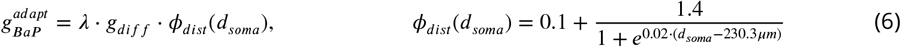

where *d_soma_* is the distance of the spine to the soma and where the factor *λ* is dynamically regulated based on a resource-use equation from ***Tsodyks and Markram (1997)*** with a dampening factor *λ_aux_* changing the size of the attenuation step *δ_decay_*:

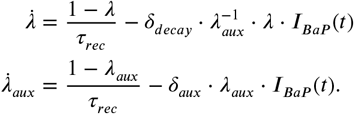

**Figure 9.**
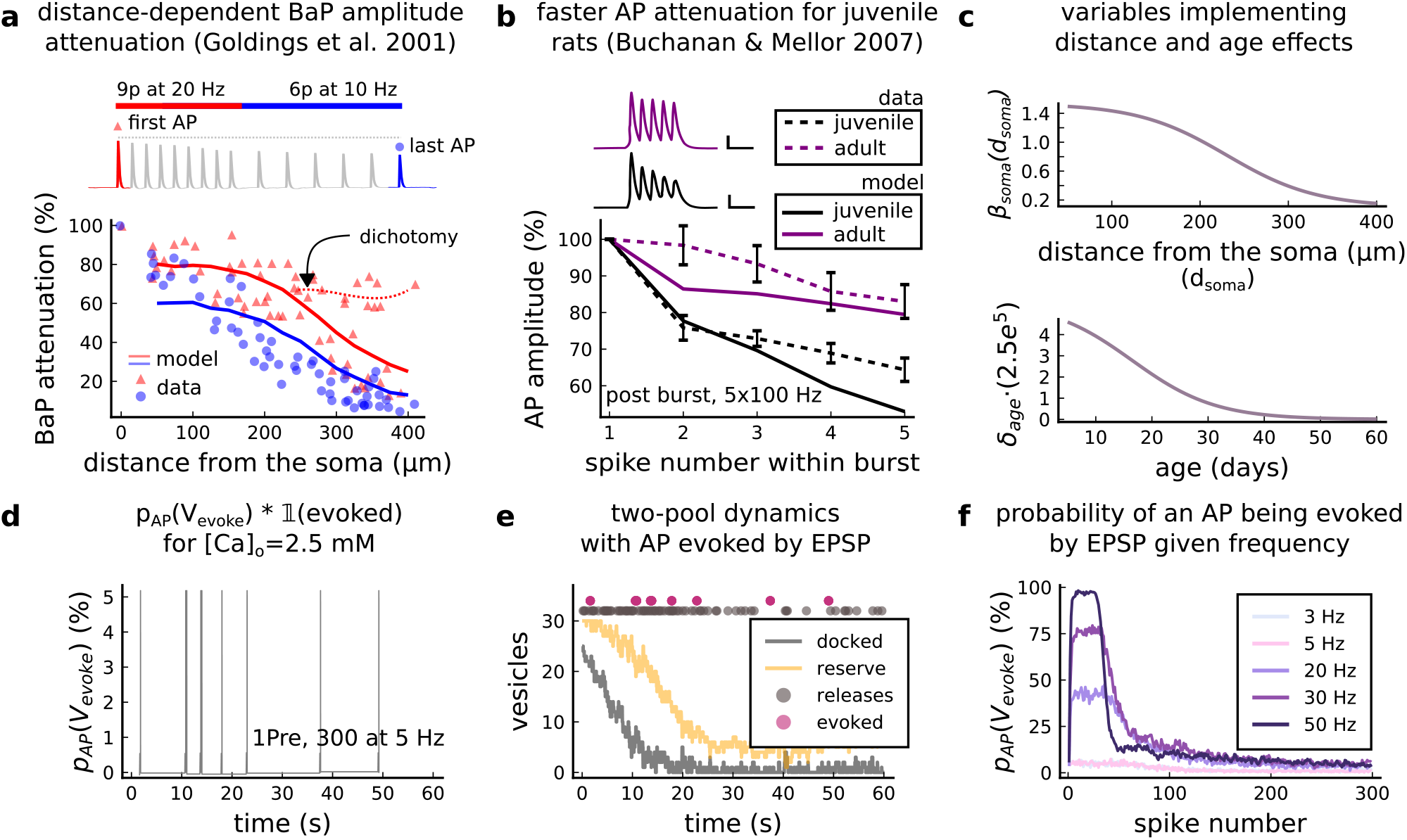
AP Evoked by EPSP. **a**, Model and data comparison for the distance-dependent BaP amplitude attenuation measured in the dendrite and varying the distance from the soma. The stimulation in panel **a** is set to reproduce the same stimulation as ***Golding et al. (2001)***. Golding described two classes of neurons: those that are strongly attenuated and those that are weakly attenuated (dichotomy mark represented by the dashed line). However, in this work we consider only strongly attenuated neurons. **b**, Attenuation of somatic action potential from ***Buchanan and Mellor (2007)*** and model in response to five postsynaptic spikes delivered at 100 Hz. The value showed for the model is the spine voltage with distance from the soma set to zero (scale 25 ms, 20 mV). **c**, Top panel shows the *λ_soma_* used in ***Equation 6*** to modify the axial conductance between the soma and dendrite. Bottom panel shows the age-dependent changes in the step of the resource-use equation (***Equation 7***) that accelerates the BaP attenuation and decreases the sodium currents in Equation ***Equation 5***. **d**, Probability of evoking an AP multiplied by the successfully evoked AP (*p_AP_*(*V_evoked_*) · **l**(*evoked*) for the protocol 1Pre, 300 at 5 Hz (2.5 mM Ca). **e**, Two-pool dynamics with the same stimulation from panel D showing the vesicle release, the reserve and docked pools, and the evoked AP. **f**, Probability of evoking an AP for the protocol 1Pre 300 pulses at different frequencies (3 and 5 Hz have the same probability).

The BaP attenuation model is based on ***Golding et al. (2001)*** data for strongly attenuating neurons. Therefore, the second type of attenuation (weakly attenuating) in neurons is not considered (dichotomy in ***Figure 9a***). ***Figure 9a*** compares Goldings data to our model and illustrates the effect of BaP attenuation in the upper panels of ***Figure 9a,b***.

***Table 5*** shows the BaP attenuation parameters. The plasticity outcomes as function of the dendritic spine distance from the soma are shown in ***Figure 3-Figure Supplement 6c*** and ***Figure 3-Figure Supplement 5e***.

#### Age-dependent BaP adaptation

Age-dependent BaP attenuation modifies the neuronal bursting properties through the maturation and expression of potassium and sodium channels (***Gymnopoulos et al., 2014***), therefore changing the interaction of hyperpolarizing and depolarizing currents (see ***Figure 9b***) (***Grewe et al., 2010; Jung et al., 1997***). We reproduce ***Buchanan and Mellor (2007)*** somatic attenuation profiles (***Figure 9b***) with our model by including an age-dependent BaP amplitude attenuation factor. We define the attenuation factor *λ_age_* (***Figure 9c bottom***), as follows.

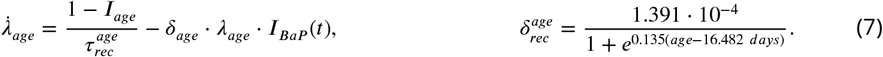

In Equation ***Equation 5***, the age effects are introduced by multiplying the sodium *I_Na_* and the external *I_BaP_* currents by the attenuation factor *λ_age_*.

#### AP evoked by EPSP

A presynaptic stimulation triggers a BaP if sufficient depolarization is caused by the EPSPs reaching the soma (***Stuart et al., 2016***). We included an option to choose whether an EPSP can evoke an AP using an event generator resembling the previous release probability model *p_rel_* as in the ***Equation 1***. Like *p_rel_*, the BaPs evoked by EPSPs are estimated before the postsynaptic simulation. We use a variable *V_evoke_* which is incremented by 1 at each presynaptic time *t* ∈ (*t*_1_,..., *t_n_*) and has exponential decay:

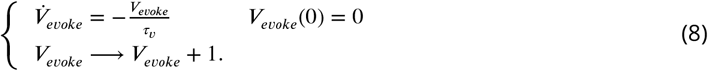

Since the BaPs evoked by EPSPs are triggered by the afferent synapses and are limited by their respective docked pools (*D),* we use the previous *p_rel_* to define the probability of an AP to occur. We test the ratio of successful releases from 25 synapses to decide if a BaP is evoked by an EPSP, setting a test threshold of 80%. Therefore, we express the probability of evoking an AP, *p_AP_*(*V_emke_*), with the following test:

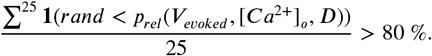

The EPSP summation dynamics on the soma and dendrites depend on the complex neuron morphology (***Etherington et al., 2010; Ebner et al., 2019***) which was not implemented by our model. Therefore, our “AP evoked by EPSP test” intends to give a simplified way to produce BaPs similar to an integrate-and-fire model (***Sterratt et al., 2011***).

Previous work suggests that BaPs can be evoked with a ~5 % probability for low-frequencies in the Dudek and Bear experiment ([Ca^2+^]_o_ = 2.5 mM) (***Mayr and Partzsch, 2010***). Our model covers this estimation, but the chance to elicit an AP increases with the frequency (***Etherington et al., 2010***). This is captured by the *V_evoke_* (in an integrate-and-fire fashion (***Stuart et al., 2016***)) as shown in ***Figure 9f***. The ***Figure 9d,e*** show how a 5 Hz stimulation evokes APs. The delay between the EPSP and the evoked AP is set to *δ_del-AP_* = 15*ms*, similar to the EPSP-spike latency reported for CA1 pyramidal neurons (Fricker and Miles, 2000).

### AMPAr

#### Markov chain

The AMPAr is modeled with the Markov chain (***Figure 10***) described by ***Robert and Howe (2003)*** and ***Coombs et al. (2017)*** and adapted to temperature changes according to ***Postlethwaite et al. (2007)***. Here, we introduce the additional parameters 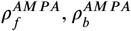 to cover AMPAr temperature-sensitive kinetics (***Postlethwaite et al., 2007***). The corresponding parameters are given in ***Table 6***.

**Figure 10.**
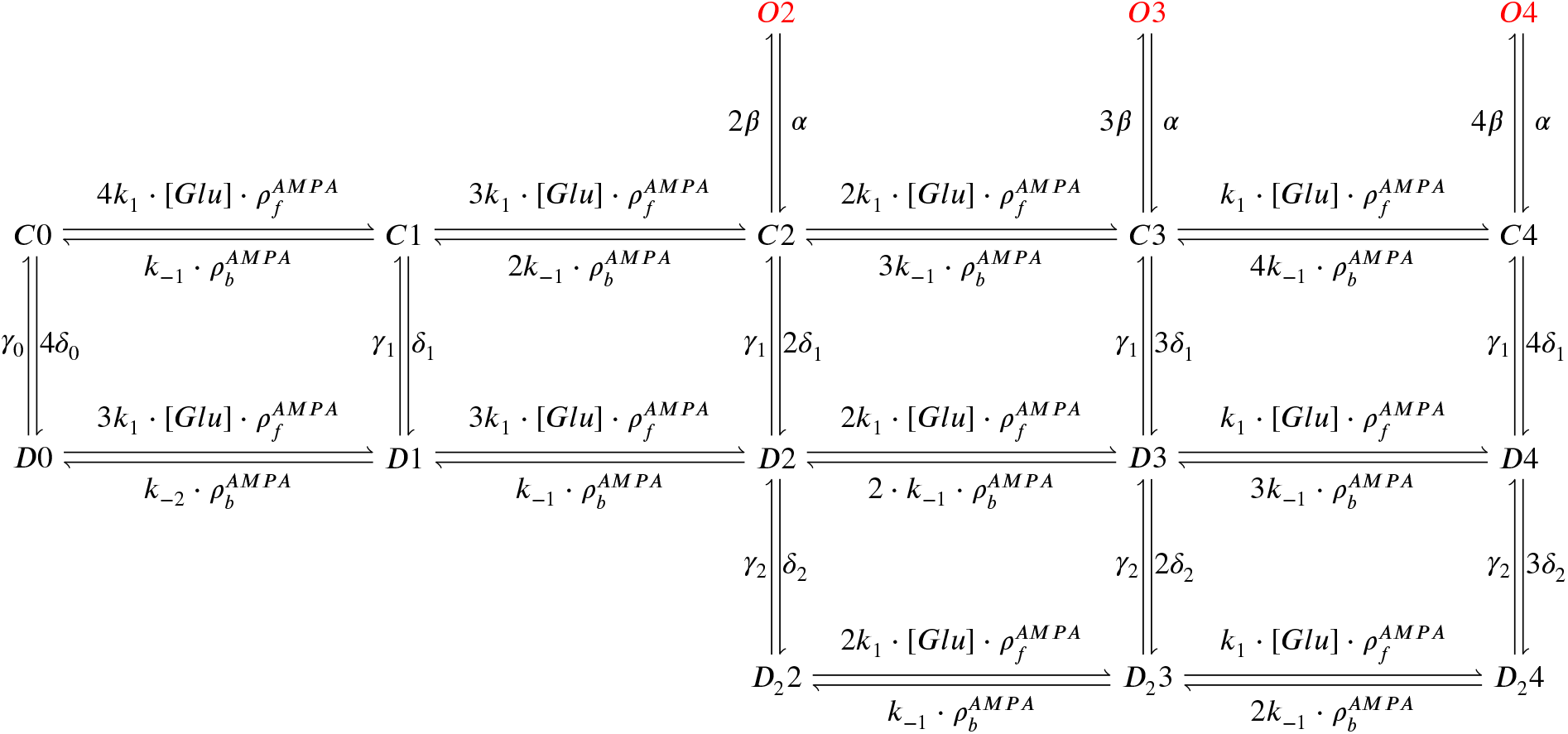
AMPAr Markov chain with three sub-conductance states and two desensitisation levels. It includes parameters 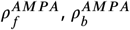 (binding and unbinding of glutamate) which depend on temperature. Open states are O2, O3 and O4; closed states are C0, C1, C2, C3 and C4; desensitisation states are D0, D1, D2, D3 and D4; deep desensitisation states are D_2_2, D_2_3 and D_2_4.

**Table 6.** Parameter values for the AMPAr Markov chain and glutamate release affecting NMDAr, AMPAr. Properties of GABA release are the same as those for glutamate.

| Name | Value | Reference |
| --- | --- | --- |
| **Glutamate parameters** |  |  |
| duration of glutamate in the cleft | $glu_{width} = 1 \text{ ms}$ | *Spruston et al. (1995)* |
| concentration of glutamate in the cleft | $glu_{amp} = 1 \text{ mM}$ | *Spruston et al. (1995)* |
| glutamate variability<br>(gamma distribution $\Gamma$) | $glu_{cv} = \Gamma(1/0.5^2, 0.5^2)$ | *Liu et al. (1999)* |
| glutamate signal | $Glu$ | $glu_{cv} \cdot glu_{amp}$<br>for AMPAR, NMDAR and copied to GABA neurotransmitter |
| **AMPA parameters** |  |  |
| number of AMPARs | $N_{AMPA} = 120$ | *Bartol et al. (2015)* |
| reversal potential | $E_{rev_{AMPA}} = 0 \text{ mV}$ | *Bartol et al. (2015)* |
| subconductance O2 | $\gamma_{A2} = 15.5 \text{ pS}$ | 16.3 pS (*Coombs et al., 2017*) |
| subconductance O3 | $\gamma_{A3} = 26 \text{ pS}$ | 28.7 pS (*Coombs et al., 2017*) |
| subconductance O4 | $\gamma_{A4} = 36.5 \text{ pS}$ | 37.8 pS (*Coombs et al., 2017*) |
| glu binding | $k_1 = 1.6 \cdot 10^7 \text{ M}^{-1} \text{ s}^{-1}$ | *Robert and Howe (2003)* |
| glu unbinding 1 | $k_{-1} = 7400 \text{ s}^{-1}$ | *Robert and Howe (2003)* |
| glu unbinding 2 | $k_{-2} = 0.41 \text{ s}^{-1}$ | *Robert and Howe (2003)* |
| closing | $\alpha = 2600 \text{ s}^{-1}$ | *Robert and Howe (2003)* |
| opening | $\beta = 9600 \text{ s}^{-1}$ | *Robert and Howe (2003)* |
| desensitisation 1 | $\delta_1 = 1500 \text{ s}^{-1}$ | *Robert and Howe (2003)* |
| desensitisation 2 | $\delta_2 = 170 \text{ s}^{-1}$ | *Robert and Howe (2003)* |
| desensitisation 3 | $\delta_0 = 0.003 \text{ s}^{-1}$ | *Robert and Howe (2003)* |
| re-desensitisation 1 | $\gamma_1 = 9.1 \text{ s}^{-1}$ | *Robert and Howe (2003)* |
| re-desensitisation 2 | $\gamma_2 = 42 \text{ s}^{-1}$ | *Robert and Howe (2003)* |
| re-desensitisation 3 | $\gamma_0 = 0.83 \text{ s}^{-1}$ | *Robert and Howe (2003)* |

The AMPAr current is the sum of the subcurrents associated to the occupancy of the three subconductance states O2, O3 and O4 of the Markov chain in Figure 10 and described as follows:

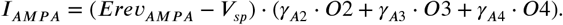

The adaptation of the Markov chain from ***Robert and Howe (2003)*** is made by changing the forward 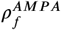 and backward 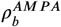 rates in a temperature-dependent manner matching the decay time reported by ***Postlethwaite et al. (2007)***:

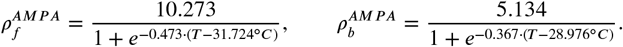

The effects of temperature change on AMPAr dynamics are presented in ***Figure 11***, which also shows that the desensitisation is not altered by temperature changes (***Figure 11b,c***). The recovery time from desensitisation is the same as at room temperature (***Robert and Howe, 2003***). Desensitisation measurements are required to account for a temperature-dependent change in the rates of the “vertical” transitions in ***Figure 10***, see ***Postlethwaite et al. (2007)***. This can be relevant for presynaptic bursts.

**Figure 11.**
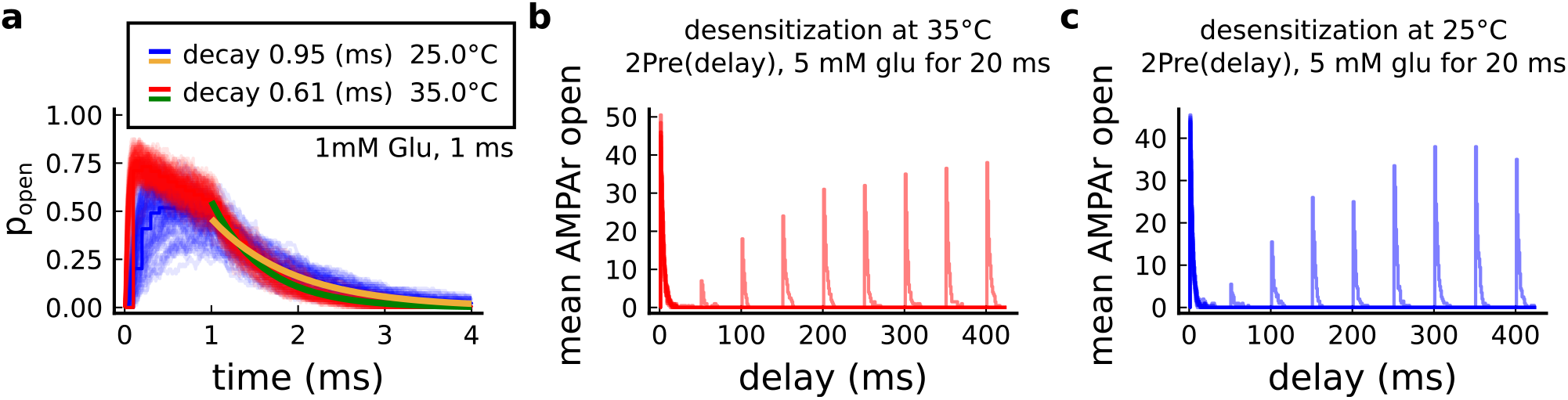
Effect of temperature in the AMPAr. **a**, Probability of AMPAr opening 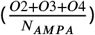 and the decay time at different temperatures in response to 1 mM glutamate during 1ms (standard pulse). ***Postlethwaite et al. (2007)*** data (our model) suggests that AMPAr decay time at 35°*C* is ~ 0.5 *ms* (~ 0.6 *ms*) and at 25°*C* is ~ 0.65 *ms* (~ 0.95 *ms*). This shows a closer match towards more physiological temperatures. **b**, Desensitisation profile of AMPAr at 35°*C* showing how many AMPAr are open in response to a glutamate saturating pulse (5 mM Glu during 20 ms) separated by an interval (x-axis). **c**, Same as in panel **b** but for 25°*C*.

#### Postsynaptic Ca^2+^ influx

The effects of experimental conditions on the calcium dynamics are due to receptors, ion channels and enzymes. A leaky term models the calcium resting concentration in the ***Equation 9***. The calcium fluxes from NMDAr and VGCCs (T, R, L types) are given in ***Equation 10***. The diffusion term through the spine neck is expressed in ***Equation 11.*** Finally, the buffer, the optional dye and the enzymatic reactions are given in ***Equation 12*** (parameter values given at the **Table 7**):

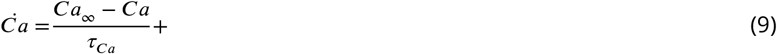

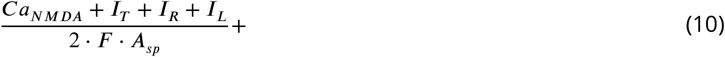

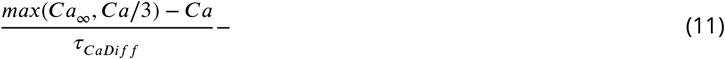

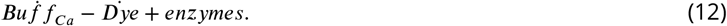

**Table 7.** Postsynaptic calcium dynamics parameters.

| Name | Value | Reference |
| --- | --- | --- |
| **Buffer and dye** |  |  |
| association buffer constant | $k_{on}^{Buf f} = 0.247 \mu M^{-1} ms^{-1}$ | *Bartol et al. (2015)* |
| dissociation buffer constant | $k_{off}^{Buf f} = 0.524 ms^{-1}$ | *Bartol et al. (2015)* |
| buffer concentration | $Buf f_{con} = 62 \mu M$ | $76.7 \mu M$ (*Bartol et al., 2015*) |
| **Calcium dynamics** |  |  |
| Calcium baseline concentration | $Ca_{\infty} = 50 \text{ nM}$ | $37 \pm 5$ to $54 \pm 5 \text{ nM}$ (*Maravall et al., 2000*) |
| Calcium decay time | $\tau_{Ca} = 10 \text{ ms}$ | 50 to 500 ms for with dye (*Maravall et al., 2000*)<br>therefore < 50 to 500 ms undyed (unbuffered) |
| Calcium diffusion | $D_{Ca} = 0.3338 \mu m^2 ms^{-1}$ | 0.22 to $0.4 \mu m^2 ms^{-1}$ (*Bartol et al., 2015*; *Holcman et al., 2005*) |
| Calcium diffusion time constant | $\tau_{CaDiff} = \frac{V_{ol_{sp}}}{2D_{Ca}^2 \cdot D_{neck}} + \frac{l_{neck}^2}{2D_{Ca}} = 0.5 \text{ ms}$ | 8 ms for a $V_{sp} = 0.7 \mu m^3$ (*Holcman et al., 2005*) |
| **GHK equation** |  |  |
| temperature | $T = 35^{\circ}C$ | converted to Kelvin in the **Equation 14** given the protocol |
| faraday constant | $F = 96.485 \text{ C mol}^{-1}$ | *Hille (1978)* |
| gas constant | $R = 8.314 \text{ J K}^{-1} \text{ mol}^{-1}$ | *Hille (1978)* |
| Calcium permeability | $P_{Ca} = 0.045 \mu m \text{ ms}^{-1}$ | adjusted to produce $3 \mu M$ Calcium in response to a Glu release<br>supplementary files from *Chang et al. (2017)* |
| Calcium ion valence | $z_{Ca} = 2$ | *Hille (1978)* |

Despite the driving force to the resting concentration, *Ca*_∞_ = 50 *nM*, the tonic opening of T-type channels causes calcium to fluctuate making its mean value dependent on temperature, extracellular calcium and voltage. The effects of this tonic opening in various experimental conditions are shown in ***Figure 6-Figure Supplement 2c***. To avoid modelling dendritic calcium sources, we use a dampening term as one-third of the calcium level since calcium imaging comparing dendrite and spine fluorescence have shown this trend (***Segal and Korkotian, 2014***). ***Equation 11*** implements the diffusion of calcium from the spine to the dendrite through the neck. The time constant for the diffusion coefficient *τ_CaDiff_*, is estimated as described in ***Holcman et al. (2005)***. The calcium buffer and the optional dye are described as a two-state reaction system (***Sabatini et al., 2002***):

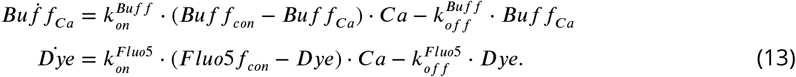

Unlike other calcium-based plasticity models (***Graupner and Brunel, 2012***) using the dye fluorescence decay as an approximation to calcium decay, our model is based on receptor and ion channel kinetics. Additionally, our model can simulate the dye kinetics as a buffer using ***Equation 13***) when appropriate. See ***Figure 12*** that highlights differences between calcium and dye dynamics which is affected by the laser-induced temperature increase (***Wells et al., 2007; Deng et al., 2014***). We estimated the calcium reversal potential for the calcium fluxes using the Goldman–Hodgkin–Katz (GHK) flux equation described in ***Hille (1978)***. The calcium ion permeability, *P_Ca_*, was used as a free parameter adjusting a single EPSP to produce a calcium amplitude of ~ 3 μM (***Chang et al., 2017***).

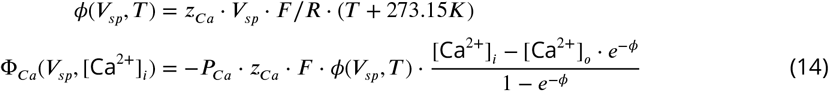

**Figure 12.**
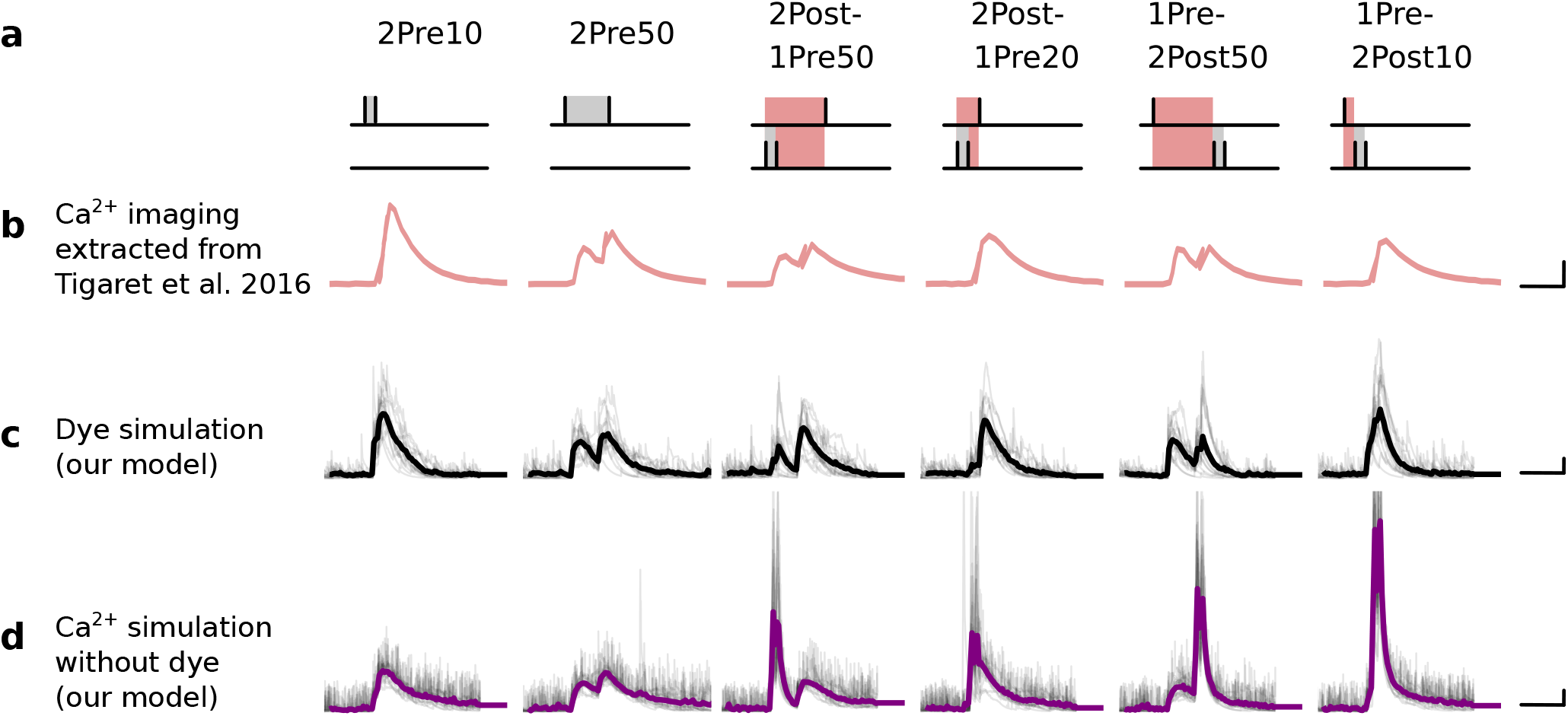
Differences between dye measurements and simulated calcium. **a,** Pre and postsynaptic stimuli as used in ***Tigaret et al. (2016)***. **b,** Calcium imaging curves (fluorescence ΔF/A) elicited using the respective stimulation protocols above with Fluo5 200 μM (extracted from ***Tigaret et al. (2016)***). Scale 100 ms, 0.05 ΔF/A. **c,** Dye simulation using the model. The dye is implemented by increasing temperature to mimic laser effect on channel kinetics and decreases the interaction between NMDAr and voltage elicited by BaP. Temperature effects over NMDAr are shown in ***Korinek et al. (2010)***. Also, the effects of temperature on calcium-sensitive probes shown in ***Oliveira et al. (2012)*** (baseline only, likely related to T-type channels). Other examples of laser heating of neuronal tissue are given in ***Deng et al. (2014)***. Such a dye curve fitting was obtained by increasing temperature by 10°*C* to mimic laser-induced heating (***Wells et al., 2007; Deng et al., 2014***). We achieved a better fit by decreasing the amplitude of the BaP that reaches the dendrite. Additionally, for fitting purposes, we assumed that a temperature increase lead to a decrease in BaP amplitude. Scale 0.6 μM dye, 100 ms. **d,** Calcium simulation without dye. Scale 0.85 μM Ca^2+^, 100 ms.

*Φ_Ca_*(*V_sp_*, [Ca^2+^]**i**.) (***Equation 14***) is used to determine the calcium influx through NMDAr and VGCC in the ***Equation 15, Equation 16, Equation 17*** and ***Equation 18*** using the spine membrane voltage and calcium internal concentration ([Ca^2+^]_*i*_). Note that for simplicity the calcium external concentration ([Ca^2+^]_*0*_) was kept fixed during the simulation and only altered by experimental conditions given by the aCSF composition.

### NMDAr – GluN2A and GluN2B

#### Markov chain

In hippocampus, NMDArs are principally heteromers composed of the obligatory subunit GluN1 and either the GluN2A or GluN2B subunits. These N2 subunits guide the activation kinetics of these receptors with the GluN1/GLUN2B heteromers displaying slow kinetics (~ 250ms) and the GluN1/GluN2A heteromers displaying faster kinetics (~ 50ms). We modeled both NMDA subtypes. The NMDAr containing GluN2A is modeled with the following Markov chain (***Popescu et al., 2004***) where we introduce the additional parameters 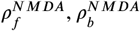:

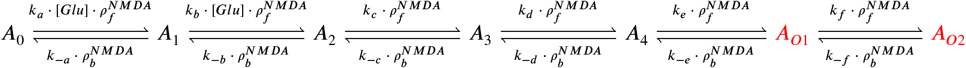

The NMDAr containing GluN2B is modeled with a Markov chain based on the above GluN2A scheme. We decreased the rates by ~75% in order to match the GluN2B decay at 25°*C* as published in ***Iacobucci and Popescu (2018)***.

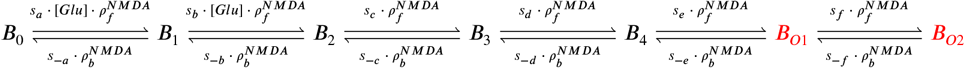

The different rates are given in ***Table 8***.

**Table 8.**
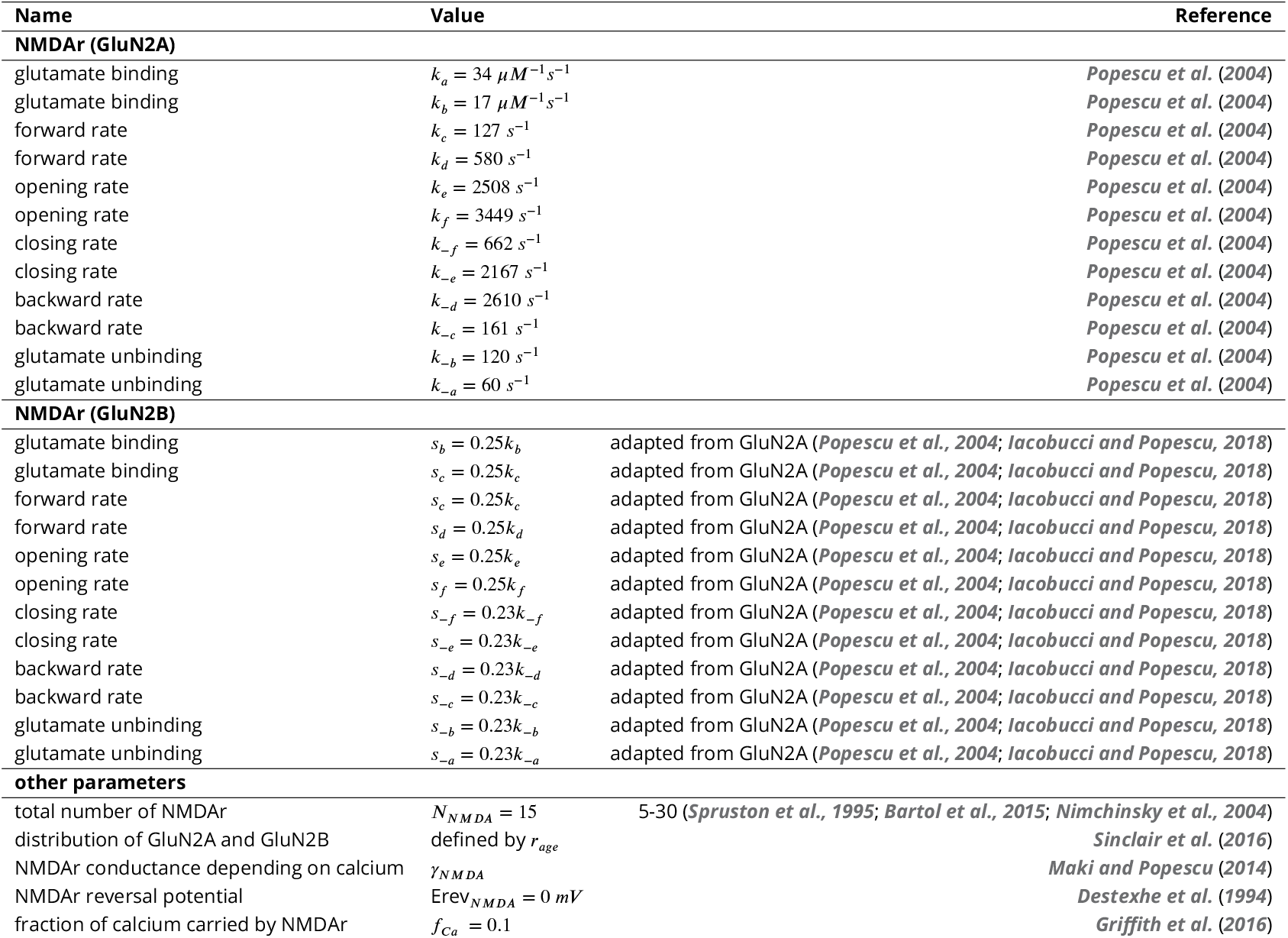
NMDAr parameters.

#### NMDAr and age switch

The age-dependent expression ratio of the subtypes GluN2A and GluN2B (*r_age_*) was obtained from experimental data of mouse hippocampus (***Sinclair et al., 2016***). We added noise to this ratio causing ~1 NMDAr subunit to flip towards GluN2A or GluN2B (see ***Figure 13e***). The population of 15 NMDAr is divided in the two subtypes according to the ratio plotted in ***Figure 13b*** as a function of age. The ratio to define the number NMDAr subtypes as function of age reads:

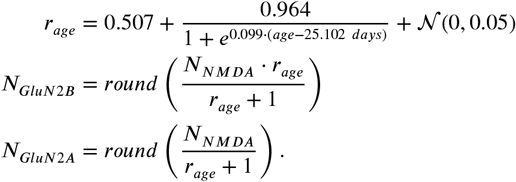

**Figure 13.**
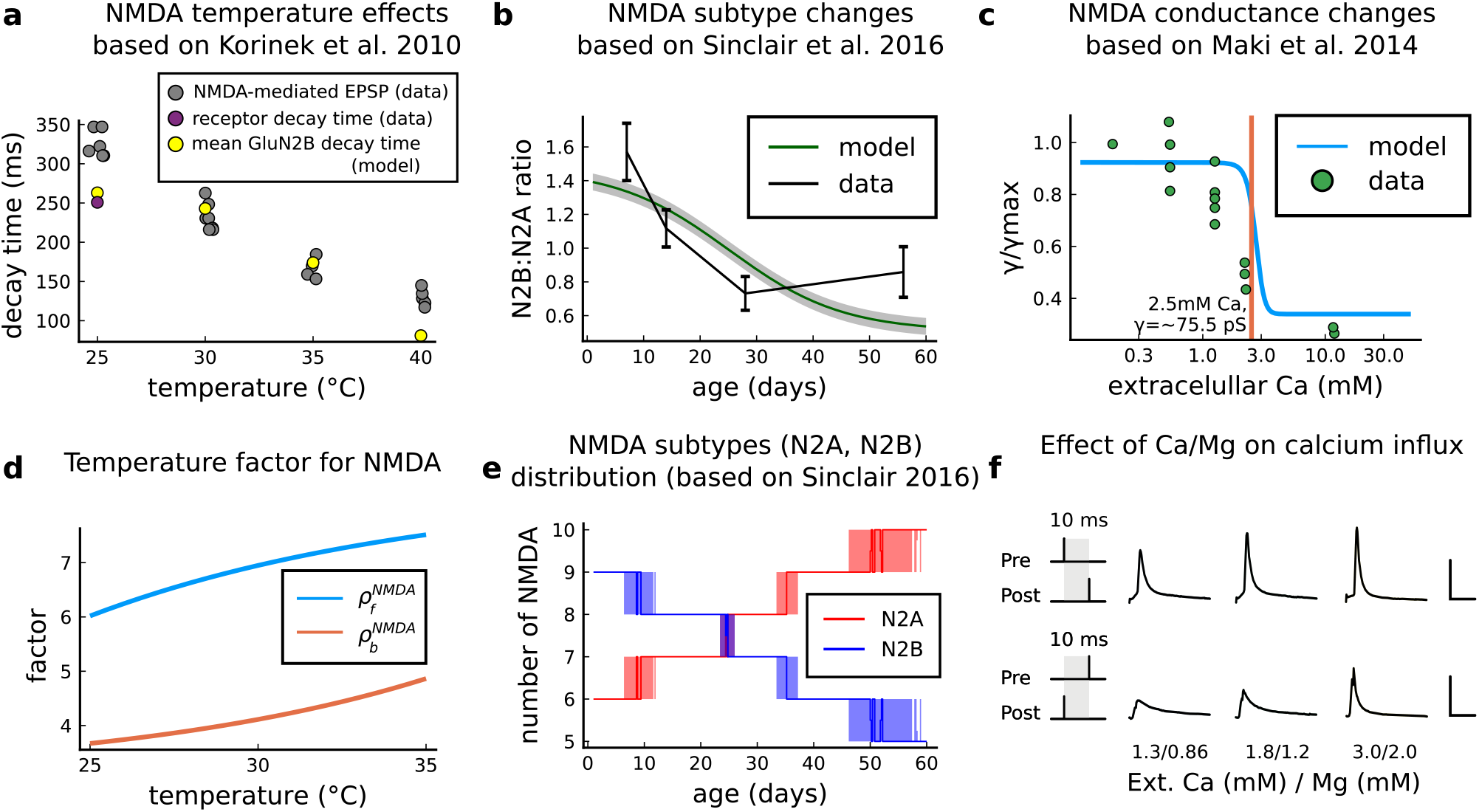
NMDAr changes caused by age, temperature and extracellular and magnesium concentrations in the aCSF. **a** Decay time of the NMDAr-mediated EPSP recorded from neocortical layer II/III pyramidal neurons (grey) (*Korinek et al., 2010)* compared to the decay time from the GluN2B channel estimated by our model (yellow) and data from Iacobussi’s single receptor recording (purple) (***Iacobucci and Popescu, 2018***). **b**, Comparison of our implementation of GluN2B:GluN2A ratio and the GluN2B:GluN2A ratio from the mouse CA1 excitatory neurons. **c**, Comparison of our implementation of NMDAr conductance change in response to the extracellular against data (***Maki and Popescu, 2014***). **d**, Forward and backwards temperature factors implemented to approximate NMDAr subtypes decay times at room temperature (***Iacobucci and Popescu, 2018***) and temperature changes observed in ***Korinek et al. (2010)***. **e**, NMDAr subtype fluctuations in our model with age. We added noise to have a smoother transition between different ages. **f**, Calcium concentration changes for causal and anticausal protocols in response to different aCSF calcium and magnesium compositions with fixed Ca/Mg ratio (1.5). Scale 50 ms and 5 *μM*.

The round term in the two previous equations ensures that we have an integer value for the NMDAr subtypes, making the stair shaped curve seen in ***Figure 13e***.

#### NMDAr and temperature

We adjusted the GluN2A and GluN2B forward and backward rates to follow the temperature effects on NMDAr-mediated EPSP (***Korinek et al., 2010***), see ***Figure 13a,d***. Because GluN2B dominates the NMDAr-mediated EPSP, we fit its decay time of the NMDAr-mediated EPSP as function of temperature as reported by ***Korinek et al. (2010)*** using logistic functions 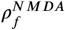 and 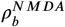. The decay time comparison is shown in ***Figure 13a***. Then, we applied the same temperature factor 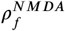 and 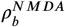 for GluN2A. The decay times of GluN2A and GluN2B are similar to those reported by ***Iacobucci and Popescu (2018)***. The forward and backward factors are described as follows:

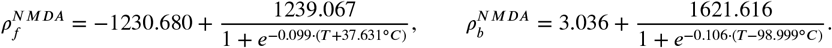

#### NMDAr current and Ca^2+^-dependent conductance

NMDAr conductance is modulated by external calcium and is modelled according to the next equations using NMDAr subconductances *A*_*O*1_ and *A*_*O*2_ (GluN2A), and *B*_*O*1_ and *B*_*O*2_ (GluN2B).

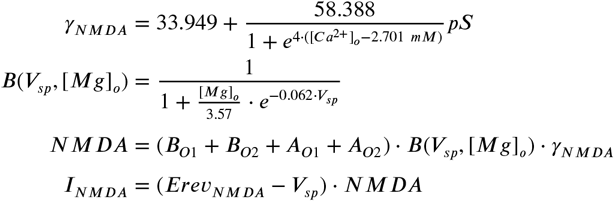

We modified the conductance *γ_NMDA_* as a funtion of extracellular calcium from that reported by ***Maki and Popescu (2014)***. The reported NMDAr conductance at [Ca^2+^]_o_ = 1.8 mM is 53 ± 5*pS*. Here, we used the higher conductance 91.3 *pS* for NMDAr (for both subtypes) at [Ca^2+^]_o_ = 1.8 mM to compensate for the small number of NMDArs reported by ***Nimchinsky et al. (2004)***. Hence, we adjusted Maki and Popescu (2014) data to take into account this constraint: this caused a right-shift in the NMDA-conductance curve (***Figure 13c***). The calcium influx *Ca_NMDA_* is modulated by the GHK factor, ***Equation 14***, as a function of the internal and external calcium concentrations and the spine voltage:

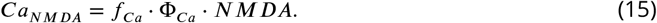

The combined effect of extracellular Magnesium (***Jahr and Stevens, 1990***) and Calcium concentration are displayed in ***Figure 13f***.

#### GABA(A) receptor

Since the precise delay of GABA release relative to glutamate is not known, we assumed GABA and glutamate release are synchronized for simplicity (see **Table 6**). We used the GABA(A) receptor Markov chain (***Figure 14***) presented in ***Busch and Sakmann (1990); Destexhe et al. (1998)*** and we estimated temperature adaptations using the measurements reported by ***Otis and Mody (1992)***.

**Figure 14.**
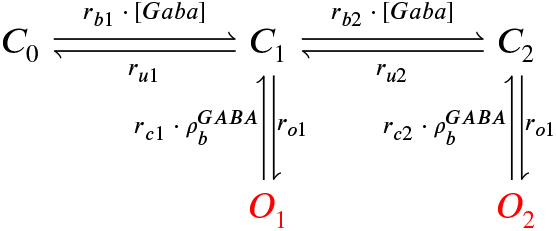
GABAr Markov chain model. Closed states (*C*_0_, *C*_1_ and *C*_2_) open in response to GABAr and can go either close again or open (*O*_1_ and *O*_2_)

#### GABA(A)r and temperature

Because the amplitude of GABA(A) current is altered by the GABAr shift during development (***Rinetti-Vargas et al., 2017***), we applied temperature changes only to the closing rates using a logistic function for 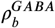, estimated by fitting to the measurements from ***Otis and Mody (1992)*** (data comparison in the ***Figure 15b,e)***.

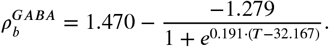

**Figure 15.**
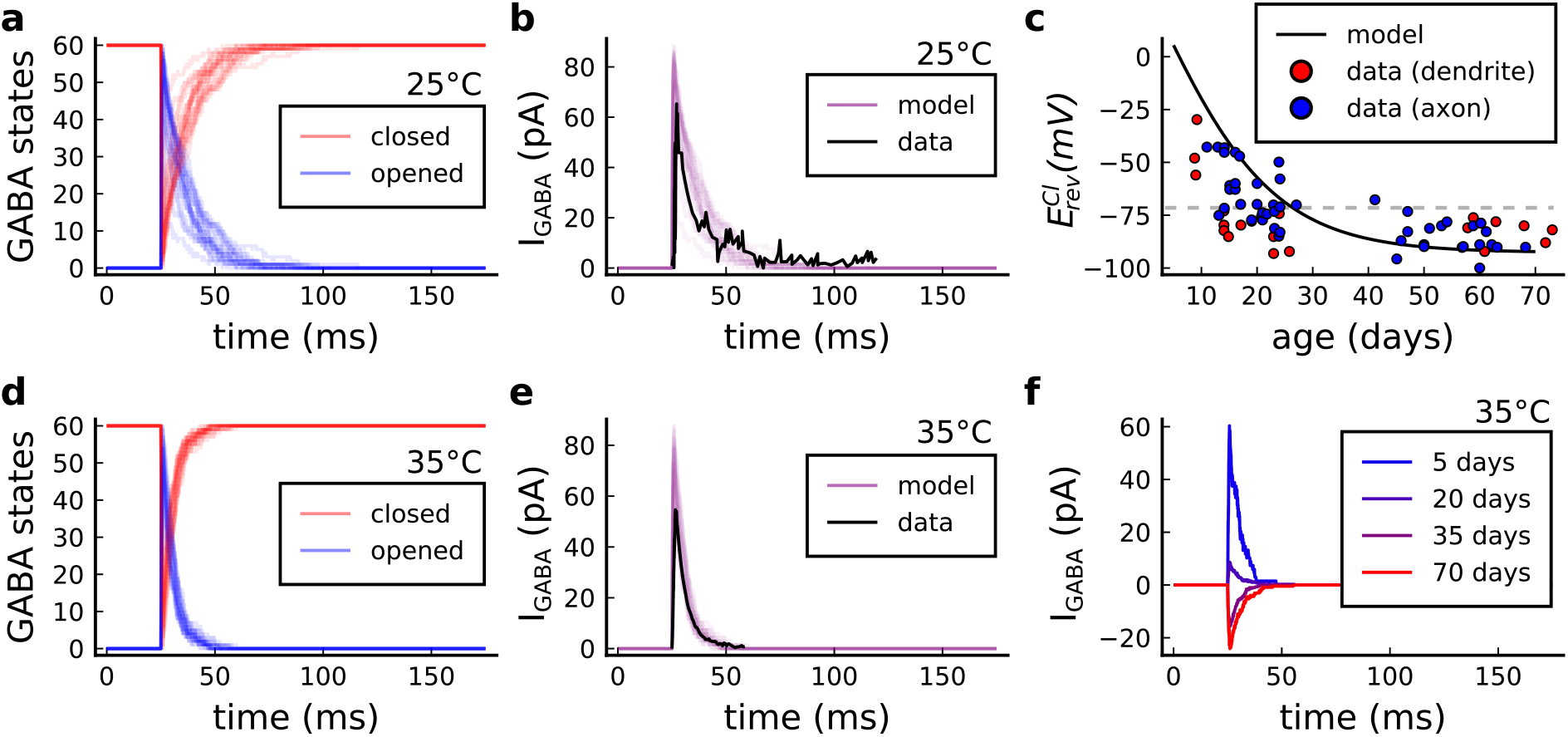
GABA(A)r current, kinetics and chloride reversal potential. **a**, States of GABA(A)r Markov chain at 25°*C* in response to a presynaptic stimulation. Opened = *O*_1_ + *O*_2_, closed = *C*_0_ + *C*_1_ + *C*_2_. **b**, Model and data comparison (***Otis and Mody, 1992***) for GABA(A)r current at 25°*C*. Even though data were recorded from P70 at 25°*C* and P15 at 35°*C,* we normalize the amplitude to invert the polarity and compare the decay time. This is done since the noise around P15 can either make GABAr excitatory or inhibitory as shown by *E_cl_* data in panel **c**. **c**, Chloride reversal potential 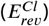 fitted to ***Rinetti-Vargas et al. (2017)*** data. Note that we used both profiles from axon and dendrite age-depended 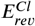 changes since exclusive dendrite data is scarce. **d**, States of simulated GABA(A)r Markov chain at 35°*C* in response to a presynaptic stimulation. **e**, Model and data comparison (***Otis and Mody, 1992***) for GABA(A)r current at 25°*C* (same normalization as in panel **b**). **f**, Change in the polarization of GABA(A)r currents given the age driven by the 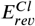.

#### GABA(A)r current and age switch

The GABA(A)r-driven current changes during development (***Meredith et al., 2003***) passing from depolarizing (excitatory) to hyperpolarizing (inhibitory) (***Chamma et al., 2012***). The reversal potential of chloride ions permeating GABA(A)r shifts from above the membrane resting potential (inward driving force – excitatory) to below the membrane resting potential (outward driving force – inhibitory) (***Rinetti-Vargas et al., 2017***). Such effect mediated by chloride ions is associated with the KCC2 pump (K Cl co-transporter) which becomes efficient in extruding chloride ions during maturation (***Rinetti-Vargas et al., 2017***). To cover the GABA(A)r age-dependent shift, we fit the chloride reversal potential 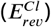 using the data published by ***Rinetti-Vargas et al. (2017)*** (***Figure 15c***):

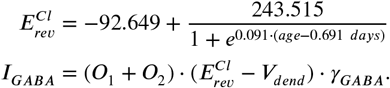

***Table 9*** presents the parameters to model the GABAr.

**Table 9.**
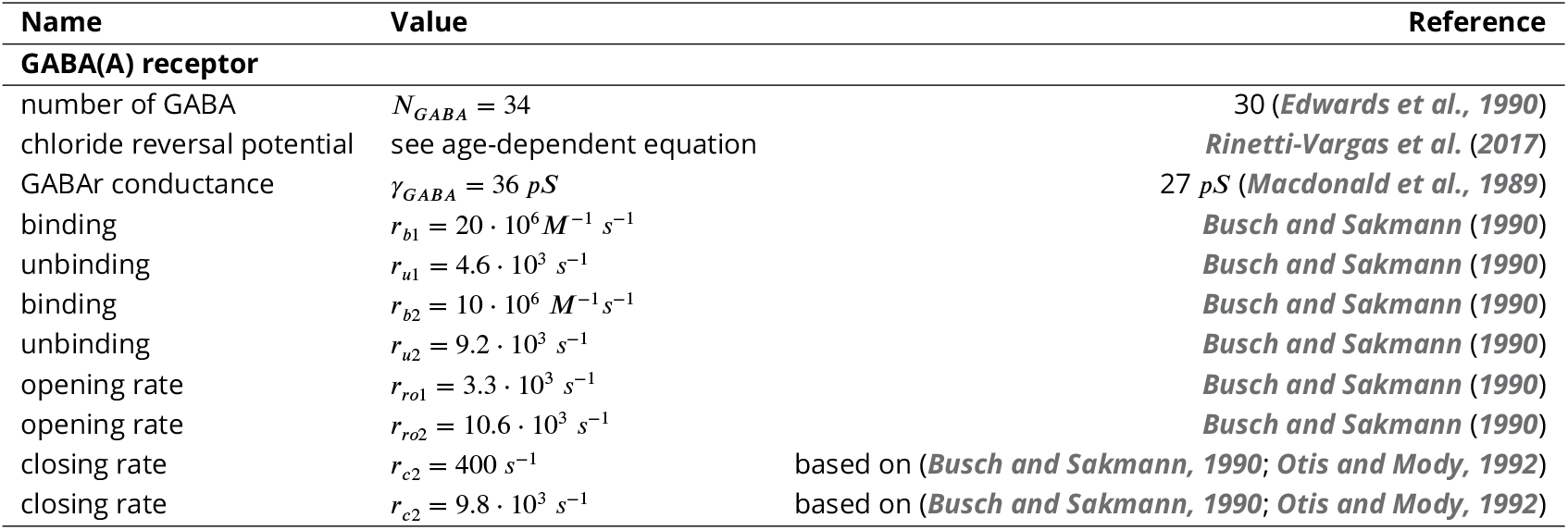
GABAr parameters.

### VGCC – T, R and L type

#### Markov chain

A stochastic VGCC model was devised using the channel gating measurements from rat CA1 (2-8 weeks) pyramidal neurons by ***Magee and Johnston (1995)*** at room temperature. Our model has three different VGCC subtypes described by the Markov chains in ***Figure 16***: the T-type (low-voltage), the R-type (medium-to-high-voltage) and the L-type (high-voltage).

**Figure 16.**
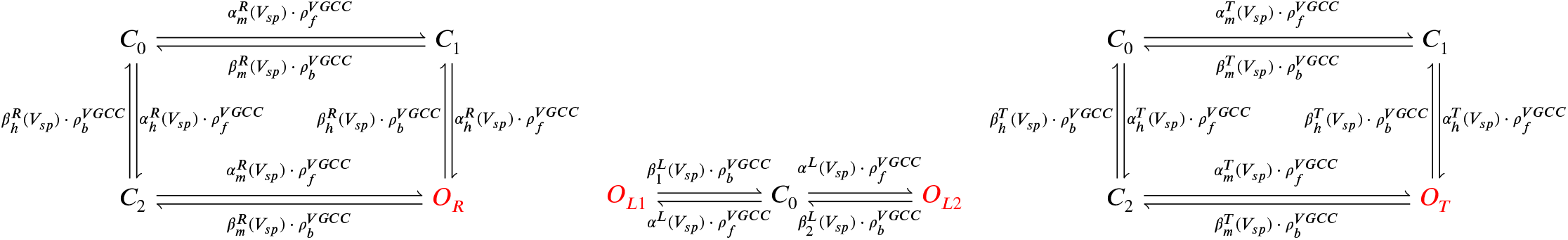
From left to right, R-, L-, and T-type VGCCs Markov chain adapted from Magee and Johnston(***Magee and Johnston, 1995***). The R-(left scheme) and T-type (right scheme) have a single open state (red colour), respectively, *O_r_* and *O_T_*. The L-type VGCC (middle) has two open states, *O*_*L*1_ and *O*_*L*2_.

The VGCC Markov chain derived from Magee and Johnston 1995 (***Magee and Johnston, 1995***) is composed of two gates (h,m) forT-(***Figure 17a,d**)* and R-types (***Figure 17b,e***) and a single gate for L-type (***Figure 17c***), as described in the equations below.

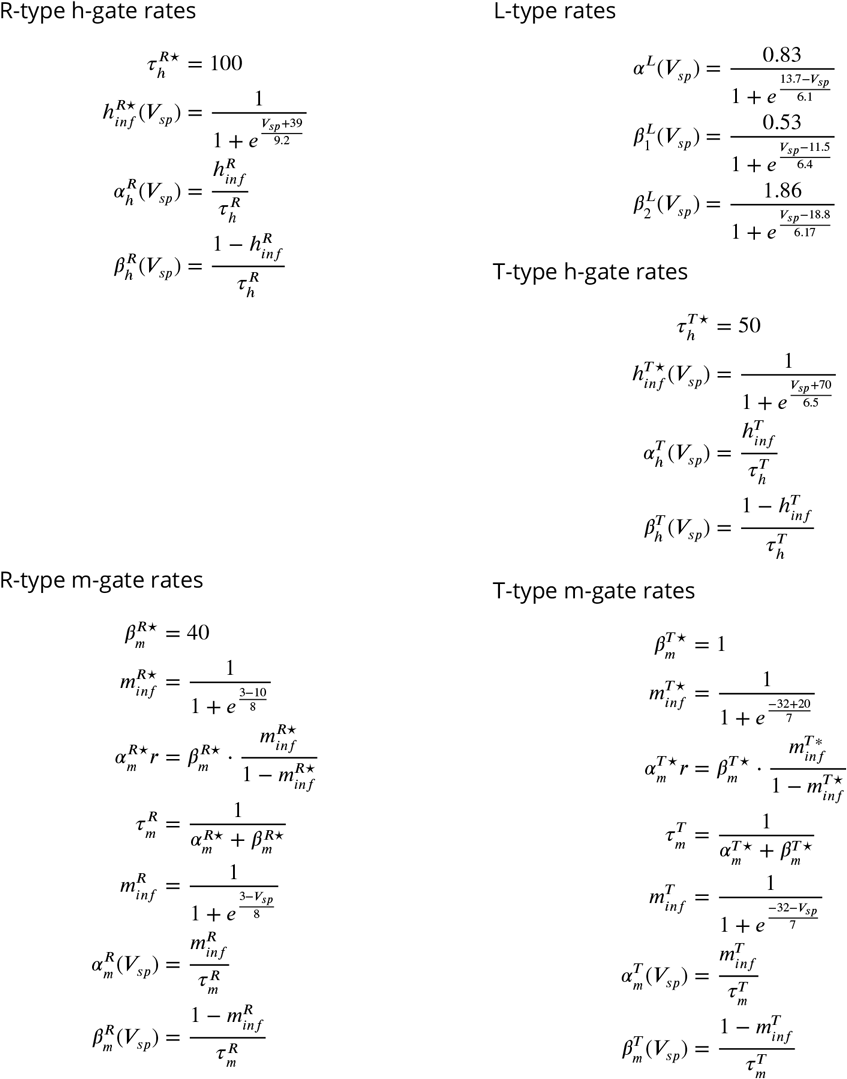

**Figure 17.**
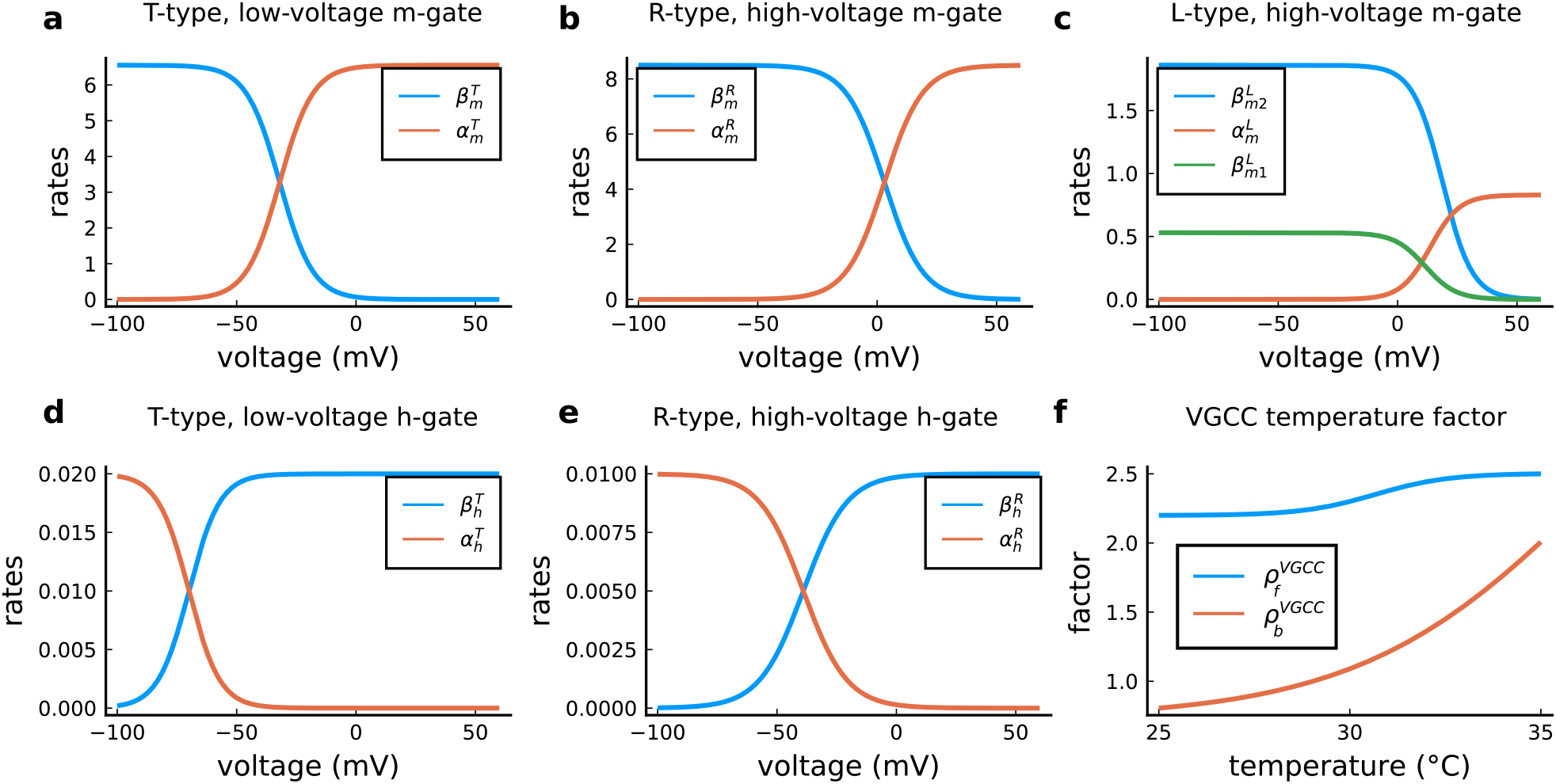
VGCC rates and temperature factors. **a**, Activation (*α_m_*(*V_sp_*)) and deactivation rates (*β_m_*(*y_sp_*)) for the T-type m-gate. **b**, Activation ( *α_m_*(*V_sp_*)) and deactivation rates (*β_m_)* for the R-type m-gate. **c**, Activation (*a_m_*(*y_sp_*)) and both deactivation rates 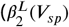 and 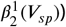 for the L-type VGCC. **d**, Activation (*a_h_*(*V_sp_*)) and deactivation rates (*β_h_*(*V_sp_*)) for the T-type h-gate. **e**, Activation (*a_h_*(*V_sp_*)) and deactivation rates (*β_h_*(*V_sp_*)) for the R-type h-gate. **f**, Temperature factor applied to all the rates, forward change 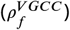 for the *a* rates and backward change 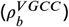 for the *β* rates.

#### VGCC and temperature

We used the same temperature factor for every VGCC subtype, respectively 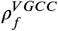 and 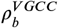 (see ***Figure 17f***), as follows:

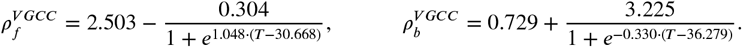

The VGCC subtypes are differently sensitive to temperature, with temperature factors for decay times ranging from 2 (***Iftinca et al., 2006***) to 50-fold (***Peloquin et al., 2008)***. It further complicates if T-type isoforms are considered. Indeed, they can have temperature factors that accelerate or slow down the kinetics. For instance, when passing from room to physiological temperatures, the isoform Ca_*v*_.3.1 has a closing time ~50 % faster (***Iftinca et al., 2006***) and the isoform Ca _*v*_3.1 becomes ~15 % slower. To simplify, the same temperature factor was adopted to all VGCC subtypes.

#### VGCC currents

The VGCC currents are integrated to the dendritic spine and estimated using the GHK ***Equation 14***, as follows:

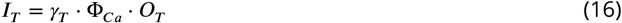

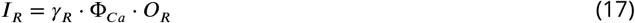

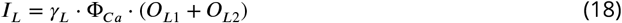

***Table 10*** presents the parameters to model the VGCC channels. VGCC rates and temperature factors are shown in ***Figure 17***.

**Table 10.** VGCC parameters

| Name | Value | Reference |
| --- | --- | --- |
| **VGCC** |  |  |
| VGCC T-type conductance | $\gamma_{CaT} = 12 \text{ pS}$ | same as (*Magee and Johnston, 1995*) |
| VGCC R-type conductance | $\gamma_{CaR} = 17 \text{ pS}$ | same as (*Magee and Johnston, 1995*) |
| VGCC L-type conductance | $\gamma_{CaL} = 27 \text{ pS}$ | same as (*Magee and Johnston, 1995*) |
| number of VGCCs | 3 for each subtype | 1 to 20 (*Higley and Sabatini, 2012*) |

#### SK channel

The small potassium (SK) channel produces hyperpolarizing currents which are enhanced in the presence of intracellular calcium elevations. We included SK channels to incorporate a key negative feedback loop between spine calcium and voltage due to the tight coupling that exists between SK channels to NMDAr function (***Adelman et al., 2012; Griffith et al., 2016***). Although SK channels can additionally be regulated by metabotropic glutamate receptors and muscarinic receptors (***Tigaret et al., 2016***), we did not include these regulatory steps in the model. The SK channel current was based on the description from ***Griffith et al. (2016)*** as follows:

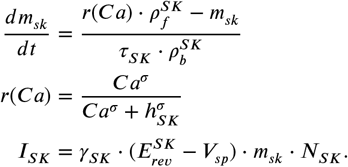

There is little information on how temperature effects SK channel function, but ***Van Herck et al. (2018)*** suggests a left-ward shift in the SK half-activation when changing from 37°*C* (*h_SK_* = 0.38 ± 0.02 *μM*) to 25°*C*(*h_SK_* = 0.23 ± 0.01 *μM*); that is a 65% decrease. Thus, to mimic temperature dependence of SK, we decided to decrease the decay time of the SK hyperpolarizing current by a factor of two when passing from physiological to room temperature.

***Table 11*** presents the parameters to model the SK channel.

**Table 11.** SK channel parameters.

| Name | Value | Reference |
| --- | --- | --- |
| **SK channel** |  |  |
| number of SK channels | $N_{SK} = 15$ | 10–200 (*Bock et al., 2019*) |
| SK conductance | $\gamma_{SK} = 10 \text{ pS}$ | *Maylie et al. (2004)* |
| SK reversal potential | $E_{rev}^{SK} = -90 \text{ mV}$ | *Griffith et al. (2016)* |
| SK half-activation | $h_{SK} = 0.333 \mu M$ | *Griffith et al. (2016)* |
| SK half-activation slope | $\sigma = 6$ | 4 (*Griffith et al., 2016*) |
| SK time constant | $\tau_{SK} = 6.3 \text{ ms}$ | *Griffith et al. (2016)* |

#### Enzymes – CaM, CaN and CaMKII

To model the enzymes dynamics, we adapted a monomeric CaM-CaMKII Markov chain from ***Chang et al. (2019)*** which was built on the model by ***Pepke et al. (2010)***. Our adaptation incorporates a simplified CaN reaction which only binds to fully saturated CaM. That is, CaM bound to four calcium ions on N and C terminals (see Markov chain in the ***Figure 18).*** A consequence of the Pepke coarsegrained model is that calcium binds and unbinds simultaneously from the CaM terminals (N,C). We assumed a lack of dephosphorylation reaction between CaMKII and CaN since ***Otmakhov et al. (2015)*** experimentally suggested that no known phosphatase affects CaMKII decay time which is probably caused only by CaM untrapping (***Otmakhov et al., 2015***). This was previously theorized in the Michalski’s model ***Michalski (2013)***, and it is reflected in Chang data (***Chang et al., 2019,2017***). The structure of the corresponding Markov chain is shown in ***Figure 18***.

**Figure 18.**
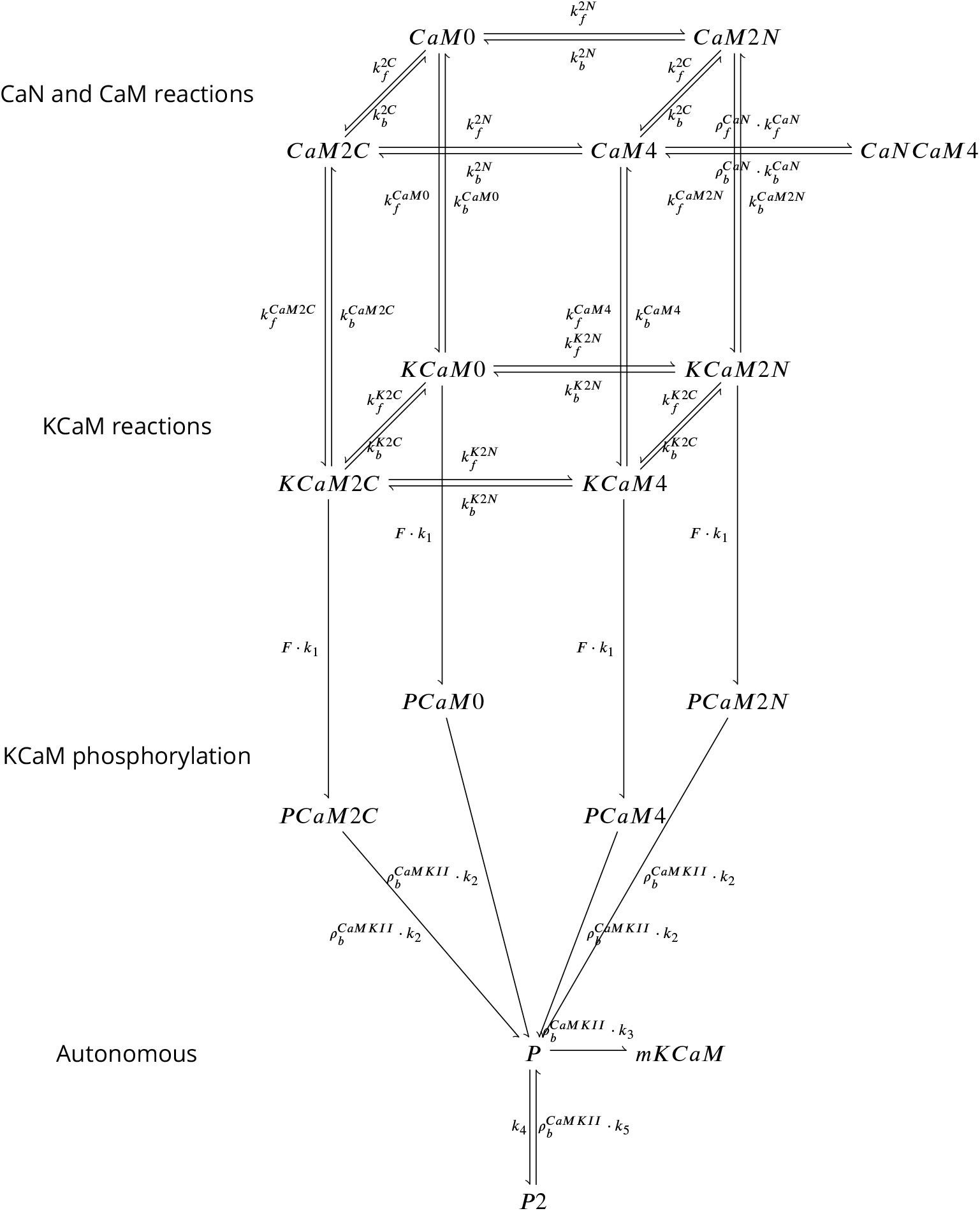
Coarse-grained model of CaM, CaMKII and CaN adapted from *Chang et al. (2019)* and *Pepke et al. (2010)*. Reaction from the CaM-Ca reactions (first layer) are attributed to 2Ca release and binding from different CaM saturation states CaM2C (2Ca bound to terminal C), CaM2N (2Ca bound to terminal N), CaM0 (no calcium bound), CaM4(Ca bound to both C and N terminal). Note that CaN is allowed to bind only to fully saturated CaM. Activated CaN is represented by the state CaNCaM4. Reactions between the first (CaM-Ca reactions) and the second layer (KCaM-Ca reactions) represent the binding of free/monomeric CaMKII (mKCaM) (***Pepke et al., 2010***) to different saturation levels of CaM. Reactions within the layer KCaM-Ca represent the binding of calcium to Calmodulin bound to CaMKII (KCaM0, KCaM2C, KCaM2N, KCaM4). Transition of layer KCaM-Ca reactions to layer KCaM-phosphorylation represents CaMKII bound to CaM that became phosphorylated (PCaM states) (***Pepke et al., 2010; Chang et al., 2017,2019***). PCaM can become self-phosphorylated (Autonomous layer with P and P^2^) and release CaM. Once the KCaM deactivates from autonomous states, it returns to free monomeric CaMKII (mKCaM). The CaMKII activity in this work represent the states (KCaM + PCaM + P + P^2^). See ***Chang et al. (2019)*** for further explanation on this system. CaNCaM4 represents the CaN activity.

***Chang et al. (2019)*** data provides a high-temporal resolution fluorescence measurements for CaMKII in dendritic spines of rat CA1 pyramidal neurons and advances the description of CaMKII self-phosphorylation (at room temperature). We modified Chang’s model of CaMKII unbinding rates *k*_2_,*k*_3_,*k*_4_,*k*_5_ to fit CaMKII dynamics at room/physiological temperature as shown by ***Chang et al. (2017)*** supplemental files. Previous modelling of CaMKII (***Chang et al., 2019; Pepke et al., 2010***) used a stereotyped waveform with no adaptation to model calcium. Our contribution to CaMKII modelling was to use calcium dynamics sensitive to the experimental conditions to reproduce CaMKII data, therefore, allowing us to capture physiological temperature measurements from ***Chang et al. (2017)***. Note that the CaMKII dynamic has two time scales and we capture only the fastest timescale which ends after stimulation ceases (at 60 s). The slowest dynamic occurs at the end of the stimulus, close to the maximum (***Figure 19a***). This can be caused by the transient volume increase in the spine as measured by ***Chang et al. (2017)***. ***Table 12*** shows the concentration of the enzymes and ***Table 13*** shows the parameters to model enzymes reactions in shown in ***Figure 18***.

**Figure 19.**
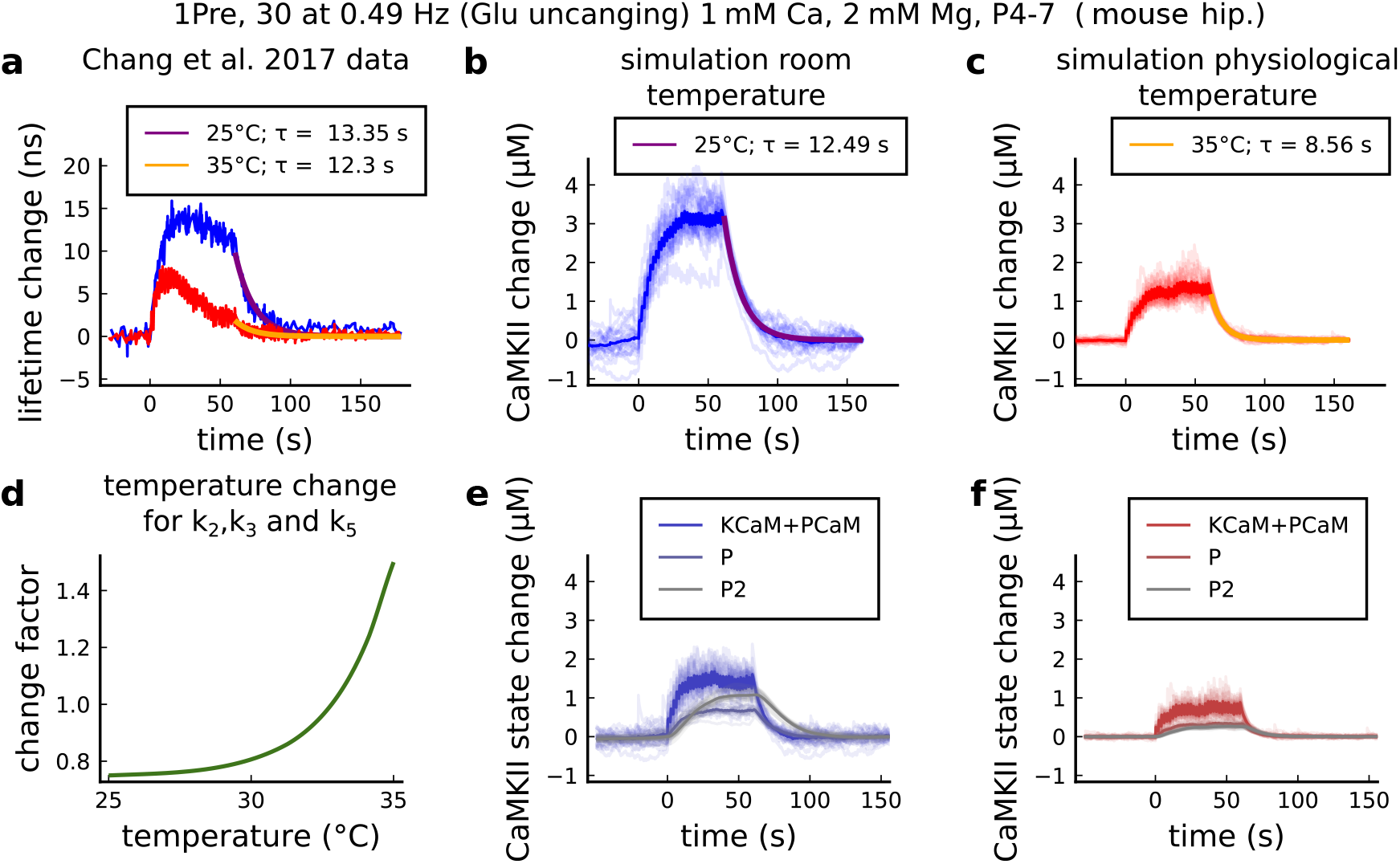
CaMKII temperature changes in the model caused by 1Pre, 30 at 0.49 Hz with glutamate uncaging (no failures allowed), 1mM Ca, 2mM Mg, P4-7 organotypic slices from mouse hippocampus. **a**, CaMKII fluorescent probe lifetime change measured by ***Chang et al. (2017)*** for 25°*C* (blue) and 35°*C* (red). The decay time (*τ*) was estimated by fitting the decay after the stimulation (30 pulses at 0.49Hz) using a single exponential decay, *y* = *a · e*^-1·*b*^; *τ* = 1/*b*. **b**, Simulation of the CaMKII concentration change (with respect to the baseline) at 25°*C* in response to same protocol applied in the panel **a**. The simulations on the panels **b, c, e, f** show the mean of 20 samples. **c**, Same as in panel **b** but for 35°*C*. **d**, Estimated temperature change factor for the dissociation rates *k*_2_, *k*_3_ and *k*_5_ in the Markov chain in ***Figure 18***. **e**, Change in the concentration of the CaMKII states (25°*C*) which are summed to compose CaMKII change in the panel **b. f**, Same as in panel **e** for 35°*C* with reference to the panel **c**.

**Table 12.** Concentration of each enzyme.

| Name | Value | Reference |
| --- | --- | --- |
| **Enzyme concentrations** |  |  |
| free CaM concentration (spine) | $CaM_{con} = 30 \mu\text{M}$ | *Kakiuchi et al. (1982)* |
| free KCaM concentration (spine) | $mKCaM_{con} = 70 \mu\text{M}$ | *Feng et al. (2011); Lee et al. (2009)* |
| free CaN spine concentration (spine) | $mCaN_{con} = 20 \mu\text{M}$ | >10 $\mu\text{M}$ (estimation from *Kuno et al. (1992)*) |

**Table 13.** Parameters for the coarse-grained model published in ***Pepke et al. (2010)*** and adapted by ***Chang et al. (2019)*** and this work. ***Pepke et al. (2010)*** rate adaptation for the coarse-grained model 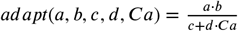. Refer to Figure 18f or definition of variables.

| REACTIONS | Value | Reference |
| --- | --- | --- |
| **Coarse-grained model, CaM-Ca reactions** |  |  |
| CaM0 + 2Ca ⇒ CaM2C | $k_f^{2C} = adapt(k_{on}^{1C}, k_{on}^{2C}, k_{off}^{1C}, k_{off}^{2C}, Ca)$ | *Pepke et al. (2010)* |
| CaM2N + 2Ca ⇒ CaM4 | | |
| CaM0 + 2Ca ⇒ CaM2N | $k_f^{2N} = adapt(k_{on}^{1N}, k_{on}^{2N}, k_{off}^{1N}, k_{off}^{2N}, Ca)$ | *Pepke et al. (2010)* |
| CaM2C + 2Ca ⇒ CaM4 | | |
| CaM2C ⇒ CaM0 + 2Ca | $k_b^{2C} = adapt(k_{off}^{1C}, k_{off}^{2C}, k_{on}^{1C}, k_{on}^{2C}, Ca)$ | *Pepke et al. (2010)* |
| CaM4 ⇒ CaM2N + 2Ca | | |
| CaM2N ⇒ CaM0 + 2Ca | $k_b^{2N} = adapt(k_{off}^{1N}, k_{off}^{2N}, k_{on}^{1N}, k_{on}^{2N}, Ca)$ | *Pepke et al. (2010)* |
| CaM4 ⇒ CaM2C + 2Ca | | |
| | $k_{on}^{1C} = 5 \cdot 10^6 M^{-1} s^{-1}$ | $1.2 \text{ to } 9.6 \cdot 10^6 M^{-1} s^{-1}$ (*Pepke et al., 2010*) |
| | $k_{on}^{2C} = 10 \cdot 10^6 M^{-1} s^{-1}$ | $5 \text{ to } 35 \cdot 10^6 M^{-1} s^{-1}$ (*Pepke et al., 2010*) |
| | $k_{on}^{1N} = 100 \cdot 10^6 M^{-1} s^{-1}$ | $25 \text{ to } 260 \cdot 10^6 M^{-1} s^{-1}$ (*Pepke et al., 2010*) |
| | $k_{on}^{2N} = 200 \cdot 10^6 M^{-1} s^{-1}$ | $50 \text{ to } 300 \cdot 10^6 M^{-1} s^{-1}$ (*Pepke et al., 2010*) |
| | $k_{off}^{1C} = 50 s^{-1}$ | $10 \text{ to } 70 s^{-1}$ (*Pepke et al., 2010*) |
| | $k_{off}^{2C} = 10 s^{-1}$ | $8.5 \text{ to } 10 s^{-1}$ (*Pepke et al., 2010*) |
| | $k_{off}^{1N} = 2000 s^{-1}$ | $1 \cdot 10^3 \text{ to } 4 \cdot 10^3 s^{-1}$ (*Pepke et al., 2010*) |
| | $k_{off}^{2N} = 500 s^{-1}$ | $0.5 \cdot 10^3 \text{ to } > 1 \cdot 10^3 s^{-1}$ (*Pepke et al., 2010*) |
| **Coarse-grained model, KCaM-Ca reactions** |  |  |
| KCaM0 + 2Ca ⇒ KCaM2C | $k_f^{K2C} = adapt(k_{on}^{K1C}, k_{on}^{K2C}, k_{off}^{K1C}, k_{off}^{K2C}, Ca)$ | *Pepke et al. (2010)* |
| KCaM2N + 2Ca ⇒ KCaM4 | | |
| KCaM0 + 2Ca ⇒ KCaM2N | $k_f^{K2N} = adapt(k_{on}^{K1N}, k_{on}^{K2N}, k_{off}^{K1N}, k_{off}^{K2N}, Ca)$ | *Pepke et al. (2010)* |
| KCaM2C + 2Ca ⇒ KCaM4 | | |
| KCaM2C ⇒ KCaM0 + 2Ca | $k_b^{K2C} = adapt(k_{off}^{K1C}, k_{off}^{K2C}, k_{on}^{K1C}, k_{on}^{K2C}, Ca)$ | *Pepke et al. (2010)* |
| KCaM4 ⇒ KCaM2N + 2Ca | | |
| KCaM2N ⇒ KCaM0 + 2Ca | $k_b^{K2N} = adapt(k_{off}^{K1N}, k_{off}^{K2N}, k_{on}^{K1N}, k_{on}^{K2N}, Ca)$ | *Pepke et al. (2010)* |
| KCaM4 ⇒ KCaM2C + 2Ca | | |
| | $k_{on}^{K1C} = 44 \cdot 10^6 M^{-1} s^{-1}$ | *Pepke et al. (2010)* |
| | $k_{on}^{K2C} = 44 \cdot 10^6 M^{-1} s^{-1}$ | *Pepke et al. (2010)* |
| | $k_{on}^{K1N} = 76 \cdot 10^6 M^{-1} s^{-1}$ | *Pepke et al. (2010)* |
| | $k_{on}^{K2N} = 76 \cdot 10^6 M^{-1} s^{-1}$ | *Pepke et al. (2010)* |
| | $k_{off}^{K1C} = 33 s^{-1}$ | *Pepke et al. (2010)* |
| | $k_{off}^{K2C} = 0.8 s^{-1}$ | $0.49 \text{ to } 4.9 s^{-1}$ (*Pepke et al., 2010*) |
| | $k_{off}^{K1N} = 300 s^{-1}$ | *Pepke et al. (2010)* |
| | $k_{off}^{K2N} = 20 s^{-1}$ | $6 \text{ to } 60 s^{-1}$ (*Pepke et al., 2010*) |
| **Coarse-grained model, CaM-mKCaM reactions** |  |  |
| CaM0 + mKCaM ⇒ mKCaM0 | $k_f^{CaM0} = 3.8 \cdot 10^3 M^{-1} s^{-1}$ | *Pepke et al. (2010)* |
| CaM2C + mKCaM ⇒ mKCaM2C | $k_f^{CaM2C} = 0.92 \cdot 10^6 M^{-1} s^{-1}$ | *Pepke et al. (2010)* |
| CaM2N + mKCaM ⇒ mKCaM2N | $k_f^{CaM2N} = 0.12 \cdot 10^6 M^{-1} s^{-1}$ | *Pepke et al. (2010)* |
| CaM4 + mKCaM ⇒ mKCaM4 | $k_f^{CaM4} = 30 \cdot 10^6 M^{-1} s^{-1}$ | $14 \text{ to } 60 \cdot 10^6 M^{-1} s^{-1}$ (*Pepke et al., 2010*) |
| mKCaM0 ⇒ CaM0 + mKCaM | $k_b^{CaM0} = 5.5 s^{-1}$ | *Pepke et al. (2010)* |
| mKCaM2C ⇒ CaM2C + mKCaM | $k_b^{CaM2C} = 6.8 s^{-1}$ | *Pepke et al. (2010)* |
| mKCaM2N ⇒ CaM2N + mKCaM | $k_b^{CaM2N} = 1.7 s^{-1}$ | *Pepke et al. (2010)* |
| mKCaM4 ⇒ CaM0 + mKCaM | $k_b^{CaM4} = 1.5 s^{-1}$ | $1.1 \text{ to } 2.3 s^{-1}$ (*Pepke et al., 2010*) |
| **Coarse-grained model, self-phosphorylation reactions** |  |  |
| KCaM0 ⇒ PCaM0 | | |
| KCaM2N ⇒ PCaM2N | $k_1 = 12.6 s^{-1}$ | *Chang et al. (2019)* |
| KCaM2C ⇒ PCaM2C | | |
| KCaM4 ⇒ PCaM4 | | |
| Fraction of activated CaMKII | $F = CaMKII / mKCaM_{con}$ | see *Equation 19* (*Chang et al., 2019*) |
| PCaM0 ⇒ P+CaM0 | | |
| PCaM2N ⇒ P+CaM2N | $k_2 = 0.33 s^{-1}$ | $0.33 s^{-1}$ ; adapted from (*Chang et al., 2019*) |
| PCaM2C ⇒ P+CaM2C | | |
| PCaM4 ⇒ P+CaM4 | | |
| P ⇒ mKCaM | $k_3 = 4 \cdot 0.17 s^{-1}$ | $0.17 s^{-1}$ adapted from (*Chang et al., 2019*) |
| P ⇒ P2 | $k_4 = 4 \cdot 0.041 s^{-1}$ | $0.041 s^{-1}$ adapted from (*Chang et al., 2019*) |
| P2 ⇒ P | $k_5 = 8 \cdot 0.017 s^{-1}$ | $0.017 s^{-1}$ adapted from (*Chang et al., 2019*) |
| **Calcineurin model, CaM-CaM4 reactions** |  |  |
| CaM4+mCaN ⇒ mCaN+CaM4 | $k_f^{CaN} = 10.75 \cdot 10^6 M^{-1} s^{-1}$ | $46 \cdot 10^6 M^{-1} s^{-1}$ (*Quintana et al., 2005*) |
| mCaN+CaM4 ⇒ CaM4+mCaN | $k_b^{CaN} = 0.02 s^{-1}$ | fit from Fujii et al. 2014 (*Fujii et al., 2013*)<br>see *Figure 20* |

The CaN concentration was chosen as the total concentration used in a previous model (***Stefan et al., 2008***) (1.6 μM) scaled by a factor of 12 due to a higher CaN concentration in dendritic spines (***Goto et al., 1986; Baumgärtel and Mansuy, 2012***) and taking into account the discrepancy between different CaN concentration studies (***Kuno et al., 1992; Goto et al., 1986): Kuno et al. (1992)*** proposes 9.6 μg/mg(7.0 + 2.6 μg/mg for Aα and Aβ isoforms) for the catalytic subunit A of CaN (CaNA) in the hippocampus, while ***Goto et al. (1986)*** proposes 1.45 μg/mg (presumably for both isoforms). There is therefore a lack of consensus on CaN concentration in neurons, which seems to range between 1 and 10 μg/mg. However, models of CaN in spines (Stefan et al., 2008) use low values of CaN concentration (eg. 1.6 μM) not specific to dendritic spines without considering that these values are taken from the whole neuropil. There is little information on CaN concentration in spines, but ***Kuno et al. (1992)*** note that the concentration of CaN is 50% to 84% higher in synaptosomes than in neuronal nuclei. With this information in mind, we set CaN spine concentration 20 μM in our model. CaN was entirely activated through CaM for the following reason: CaNA is activated by calcium-CaM in a highly cooperative manner (Hill coefficient 2.8-3), whereas the activation of CaN by calcium (via CaNB) is at most 10% of that achieved with CaM (***Stemmer and Klee, 1994)***. In other words, CaNA affinity for CaM is 16 nM to 26 pM (***Creamer, 2020***), while CaNB affinity for calcium ranges from 15 μM to 24 nM (***Kakalis et al., 1995***). CaN decay time was modeled using experimental spine CaN activity dynamics measured in ***Fujii et al. (2013)***.

#### The lack of reactions between CaN and CaMKII

The protein phosphatases responsible for CaMKII dephosphorylation have not been established unequivocally (***Lisman, 1989***). Our model of CaMKII is based directly on a quantitative model fit to FRET imaging data (***Chang et al., 2017, 2019***), which implicitly account for the effects of any ‘hidden’ phosphatases, absorbing their contribution into the decay rates of the CaMKII activity. As pointed out by ***Otmakhov et al. (2015)***, FRET sensor imaging of CaMKII activity unfortunately does not capture the identity of the phosphatases involved in the dephosphorylation of CaMKII. More specifically, ***Otmakhov et al. (2015)*** observed no significant changes in the decay constant of their CaMKII FRET sensor when selectively inhibiting PP1 and PP2A. Given that these two phosphatases are widely used in models to determine plasticity, we believe that our model is more aligned with data of CaMKII activity *in vivo.*

Yet, our decision to include CaN in the model was determined by the evidence supporting CaN as the strongest candidate for calcium-sensitive protein phosphatase in the brain (***Baumgärtel and Mansuy, 2012***). Furthermore, the central role of CaN in synaptic plasticity has been demonstrated both pharmacologically and with genetic manipulation (***Onuma et al., 1998; Malleret et al., 2001***).

#### Temperature effects on enzymatic activity

We included temperature factors in the coarse-grained model using Changs data (***Chang et al., 2019***), as shown in ***Figure 19***. For CaMKII, we fit the modified dissociation rates of the phosphorylation states *k*_2_, *k*_3_ and *k*_5_ to match the data on relative amplitude and decay time using the following logistic function:

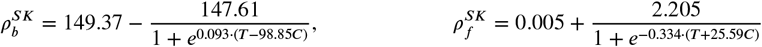

For CaN, we fit the ***Fujii et al. (2013)*** data at 25°*C* as seen in ***Figure 20a***. However, since CaNCaM dissociation rates at physiological temperatures were not reported, we set the temperature factorto CaN that fits the outcomes of the protocols we proposed to reproduce. A reference value from the CaN-AKAP79 complex (***Li et al., 2012***) showed a *Q*_10_ = 4.46 = (2.19 *s*^-1^/9.78 *s*^-1^) which is nearly the temperature factor used in our model for CaN. Therefore, both the association and dissociation rates are modified using the following logistic functions:

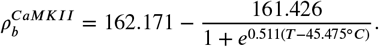

**Figure 20.**
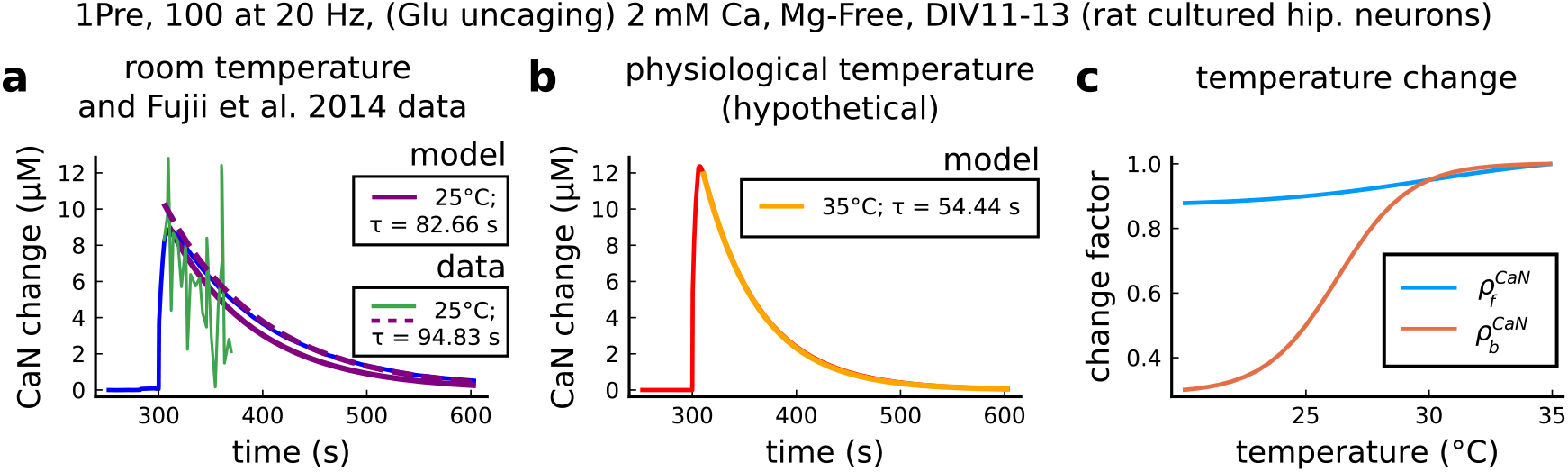
CaN temperature changes in our model caused by 1Pre, 100 at 20 Hz with glutamate uncaging (no failures allowed), 2mM Ca, Mg-free, 11-13 days in vitro. **a**, Simulated CaN change (blue solid line) in response to the same stimuli of the CaN measurement from ***Fujii et al. (2013)*** RY-CaN fluorescent probe (green solid line). The decay time (*τ*) estimated from data (*y* = *a · e*^-1·*b*^) is 94.83 s (dashed purple line) and 82.66 s for our model (solid purple line). **b**, Simulated CaN change for physiological temperature with decay time of 54.44 s. **c**, Temperature change, 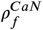 and 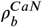, applied to CaN association and dissociation rates.

#### Geometrical Readout

We describe here the geometrical readout mechanism which allows for plasticity outcome assignment. First, we define the following variables which are representative of “active CaMKII” and “active CaN”:

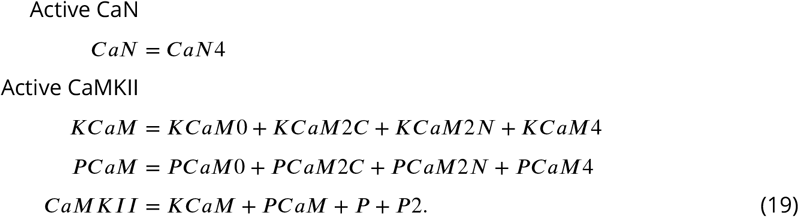

Calcium entry in the spine initiates a cascade of events that ultimately leads to long term plasticity changes. Specific concentrations of CaMKII and CaN trigger activation functions *act_D_* and *act_P_* when they belong to one of the two polygonal regions (P and D), termed plasticity regions in the main text:

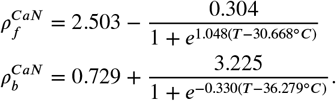

The variables *act_D_* and *act_p_* act as low pass filters of CaMKII and CaN activities with some memory of previous passages in the respective plasticity regions. To specify the LTP/LTD rates, termed *D_rale_* and *P_rate_*, we use the activation functions, *act_D_* and *act_p_*, as follows:

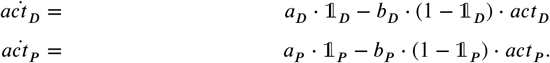

The Markov plasticity chain (see ***Figure 21***) starts with initial conditions *NC* = 100, *LTD* = 0 and *LTP* = 0. ***Figure 22*** shows how the readout works to predict plasticity for a single orbit. ***Figure 22a*** shows the enzyme’s activity alone which is combined to form an orbit as shown in ***Figure22b***. The region indicator of the respective orbit is shown in ***Figure 22c***. Simultaneously, ***Figure 22d*** depicts the leaky activation *act_P_* and *act_D_*, which will define the rate of plasticity induction in ***Figure 22e and f***. The rates in the plasticity Markov chain will not reset to 0 if the orbit leaves the readout. The plasticity Markov chain is shown in ***Figure 22g*** with the prediction outcome represented as a weight change (%). ***Figure 22h*** shows the rate, *P_rate_* and *D_rate_*, activation profile. The LTP activation rate is steep, meaning that orbits do not need to spend a long time inside the readout to promote LTP induction, while the LTD region requires five-fold longer activation times. ***Table 14*** shows the parameters that define the polygons of the plasticity regions (see ***Figure 22b***).

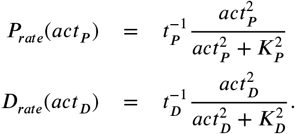

**Figure 21.**
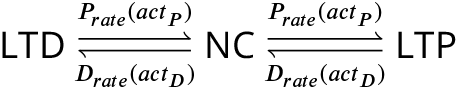
Plasticity Markov Chain.

**Figure 22.**
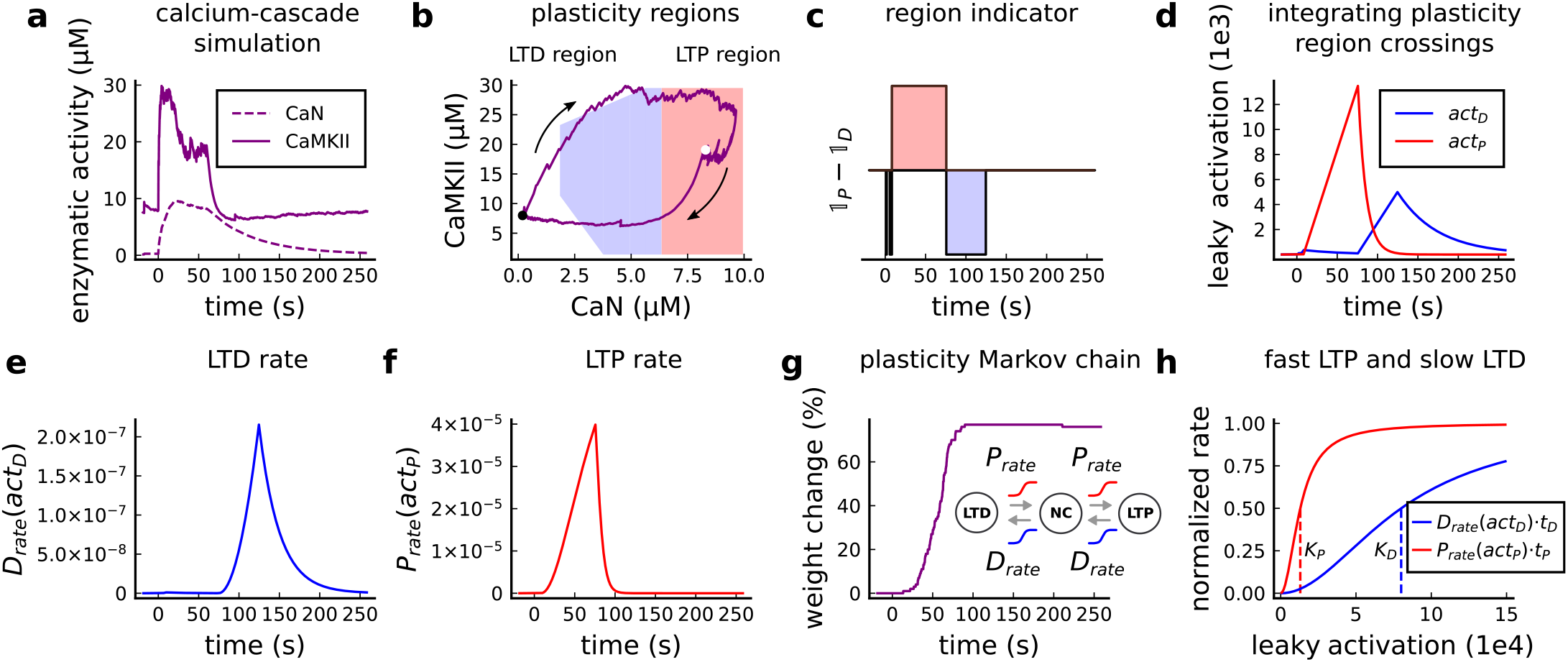
Plasticity readout for the protocol 1Pre2Post10, 300 at 5Hz, from *Tigaret et al. (2016)*. **a**, CaMKII and CaN activity in response to protocol 1Pre2Post10. **b**, Enzymatic joint activity in the 2D plane showing LTP and LTD’s plasticity regions. The black point marks the beginning of the stimulation, and the white point shows the end of the stimulation after 60 s. **c**, Region indicator illustrating how the joint activity crosses the LTP and the LTD regions. **d**, The leaky activation functions are used as input to the LTP and LTD, ratesrespectively. The activation function has a constant rise when the joint-activity is inside the region, and exponential decay when it is out. **e**, The LTD rate in response to the leaky activation function, *act_D_*, in panel **d**. Note that this rate profile occurs after the stimulation is finished (60 s). The joint-activity is returning to the resting concentration in panel A. **f**, The LTP rate in response to the leaky activation function, *act_P_*, in panel D. **g**, Outcome of the plasticity Markov chain in response to the LTD and LTP rates. The EPSP change (%) is estimated by the difference between the number of processes in the states LTP and LTD, *LTP – LTD.* **h**, Normalized LTP and LTD rates (multiplied to their respective time constant, *t_D_, t_P_*) sigmoids. The dashed line represents the half-activation curve for the LTP and LTD rates. Note in panel **d** that the leaky activation function reaches the half-activation *K_p_* = 1.3*e*4.

**Table 14.**
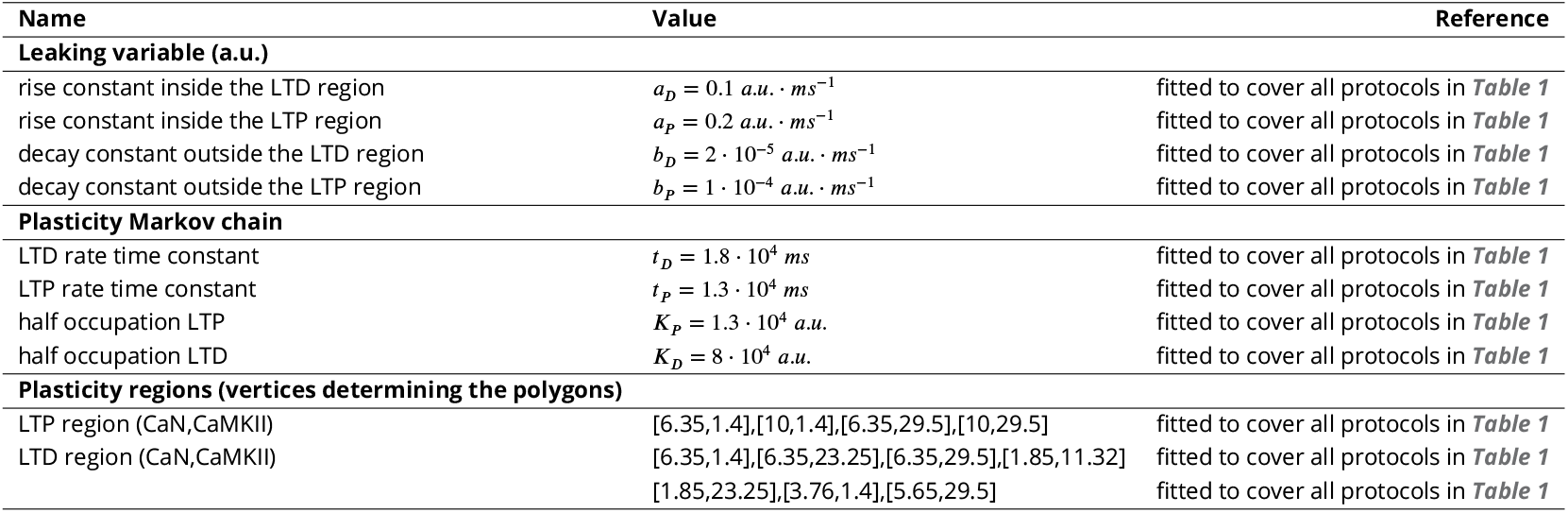
Parameters to define the plasticity readout.

#### Positioning of the boundaries of the plasticity regions

The tuning of the plasticity region boundaries was based on four different experiments. The LTP region was defined using Tigaret (***Figure 3***). The refinement of the LTD region was made using the simulated dynamics from ***Inglebert et al. (2020)*** (***Figure 6d***, top part of the LTD boundary) and Dudek and ***Dudek and Bear (1992, 1993)*** (***Figure 4d*** and ***Figure 5f***, bottom-left part of the LTD boundary).

## Supplemental files

***Figure 3-Figure Supplement 1*** shows best fit to the ***Tigaret et al. (2016)*** data from seven spike-timing dependent plasticity protocols, for three leading STDP models in the field: classic pairwise STDP (***Song et al., 2000***), triplet STDP (***Pfister and Gerstner, 2006***), and calcium-based Graupner-Brunel STDP (***Graupner and Brunel, 2012***) models. Parameters for each model that mimized the mean-squared error with the data were discovered using Bayesian optimization using the Bayesian Optimization package in the Julia programming language. ***Figure 4-Figure Supplement 1*** shows variations of ***Dudek and Bear (1992)*** parameters for [Ca^2+^]_o_, [Mg^2+^]_o_, temperature and dendritic spine distance from the soma. Also, it shows the Poisson spike train protocol (as in *Figure 7g,h*.) for temperature and age parameters obtained from an estimation of the body temperature regulation during development (or thermoregulation maturation, also called maturation of temperature homeostasis, estimated in ***Figure 3-Figure Supplement 1g***). ***Figure 5-Figure Supplement 1*** expands the presynaptic burst strategy hypothesized to recover the LTD in adult slices (***Figure 5c***) for 900 pairing repetitions. Also, ***Figure 5-Figure Supplement 1*** tries to isolate the contribution of each age-dependent mechanism (NMDAr, GABAr, BaP efficiency switches) for 3 and 5 Hz predictions in ***Dudek and Bear (1993)*** experiment. We fixed each of the three mechanisms coding for age in our model at P5 and P50, to observe how they shape the plasticity. Note the experiment in ***Figure 6-Figure Supplement 1d-i*** is only to theoretically show how each age mechanism contributes to plasticity in ***Figure 5***. Also we compare predictions between different STDP experiments across age. ***Figure 3-Figure Supplement 4*** presents modifications of ***Inglebert et al. (2020)*** STDP experiment and the reproduction of ***Mizuno et al. (2001)*** data. ***Figure 6-Figure Supplement 2*** shows multiple aspects related to temperature in STDP experiments and the temperature and age choices for the publications described in ***Table 1*** compared to physiological conditions. We estimate how the rat’s body temperature physiologically evolves in function of age using ***McCauley et al. (2020)*** and ***Wood et al. (2016)*** data.

**Figure 3 – Supplement 1.**
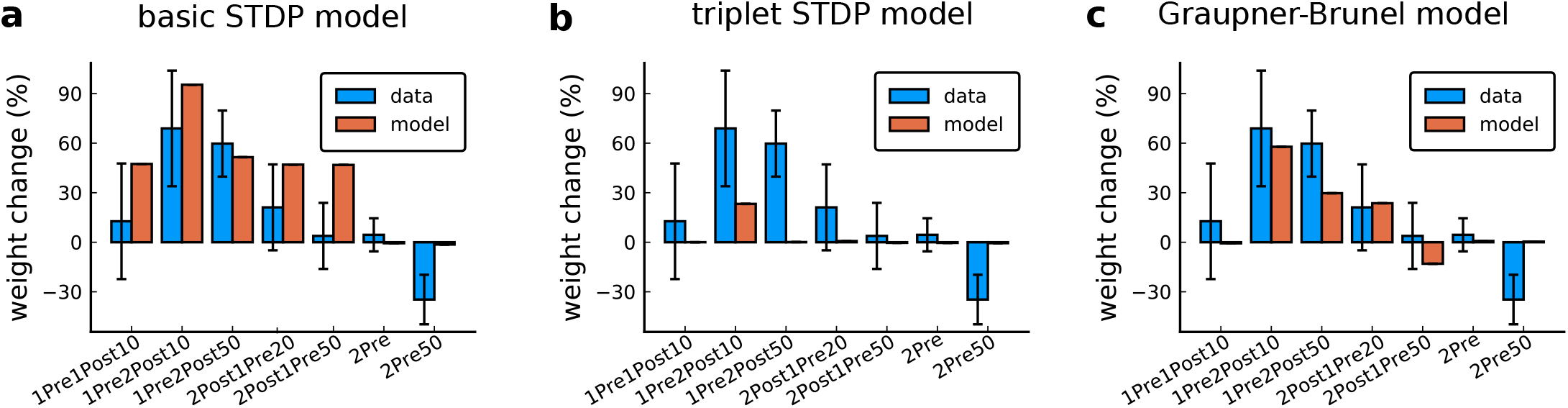
Standard models for predicting plasticity fail to account for the data from *Tigaret et al. (2016)*. **a–c**, Mean weight change for the Tigarets data (blue), error bars denote ±1 s.d. Plasticity protocols indicated by labels on x-axis. Green bars show mean plasticity predicted for the same protocols by classic STDP (***Song et al., 2000***) (panel a), triplet STDP (***Pfister and Gerstner, 2006***) (panel b), or Graupner-Brunel calcium-based STDP (***Graupner and Brunel, 2012***) model (panel c).

**Figure 3 – Supplement 2.**
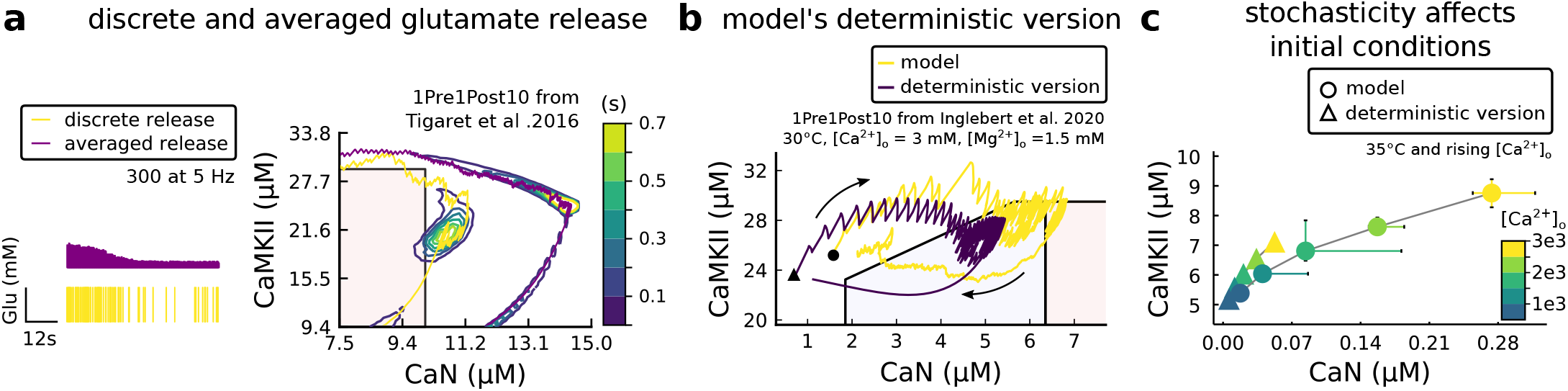
Comparison showing different roles of stochasticity in the model. **a**, Left, Glutamate concentration from a single realization of the model (yellow) and averaged Glutamate concentration (purple) from 100 repetitions of the model for 300 pulses train at 5 Hz. Right, 1Pre1Post10 from ***Tigaret et al. (2016)*** using the model (yellow) and a version of the model (purple) in which the glutamate concentration is the average one (as in Left panel). The time spent (s) is shown for the different glutamate release modes (stochastic and averaged) with an example trajectory (purple and solid yellow lines). There are no failures in averaged release; therefore, enzymes are over-activated. **b**, A comparison between our model and a fully deterministic version for the 1Pre1Post10 from ***Inglebert et al. 2020***(***Inglebert et al., 2020***). Note the significant mismatch, which does not allow the deterministic model to reach the LTP region that determines the plasticity outcome. This effect is mainly caused by the stochastic calcium sources, which the deterministic model fails to reproduce. The black triangle (circle) marks the initial conditions of the deterministic version (model). This initial condition is reached by letting the model evolve with no input. **c** The initial conditions are increasingly different when comparing the model and its deterministic version for rising concentrations of external calcium concentrations.

**Figure 3 – Supplement 3.**
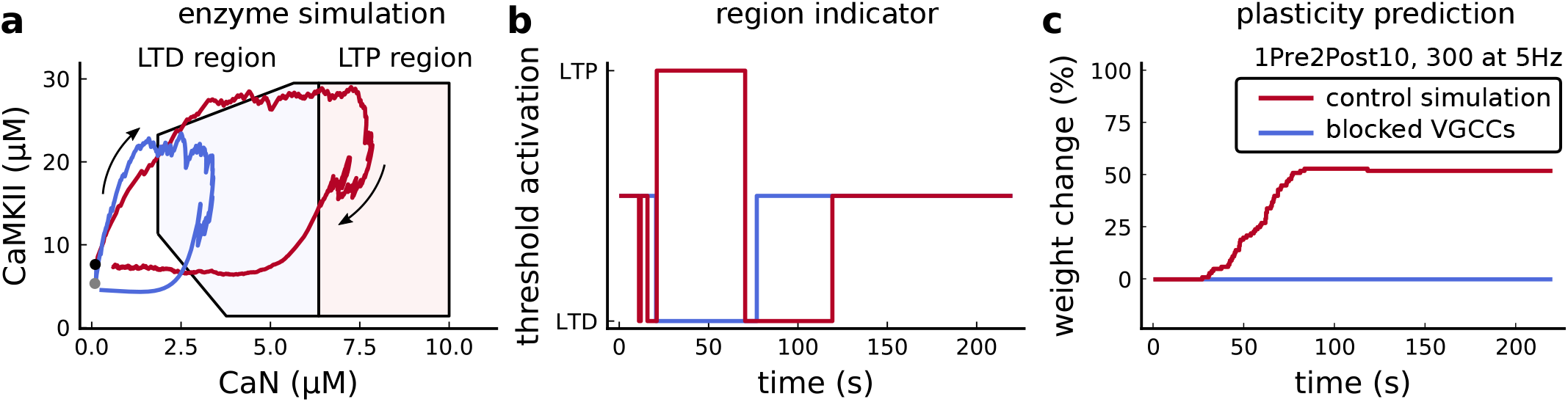
Effects of blocking VGCCs. **a**, Combined enzyme activity of the experiment 1Pre2Post10, 300 at 5 Hz described in ***Tigaret et al. (2016)*** with and without VGCCs (legend in panel **c**). The arrows indicate time flow, and the grey and black dots represent the initial conditions. Note the effect of VGCC blocking on the initial conditions. **b**, Region indicator associated to panel **a**. **c** Plasticity prediction for the simulated experiment with and without VGCCs. Note that when VGCCs are blocked LTP cannot be induced, in agreement with ***Tigaret et al. (2016)*** experimental data.

**Figure 3 – Supplement 4.**
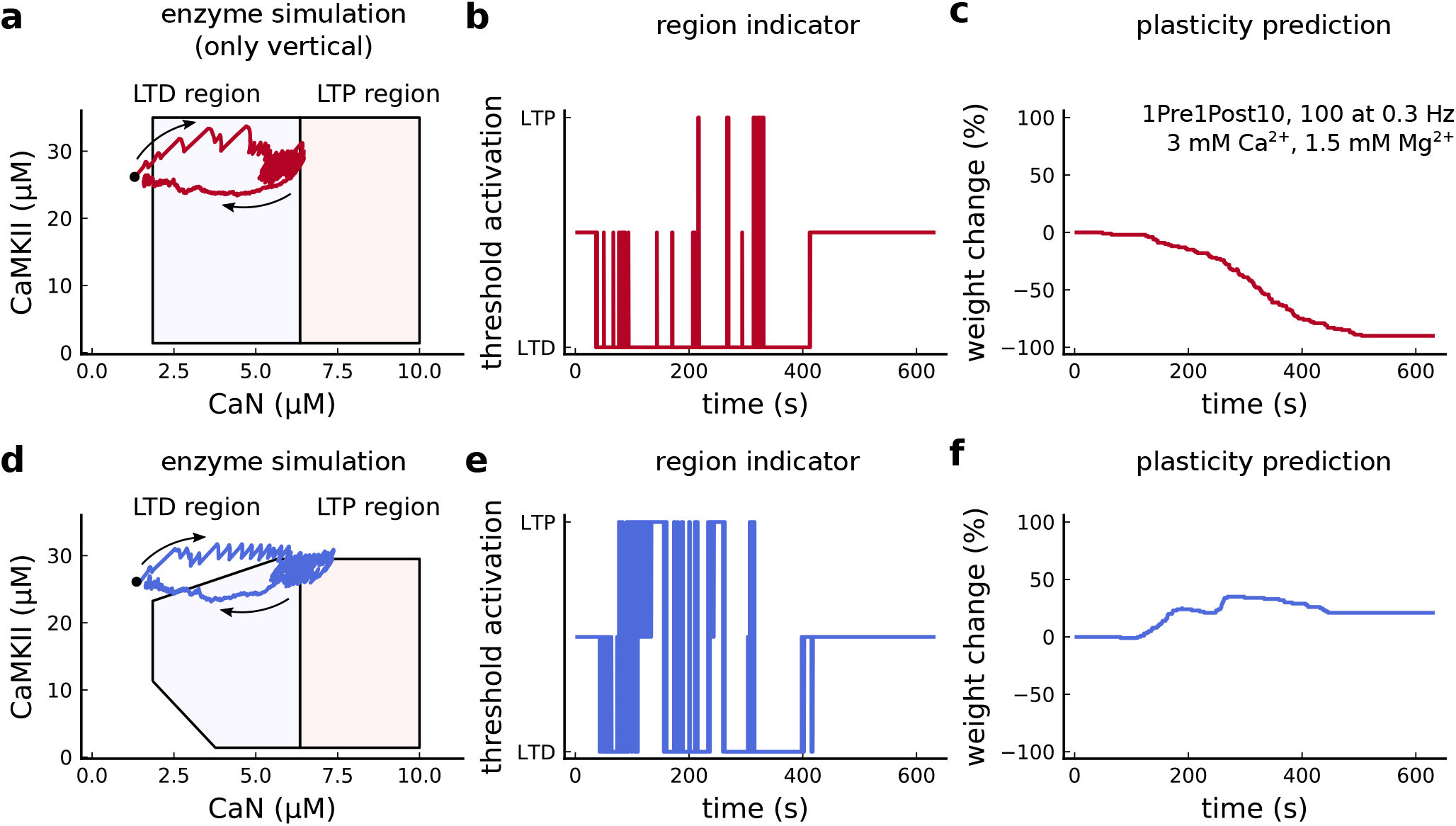
Exclusively setting vertical boundaries (no CaMKII selectivity) fails to capture the correct plasticity outcome. **a** Combined activity of the protocol 1Pre1Post10,100 at 0.3 Hz with experimental conditions as in Figure 6c considering the polygonal regions responding only to CaN thresholds. Note that most of the activity resides in the LTD region. The arrows indicate time flow and black dot represents the initial condition. **b,** Region indicator related to panel **a**. **c,** Plasticity prediction shows LTD, instead of LTP. **d,** Same as **a** but considering the plasticity regions sensitivity both to CaMKII and CaN. **e,** Region indicator related to panel **d**. **f,** Plasticity prediction for panel **d** showing LTP agreeing with data described in ***Figure 6c***.

**Figure 3 – Supplement 5.**
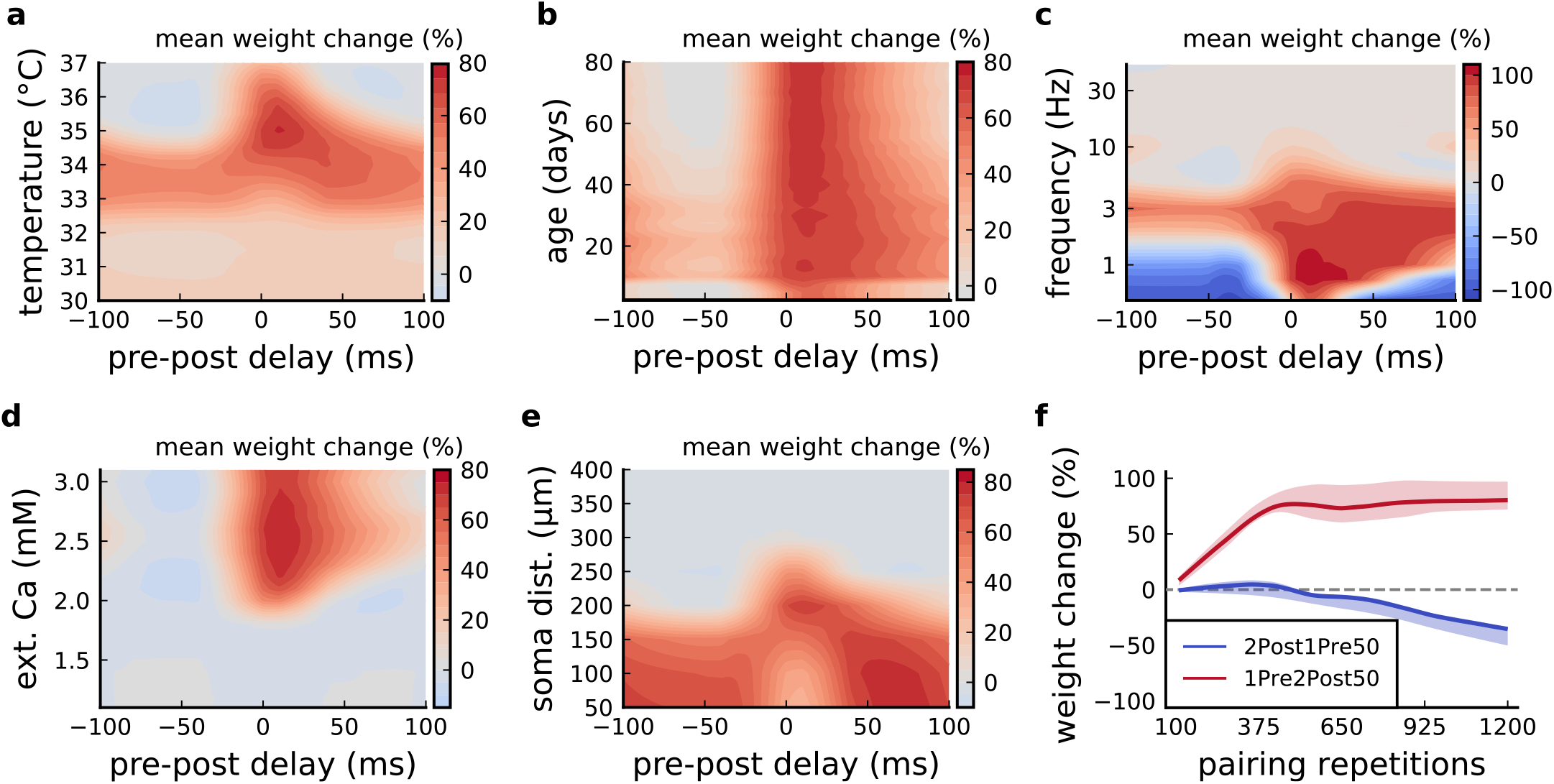
Varying ***Tigaret et al. (2016)*** experimental parameters. **a**, Mean synaptic weight change for 1Pre2Post(delay) varying the temperature. **b**, Mean synaptic weight change for 1Pre2Post(delay) varying the age. **c**, Mean synaptic weight change for 1Pre2Post(delay) varying the frequency. **d**, Mean synaptic weight change for 1Pre2Post(delay) varying the [Ca^2+^]_o_. **e**, Mean synaptic weight change for 1Pre2Post(delay) varying the distance from the soma. A similar trend in distal spines was previously found in ***Ebner et al. (2019)***. **f**, Mean synaptic weight change of 1Pre2Post50 and 2Post1Pre50 when number of pulses increases or decreases. Note the similarity with *Mizuno et al. (2001)* in ***Figure 161c***.

**Figure 4 – Supplement 1.**
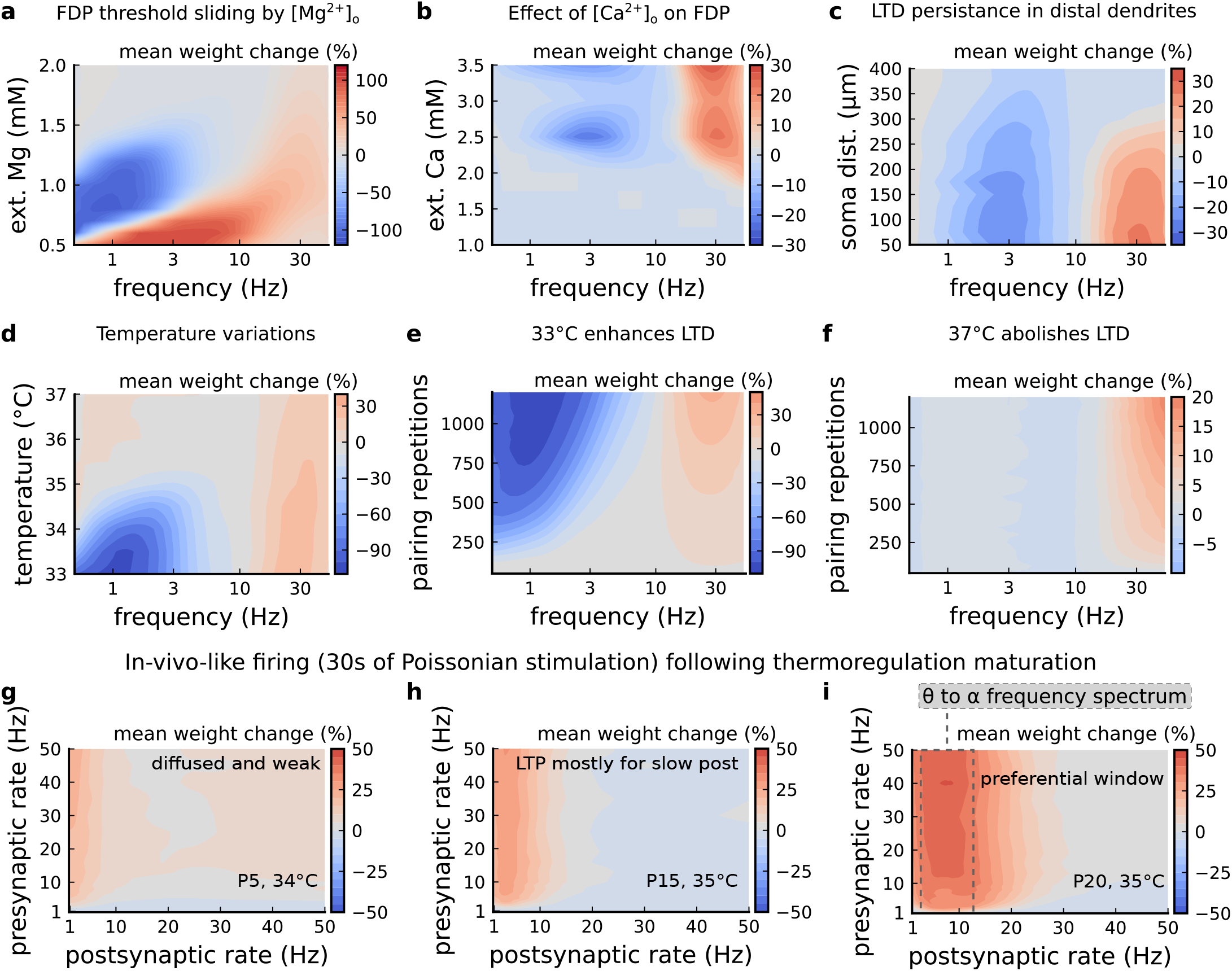
Varying experimental parameters in *Dudek and Bear (1992)* and Poisson spike train during development. **a**, Mean synaptic weight change for the FDP experiment varying the [Mg^2+^]_o_. Original [Mg^2+^]_o_ in ***Dudek and Bear (1992)*** is 1.5 mM (dashed grey line). **b**, Mean synaptic weight change for the FDP experiment varying the [Ca^2+^]_o_. Original [Ca^2+^]_o_ in ***Dudek and Bear (1992)*** is 2.5 mM (dashed grey line). **c**, Mean synaptic weight change for the FDP experiment varying the distant from the soma. Original distance in ***Dudek and Bear (1992)*** is 200 μm (dashed grey line). Changing the distance from the soma modifies how fast BaPs evoked by EPSP will attenuate. Note that LTD is prevalent for a spine situated far from the soma. **d**, Mean synaptic weight change for the FDP experiment varying the temperature. Original temperature in ***Dudek and Bear (1992)*** is 35°*C* (dashed grey line). **e**, Mean synaptic weight change for the FDP experiment varying the pairing repetitions at 33° C showing how LTD is enhanced. **f**, Mean synaptic weight change for the FDP experiment varying the pairing repetitions at 37°*C* showing how LTD is abolished. **g**, Mean synaptic weight change for pre and postsynaptic Poisson spike train during 30 s for P5 and 34°*C*. The panel shows that there is weak and diffused LTP. **h**, Mean synaptic weight change for pre and postsynaptic Poisson spike train during 30 s for P15 and 35°*C*. The panel shows that there is a start of LTP window forming for slow postsynaptic rates (<1 Hz). **i**, Mean synaptic weight change for pre and postsynaptic Poisson spike train during 30 s for P20 and 35°*C*. The panel shows that a window forms around 10 Hz postsynaptic rate similar to what is shown by ***Graupner et al. (2016)*** and in ***Figure 7h***.

**Figure 5 – Supplement 1.**
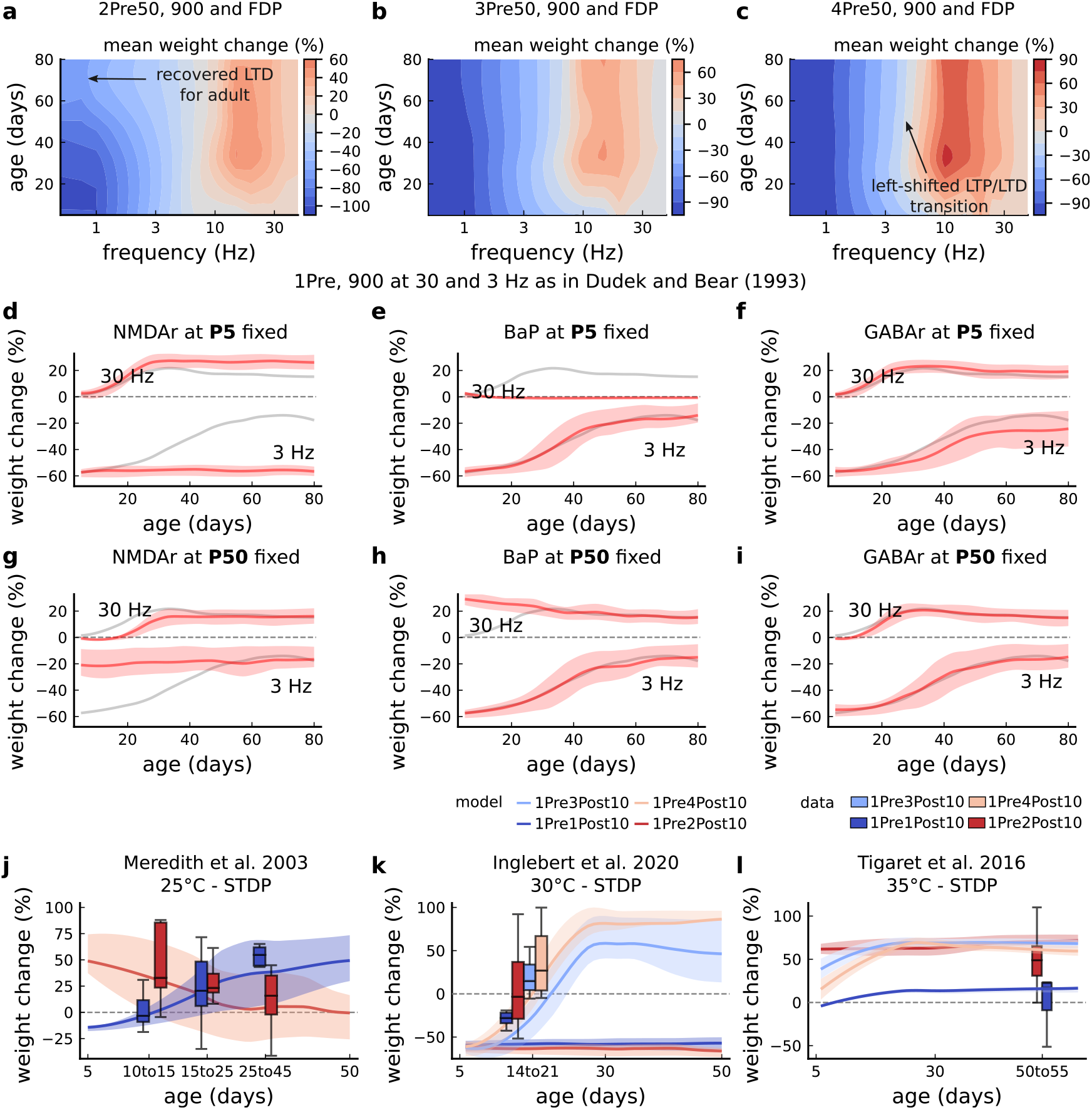
Duplets, triplets and quadruplets for FDP, perturbing developmental-mechanisms for LFS and HFS in *Dudek and Bear (1993)*, and age-related changes in STDP experiments (***Inglebert et al., 2020; Tigaret et al., 2016; Meredith et al., 2003***). **a**, Mean synaptic weight change (%) for the duplet-FDP (2Pre50) experiment varying age. The panel shows showing that not only LTD is enhanced but also LTP. **b**, Mean synaptic weight change (%) for the triplet-FDP (3Pre50) experiment varying age. The panel shows that LTD magnitude is enhanced for adult rats and the LTD-LTP transition is shifted leftward. **c**, Mean synaptic weight change (%) for the quadruplet-FDP (4Pre50) experiment varying age. The panel shows a further leftward shift on the LTD-LTP transition (compared to 3Pre50). **d**, Mean synaptic weight change (%) for the 1 Pre 900 at 30 and 3 Hz with ***Dudek and Bear (1993)***. Fixing NMDAr at P5 (more GluN2B than GluN2A) causes an increase of LTD and a slight increase of LTP for adult rats compared to baseline (grey solid line). **e**, Same experiment as panel **d** but fixing BaP maturation at P5 (higher BaP attenuation). LTP is abolished, but LTD is not affected. This is because AP induced by the EPSP attenuate too fast for 30 Hz and are thus not able to produce enough depolarization to activate NMDArs. **f**, Same experiment as in panel **d** but fixing GABAr maturation at P5 (excitatory GABAr) which only slighlty enhances LTD (3 Hz) for adult rats. **g**, Same experiment as panel **d** but fixing NMDAr at P50 (more GluN2A than GluN2B). LTD appears with decreased magnitude for young rats compared to baseline (grey solid line). **h**, Same experiment as panel **d** but fixing BaP maturation at P50 (less BaP attenuation). LTP is enhanced for young rats because the BaP pairing with the slow closing GluN2B produces more calcium influx. **i**, Same experiment as panel **d** but fixing GABAr maturation at P50 (inhibitory GABAr) which does not affect the FDP experiment. **j**, Mean synaptic weight change (%) for ***Meredith et al. (2003)*** single versus burst-STDP experiment for different ages. The data from Meredith (boxplots) were pooled by the age as shown in the x-axis. The solid line represents the mean, and the shaded ribbon the 2nd and 4th quantiles simulated by the model (same for panels **a-f**). **k**, Mean synaptic weight change (%) for ***Inglebert et al. (2020)*** STDP experiment in which the number of postsynaptic spikes increases. The x-axis marker from 14-21 indicates that only this interval was published without further specification. We use our model to estimate age related changes to ***Inglebert et al. (2020)*** protocols. Note that the model does not cover the 1Pre2Post10 properly (model predicts only outcomes near the first data quantile). Notice that single and burst STDP leads to LTD, meanwhile ***Meredith et al. (2003)*** lead to LTP or NC. **l**, Mean synaptic weight change (%) for ***Tigaret et al. (2016)*** STDP experiment which compares single versus burst STDP. The x-axis marker from 50-55 indicates that only a interval was published without further specification. We use our model to estimate age related changes to ***Tigaret et al. (2016)*** protocols. It is noticeable that each STDP experiment has a different development.

**Figure 6 – Supplement 1.**
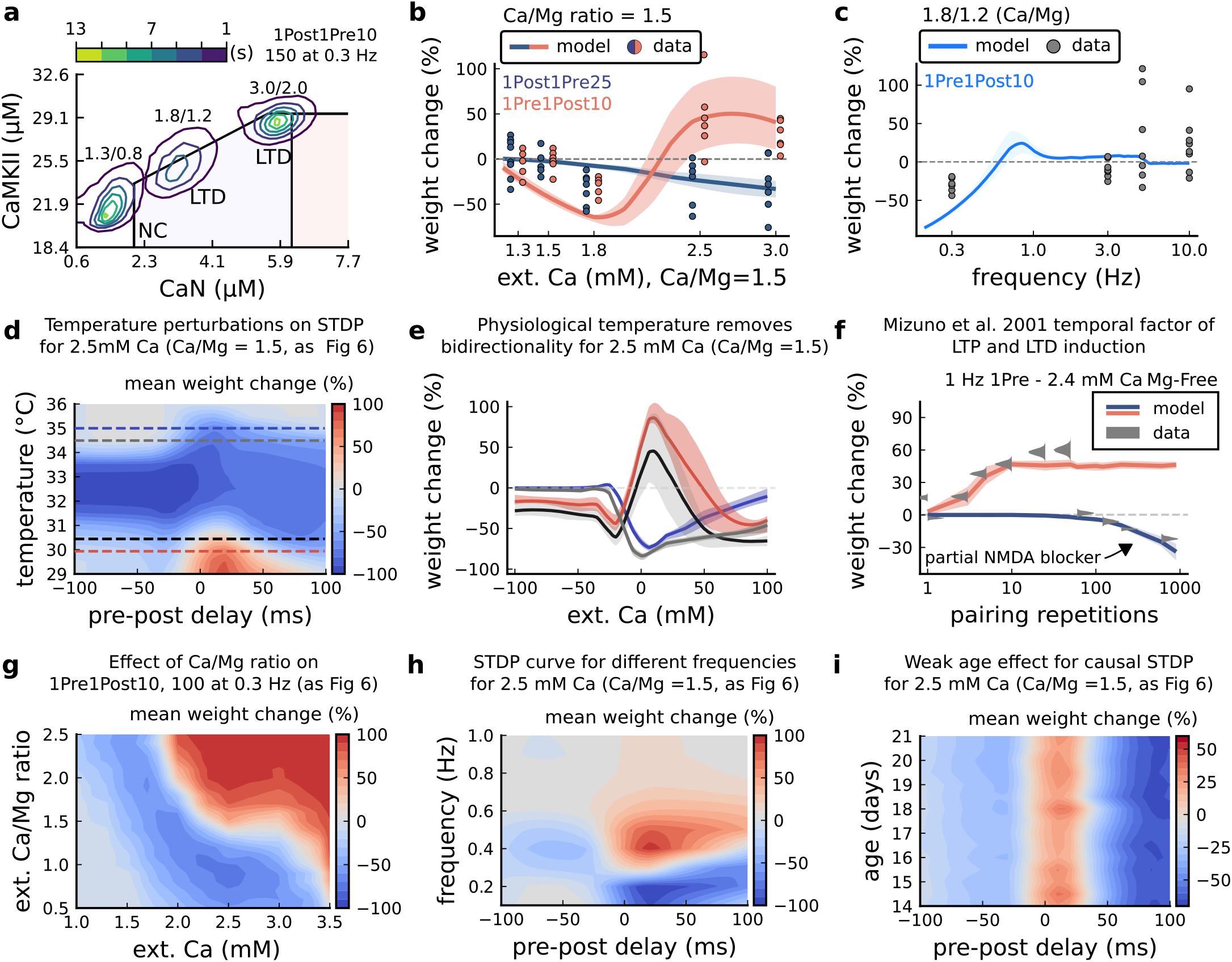
[Ca^2+^]_o_ and [Mg^2+^]_o_ related modifications for *Inglebert et al. (2020)* experiment. **a**, Mean time spent for anticausal pairing, 1Post1Pre10, at different Ca/Mg concentrations. The contour plots are associated with the ***Figure 6a-c.*** **b**, STDP and extracellular Ca/Mg. Synaptic weight change (%) for causal (1Pre1Post10,100 at 0.3 Hz) and anticausal (1Post1Pre10,150 at 0.3 Hz) pairings varying [Ca^2+^]_o_ from 1.0 to 3 mM (Ca/Mg ratio = 1.5). **c**, Varying frequency and extracellular Ca/Mg for the causal pairing 1Pre1Post10,100 at 0.3 Hz. Synaptic weight change (%) for a single causal pairing protocol varying frequency from 0.1 to 10 Hz. [Ca^2+^]_o_ was fixed at 1.8 mM (Ca/Mg ratio = 1.5). **d**, Mean synaptic weight change (%) for ***Inglebert et al. (2020)*** STDP experiment showing how temperature qualitatively modifies plasticity. The dashed lines are ploted in panel **b**. **e**, Mean synaptic weight change (%) showing effects 0.5°*C* from panel **a**. Black and grey solid lines represent the same color dashed lines in panel **a** (30 and 30.5°*C*). The bidirectional curves, black and grey lines in panel **a** (dashed) and panel **b** (solid), becoming full-LTD when temperature increases to 34.5 and 35°*C*, respectively yellow and purple lines in panel **a** (dashed) and panel **b** (solid). Further increase abolishes plasticity. **f**, Mean synaptic weight change (%) for ***Mizuno et al. (2001)*** experiment in Mg-Free ([Mg^2+^]_o_= 10^-3^mM for best fit) showing the different time requirements to induce LTP and LTD. For LTD, to simulate the NMDAr antagonist D-AP5 which causes a NMDAr partial blocking we reduced the NMDAr conductance by 97%. Note the similarity with ***Figure 3-Figure Supplement 5f***. **g**, Mean synaptic weight change (%) of ***Inglebert et al. (2020)*** STDP experiment changing [Ca^2+^]_o_ and Ca/Mg ratio. **h**, Mean synaptic weight change (%) of ***Inglebert et al. (2020)*** STDP experiment changing pre-post delay time and frequency. Note the similarity with ***Figure 3-Figure Supplement 5c***. **i**, Mean synaptic weight change (%) of ***Inglebert et al. (2020)*** STDP experiment changing pre-post delay time and age. Age has a weak effect on this experiment done at [Ca^2+^]_o_ = 2.5 mM.

**Figure 6 – Supplement 2.**
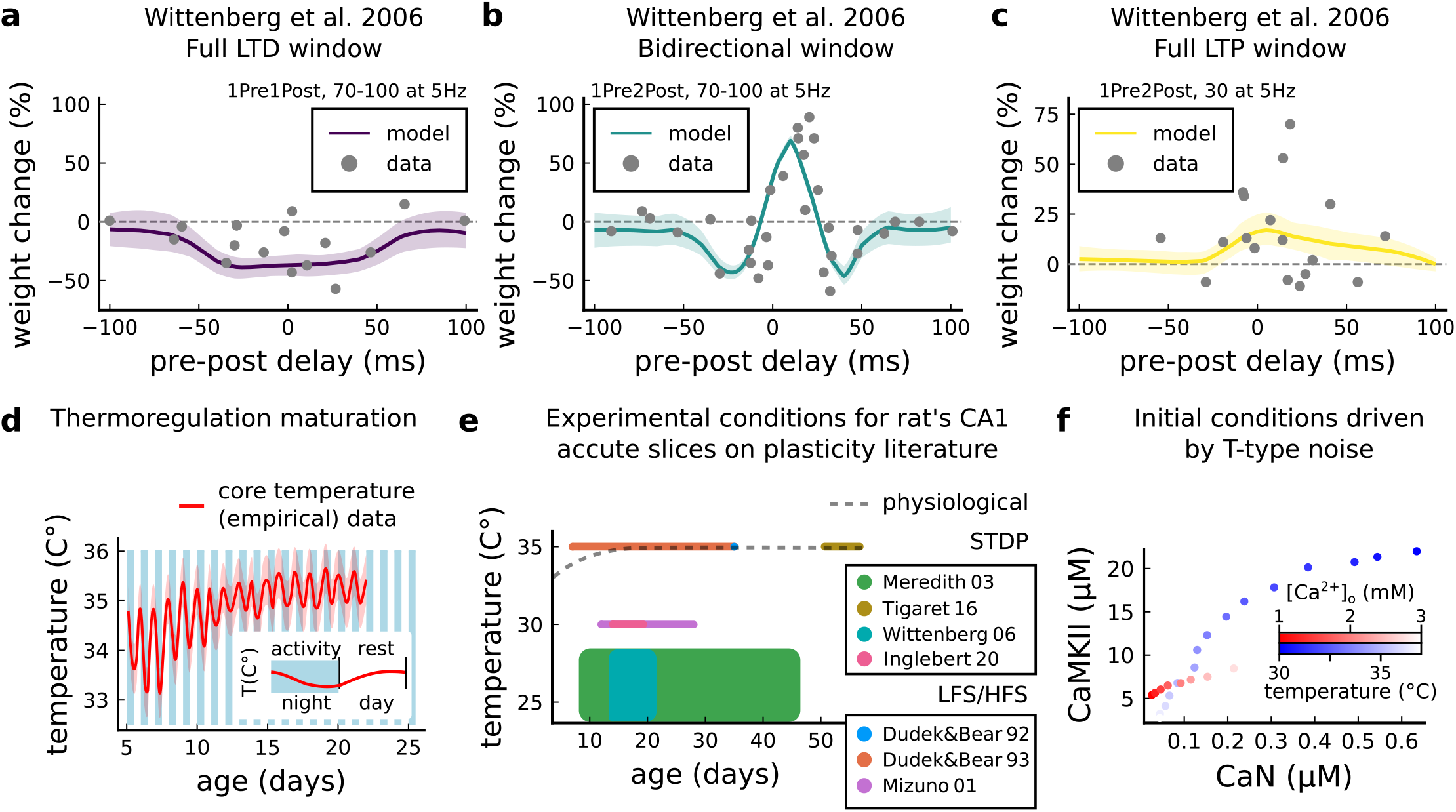
Temperature and age effects. **a**, Mean synaptic weight change (%) for ***Wittenberg and Wang (2006)*** STDP experiment for 1Pre1Post10, 70-100 at 5 Hz (see (***Table 1***)) showing a full LTD window. Our model also reproduces the data showing that when temperature is increased to 32 – 34°*C* LTD is abolished (data not shown). **b**, Mean synaptic weight change (%) for ***Wittenberg and Wang (2006)*** STDP experiment for 1Pre2Post10, 70-100 at 5 Hz (see (***Table 1***)) showing a bidirectional window. **c**, Mean synaptic weight change (%) for ***Wittenberg and Wang (2006)*** STDP experiment for 1Pre2Post10, 20-30 at 5 Hz (see (***Table 1***)) showing a bidirectional window. We noticed that for ***Wittenberg and Wang (2006)*** experiment, done in room temperature, the temperature sensitivity was higher than for other experiments. **d**, Core temperature varying with age representing the thermoregulation maturation. This function (not shown) was fitted using rat (***Wood et al., 2016***) and mouse data (***McCauley et al., 2020***) added by 1°*C* to compensate species differences (***Wood et al., 2016***). The blue and white bars represent the circadian rhythm as shown in ***McCauley et al. (2020)***. However, the “rest rhythm” for young rats (P5-14) may vary. **e**, Dotted grey line represents the averaged physiological temperature at different ages in the rat (estimated from mean value of panel **d**). For the papers the we fitted by the model, we depict the range of temperature and age used. Note that only few experiments were performed at near physiological conditions. **f**, Initial conditions for CaN-CaMKII resting concentration for different [Ca^2+^]_o_ and temperature values. When [Ca^2+^]_o_ is changed, temperature is fixed at 35°*C*, while when temperature is changed, [Ca^2+^]_o_ is fixed at 2 mM.

**Table 1 – Supplement 1.**
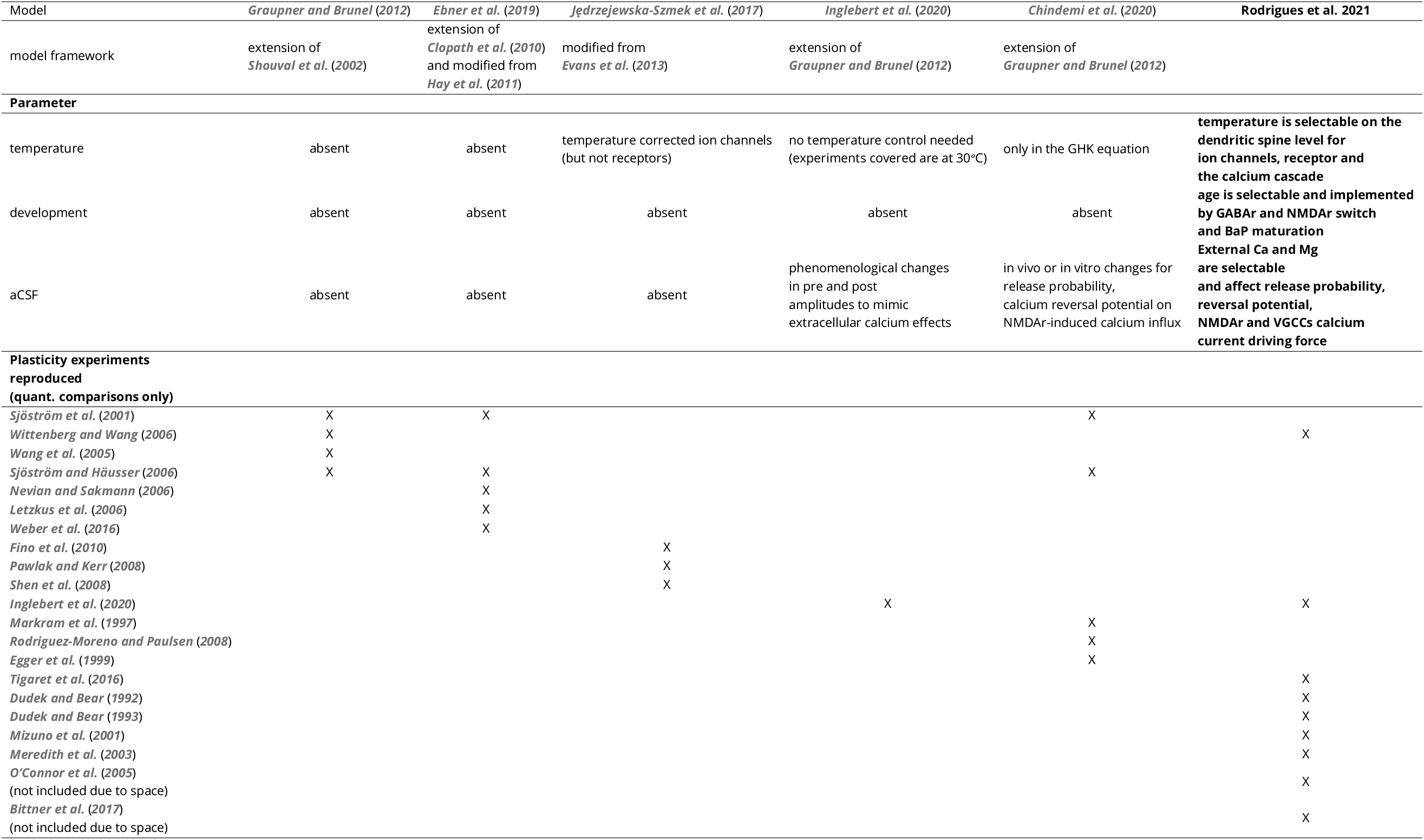
Comparison of recent computational models for plasticity highlighting the experimental conditions implemented and the experiments in the hippocampus and cortex they reproduce. See ***Table 1-Table Supplement 2*** for additional details on experimental conditions of experimental works.

**Table 1 – Supplement 2.** Comparison of the experimental conditions for the different reproduced datasets in ***Table 1-Table Supplement 1*** covering experiments from neocortex, hippocampus and striatum

| Experimental work | Age (days) | [Ca<sup>2+</sup>]<sub>o</sub> (Mm) | [Mg<sup>2+</sup>]<sub>o</sub> (Mm) | Temperature (°C) |
| --- | --- | --- | --- | --- |
| *Sjöström et al. (2001)* | 12-21 | 2.5 | 1 | 32-34 |
| *Wittenberg and Wang (2006)* | 14-21 | 2 | 1 | 24-30 or 30-34 |
| *Wang et al. (2005)* | embryonic day 17-18 | 3 | 2 | room |
| *Sjöström and Häusser (2006)* | 14-21 | 2 | 1 | 32-35 |
| *Nevian and Sakmann (2006)* | 13-15 | 2 | 1 | 32-35 |
| *Letzkus et al. (2006)* | 21-42 | 2 | 1 | 34-35 |
| *Weber et al. (2016)* | 49-77 | 1.25 | 1.3 or 0.1 | 32-35 |
| *Fino et al. (2010)* | 15-21 | 2 | 1 | 34 |
| *Pawlak and Kerr (2008)* | 19-22 | 2.5 | 2 | 31-33 |
| *Shen et al. (2008)* | 19-26 | 2 | 1 | room |
| *Inglebert et al. (2020)* | 14-20 | 1.3-3.0 | Ca/1.5 | 30 |
| *Markram et al. (1997)* | 14-16 | 2 | 1 | 32-34 |
| *Rodriguez-Moreno and Paulsen (2008)* | 9-14 | 2 | 2 | room |
| *Egger et al. (1999)* | 12-14 | 2 | 1 | 34-36 |
| *Tigaret et al. (2016)* | 50-55 | 2.5 | 1.3 | 35 |
| *Dudek and Bear (1992)* | 35 | 2.5 | 1.5 | 35 |
| *Dudek and Bear (1993)* | 7-35 | 2.5 | 1.5 | 35 |
| *Mizuno et al. (2001)* | 12-28 | 2.4 | Mg-Free (most experiments) | 30 |
| *Meredith et al. (2003)* | 9-45 | 2 | 2 | 24-28 |
| *O'Connor et al. (2005)* | 14-21 | 2 | 1 | 27.5-32 |
| *Bittner et al. (2017)* | 42-63 | 2 | 1 | 35 |

## Notes

### Competing Interest Statement

The authors have declared no competing interest.

### Summary of Updates

Grammar revised, this version brings additional support figures/tables, modification of intro/discussion mainly.

